# Propulsive cell entry diverts pathogens from immune degradation by remodeling the phagocytic synapse

**DOI:** 10.1101/2023.04.25.538287

**Authors:** Zihan Zhang, Thomas K. Gaetjens, Yanqi Yu, D. Paul Mallory, Steven M. Abel, Yan Yu

## Abstract

Phagocytosis is a critical immune function for infection control and tissue homeostasis. This process is typically described as non-moving pathogens being internalized and degraded in phagolysosomes. For pathogens that evade immune degradation, the prevailing view is that virulence factors that biochemically disrupt the biogenesis of phagoslysosomes are required. In contrast, here we report that physical forces exerted by pathogens during cell entry divert them away from the canonical phagolysosomal degradation pathway, and this altered intracellular fate is determined at the time of phagocytic synapse formation. We used the eukaryotic parasite *Toxoplasma gondii* as a model because live *Toxoplasma* uses gliding motility to actively invade into host cells. To differentiate the effect of physical forces from that of virulence factors in phagocytosis, we developed a strategy that used magnetic forces to induce propulsive entry of inactivated *Toxoplasma* into macrophage cells. Experiments and computer simulations collectively reveal that large propulsive forces suppress productive activation of receptors by hindering their spatial segregation from phosphatases at the phagocytic synapse. Consequently, the inactivated parasites, instead of being degraded in phagolysosomes, are engulfed into vacuoles that fail to mature into degradative units, following an intracellular pathway strikingly similar to that of the live motile parasite. Using opsonized beads, we further confirmed that this mechanism is general, not specific to the parasite used. These results reveal previously unknown aspects of immune evasion by demonstrating how physical forces exerted during active cell entry, independent of virulence factors, can help pathogens circumvent phagolysosomal degradation.

## Introduction

Phagocytosis is a vital process by which innate immune cells ingest and degrade invading pathogens. The effective clearance of pathogens requires a complex, tightly interwoven cascade of biochemical transformations and physical activities. First, the pathogen is recognized by receptors on the immune cell. The function of these receptors involves more than just binding to its specific ligand. At the contact site between pathogen and immune cell, receptor-ligand complexes cluster and organize spatially to form a microscale pattern of proteins called the phagocytic synapse (1-3). Signaling from activated receptors in this synapse triggers remodeling of the actin network in the cell to facilitate engulfment of the pathogen by the cell membrane protrusion. Once trapped inside a membrane-bounded compartment, namely the phagosome, the pathogen is then destroyed. Nascent phagosomes transform into compartments that degrade and destroy the pathogen through a sequence of maturation steps, involving the increasing acidification of their lumen (4), fusion with endosomes and lysosomes to develop into phagolysosomes (5, 6), and acquisition of hydrolytic enzymes for content digestion (7-11).

Pathogens, including bacteria, fungi, and parasites, have evolved diverse strategies to evade surveillance and degradation by immune cells. Many pathogens, such as *Bacillus subtilis*, avoid receptor-mediated recognition and internalization. They do this either by injecting into host cells virulence factors that inactivate signaling pathways responsible for initiating phagocytosis (12-15), or by modifying their own cell envelopes to avoid triggering phagocytosis (16-19). Other pathogens, such as *M. tuberculosis* (20, 21) and *S. pyogenes (22, 23)*, are phagocytosed by the host immune cells, but survive and replicate intracellularly by circumventing the phagolysosomal degradation pathway. Their known mechanisms are to use virulence effectors to prevent the acidification of phagosomes or to suppress the fusion of phagosomes with lysosomes (23-26). Despite the diversity of immune-evasion mechanisms used by pathogens, the prevalent view is that all of them work biochemically, by using virulence factors that inhibit one or more steps in phagocytosis. However, emerging evidence raises the possibility that the physical process of pathogen motility may also be critical to the pathogen’s invasion of its host.

Motility is known to be essential to the infection cycle of several pathogens, including parasites like *Trypanosoma cruzi* and bacteria like *Legionella pneumophila* (27, 28). For example, a mutant strain of *Legionella* lacking the flagella needed for motility was found to be much less efficient at invading cells than the wild-type strain, leading to the finding that the flagellum of *Legionella* facilitates its encounter with the host cell (29). Trypomastigotes of the parasite *Trypanosoma cruzi* were shown to modify their movements depending on the type of mammalian cell being invaded (30). Apicomplexans are a diverse group of obligate intracellular protozoan parasites that cause severe human diseases like malaria (31, 32). They are also known to use gliding motility to facilitate their invasion of host cells (33-35). However, the role of pathogen motility in infection has been thought to be confined only to the initial establishment of the infection. Studies suggested that the motility of pathogens might promote their binding to host cells (29, 36) and their active penetration of the host cell’s membrane (37-39). Little is known about how the motility of pathogens might impact their fate after they actively penetrate the cell.

Towards addressing this broad question, we investigate here how the physical forces, as expected from motile pathogens, might impact their intracellular trafficking. Our study uses the protozoan parasite *Toxoplasma gondii* as the pathogen model. *Toxoplasma* is an opportunistic pathogen that causes congenital disorders and life-threatening complications in immunocompromised individuals (40-42). Like the many pathogens highlighted above, its gliding motility, driven by its cytoskeleton and myosin motors, was found to be critical for its infection (37-39). The propulsive motility helps the parasite actively penetrate the host cell membrane (43-45). After internalization, *Toxoplasma* resides in parasitophorous vacuoles (PVs). This unique type of membrane compartment does not acidify (46) or fuse with lysosomes (47, 48). The lack of degradative functions within these vacuoles allows the parasite to survive and replicate inside. One proposed explanation for the biogenesis of the non-degradative vacuoles is that the parasite, during active invasion, secrets effector proteins that block the residing vacuoles from fusing with lysosomes (47, 49-51). However, as in the case of other motile pathogens, it is unclear if and how the propulsive force from active invasion might contribute to the ability of the parasite to evade degradation.

A major challenge in addressing this question is differentiating the effects of physical forces from the biochemical effects of virulence systems, which are often coordinately regulated with the motility apparatus (27). To study the role of pathogen motility in infection, one widely used approach is to generate motility mutants that are viable in culture. This approach, however, has drawbacks. Motility mutants can still have residual motility; the same mutation that causes a motility deficit might also cause loss of the functions of other proteins that are involved in virulence systems. Such drawbacks might be the reason for the discrepancies between previous studies (36, 52).

In this study, we devised a method to pinpoint the effect of propulsive cell entry by using magnetic manipulation to apply propulsion during the internalization of pathogens. Using *Toxoplasma* as the model pathogen, we tethered magnetic nanoparticles to the surfaces of inactivated parasites to render them magnetically responsive. As we magnetically propelled the parasite to enter macrophage cells, we simultaneously monitored the intracellular fate of the pathogen-residing vacuoles. If the inactivated *Toxoplasma* can be induced to actively penetrate the macrophage cells, just as the live parasite does, can this physical force alone alter their phagocytic fate? We demonstrate that inactivated *Toxoplasma* that is passively internalized is degraded in phagolysosomes, but the magnetic propulsive force applied during cell entry results in the parasite being encapsulated in vacuoles that fail to mature into degradative units. With induced propulsive cell entry, the inactivated *Toxoplasma* enters an intracellular pathway strikingly similar to that of the live parasite. Using opsonized synthetic particles instead of inactivated *Toxoplasma*, we further showed that this effect of physical forces is general regardless of the type of phagocytic targets. Experiments and computer simulations revealed a possible mechanistic explanation for this effect. It appears that large propulsive forces suppress the productive activation of receptors by hindering their spatial segregation from phosphatases at the phagocytic synapse. Consequently, instead of being degraded in phagolysosomes, cargos are engulfed into vacuoles that fail to proceed through the usual sequence of maturation events, from the recruitment of endocytic markers, to acidification, fusion with lysosomes, and proteolytic digestion.

## Results

### Live *Toxoplasma* resides in vacuoles lacking degradative functions, but heat-killed *Toxoplasma* is degraded in phagolysosomes

Previous studies have shown that *Toxoplasma*, upon penetrating the host cell membrane, enters parasitophorous vacuoles that fail to acidify or fuse with endocytic vesicles (46-48). To first confirm this, we added live *Toxoplasma* to RAW 264.7 macrophage cells and then examined the recruitment of endocytic and lysosomal markers on the *Toxoplasma*-containing vacuoles (5, 7, 53-55). These markers, including small GTPase Rab5, Rab7, and lysosome-associated membrane proteins (LAMP)1, are sequentially recruited to phagosome membranes to regulate their fusion with endosomes and lysosomes (Fig. 1*A*) (56-59). It is these fusion events that deliver degradative enzymes into phagosomes that destroy the pathogen (7-10). Therefore, the accumulation of those markers is a key indicator of proper maturation and degradative function of phagosomes (60, 61). In our experiments, we used fluorescence microscopy to image the recruitment of Rab5-GFP, Rab7-RFP, and LAMP1-RFP on vacuoles containing live *Toxoplasma (*expressing tubulin GFP for visualization). We found negligible recruitment of any of these three markers on most *Toxoplasma*-containing vacuoles at 7.5 min, 15 min, and 40 min after *Toxoplasma* was added to cells (Fig. 1 *B*-*D*). Only a small fraction of the vacuoles was positive for Rab5, Rab7, and LAMP1. This was likely because some of the *Toxoplasma* tachyzoites were dead during the isolation and purification process. We then prepared heat-killed *Toxoplasma* and added them to the cells. Different from live *Toxoplasma*, the heat-killed ones were engulfed into phagosomes that showed accumulation of all three markers (Fig. 1 *B*-*D*). Rab5 accumulation occurred shortly after internalization (7.5 min). Following Rab5 dissociation, Rab7 and LAMP1 were then recruited to the phagosomes and remained on the membrane for the imaging duration of 40 min (Fig. 1 *B-D*). These results confirm that live *Toxoplasma* resided in intracellular vacuoles that lack degradative functions, whereas heat-killed *Toxoplasma* was passively internalized via the canonical phagolysosomal degradative pathway. Having confirmed this result, we then asked: if the heat-killed parasite can be made to exert propulsion force during cell entry, like the live parasite does, can the physical force alter its intracellular trafficking fate?

**Fig. 1.**
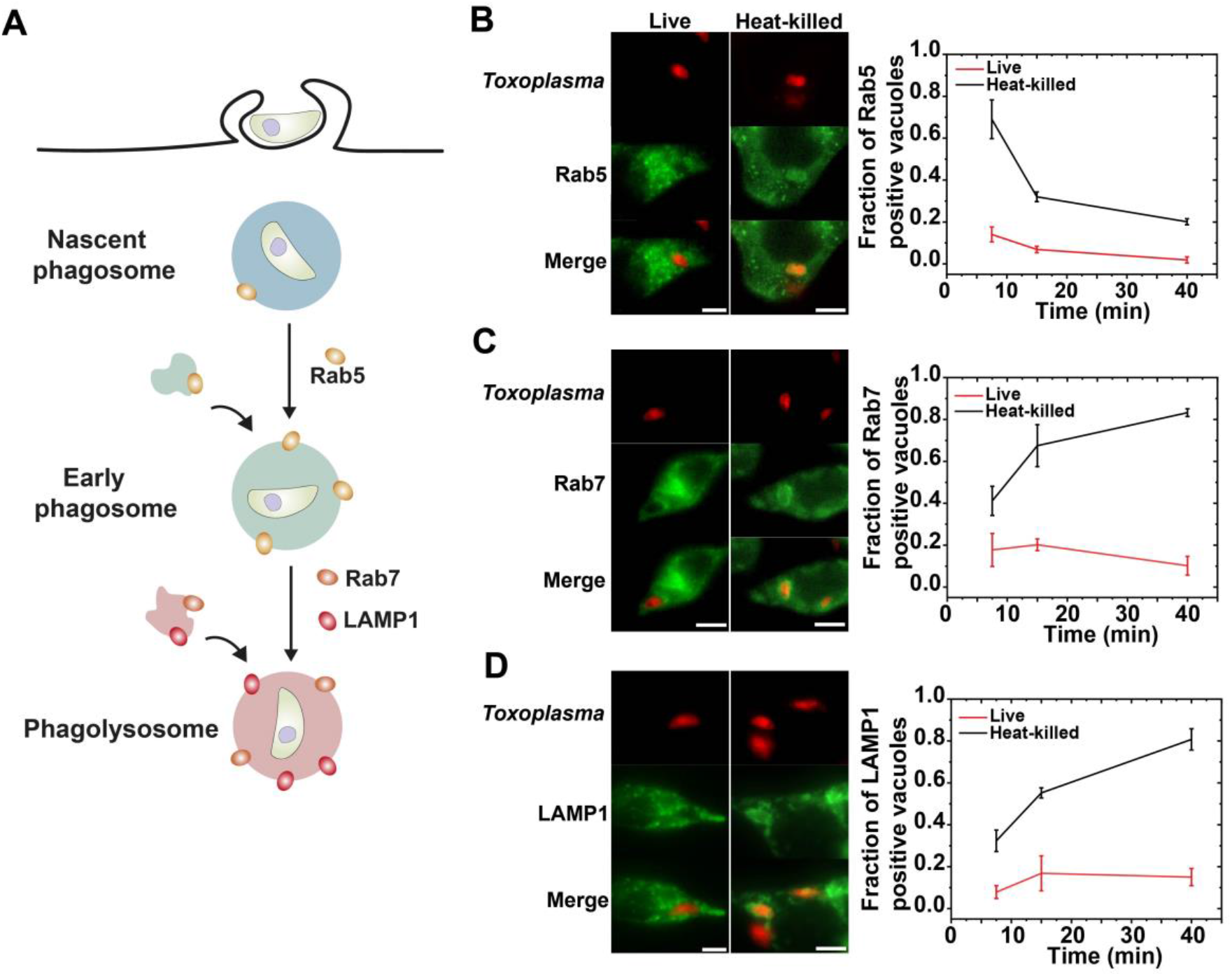
Live and heat-killed *Toxoplasma* entered distinct intracellular trafficking pathways in RAW 264.7 macrophage cells. (*A*) Schematic illustration showing Rab5, Rab7, and LAMP1 recruitment during phagosome maturation along the canonical phagolysosomal degradation pathway. (*B*-*D*) Fluorescence images and quantitative analysis of the assembly of endocytic markers Rab5, Rab7, and LAMP1 on phagosomes encapsulating heat-killed *Toxoplasma* versus vacuoles encapsulating live *Toxoplasma* in macrophage cells. Data were obtained at 7.5 min, 15 min and 40 min after parasites were added to cells. Error bars in line plots represent standard deviations from 3 independent experiments for each maker tested. Scale bars in fluorescence images, 5 µm.

### Design and characterization of magnetically responsive heat-killed *Toxoplasma*

We included the propulsive cell entry of heat-killed *Toxoplasma* by using a magnetic tweezers setup (SI Appendix, Fig. S1) and then simultaneously imaged their intracellular traffickiing in RAW 264.7 cells (Fig. 2*A*). Briefly, we attached 200 nm magnetic nanoparticles onto the surface of heat-killed *Toxoplasma* using carbodiimide crosslinking, thereby rendering them magnetically responsive (Fig. 2*A*). Using fluorescent magnetic nanoparticles, we found that ≈ 46% of the heat-killed *Toxoplasma* tachyzoites were conjugated with magnetic nanoparticles (SI Appendix, Fig. S2 *A* and *B*). We referred to those conjugated with magnetic nanoparticles as mag-*Toxoplasma*, for simplicity. Most mag-*Toxoplasma* had 1-5 magnetic nanoparticles, as shown using scanning electron microscopy (SEM) micrographs (Fig. 2*B*). We selected 200 nm magnetic particles as they provide robust magnetic forces without interfering with the major surface antigen (SAG-1) of *Toxoplasma* or hindering their host cell entry (Fig. S2*A*). One single 200 nm magnetic nanoparticle can exert forces of 5-10 pN (Fig. S2*C*), and the magnetic force decays with increased distance between the nanoparticle and the solenoid tip, consistent with theoretical predictions (62-65). We cannot establish a similar force-distance plot for heat-killed *Toxoplasma*, because the movement of these irregularly shaped and deformable parasites does not follow the Stokes’ equation. Nevertheless, we confirmed that the mag-*Toxoplasma* are magnetically responsive, by observing their faster directional movements towards the solenoid tip (SI Appendix, Fig. S3). We estimated the magnetic force on a single mag-*Toxoplasma* to be around 10-50 pN, depending on the number of conjugated nanoparticles per parasite. This range of force is comparable to forces generated from the gliding motion of living *Toxoplasma* (66) and *M. pneumoniae* on surfaces (67).

**Fig. 2.**
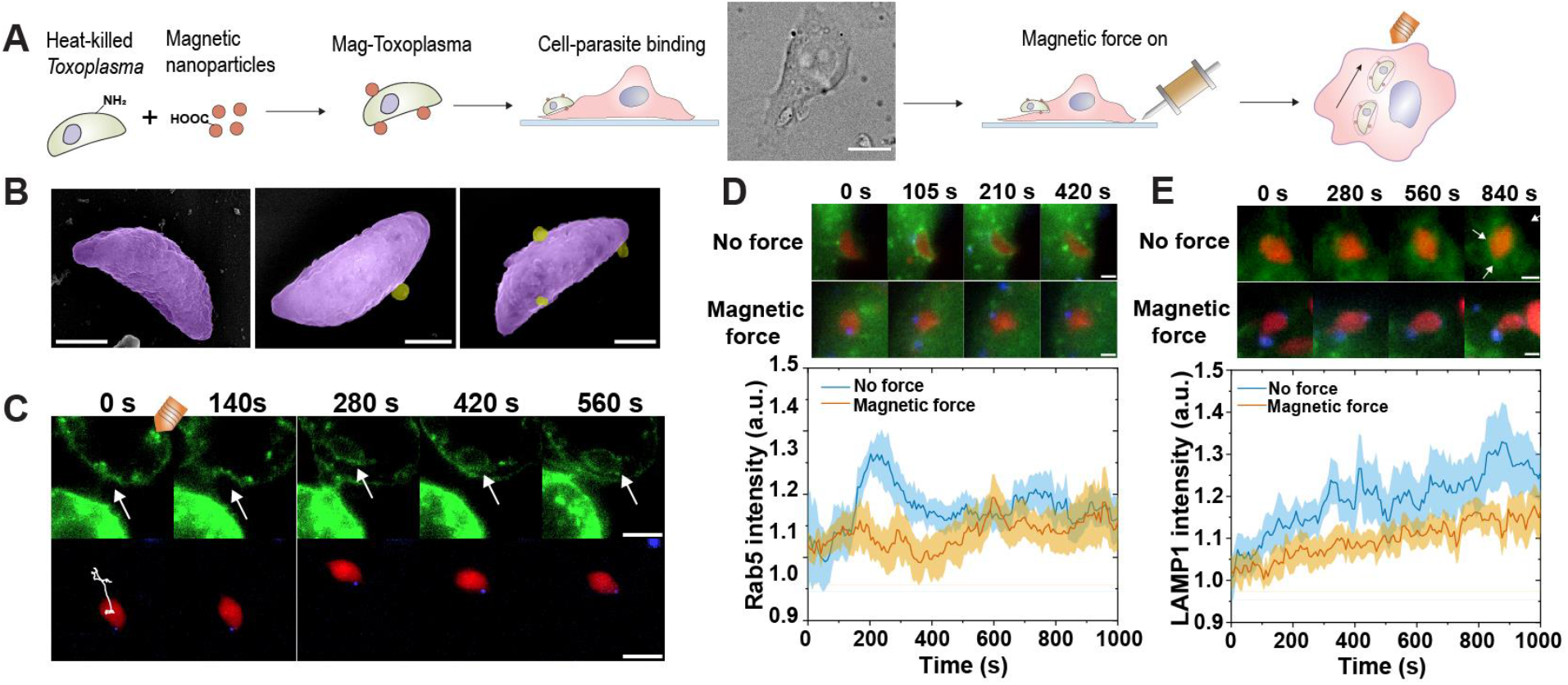
Propulsion force during cell entry causes heat-killed *Toxoplasma* to enter vacuoles lacking endocytic markers. (*A*) Schematic illustration of the experiment design. RH strain *Toxoplasma gondii* tachyzoites were killed by heat, then conjugated with 200 nm magnetic nanoparticles via EDC conjugation. The resulting mag-*Toxoplasma* was each tethered with about 1-5 magnetic nanoparticles. The magnetic tweezers tip was positioned on the opposite side of the cell body from the parasite of interest. Magnetic pulling force was applied on mag-*Toxoplasma* after they were bound to RAW 264.7 cell membrane and kept on throughout the imaging measurement. Scale bar in bright field image, 10 µm. (*B*) SEM images of mag-*Toxoplasma*. Images are assigned with pseudo colors, *Toxoplasma* in magenta and magnetic nanoparticles in yellow. Scale bars, 2 µm. (*C*) Fluorescence images showing the cell entry of a mag-*Toxoplasma* under magnetic pulling force. In the sequential images, the mag-*Toxoplasma* was labeled with Alexa 568 (shown in red), magnetic nanoparticles were labeled with CF640R (shown in blue), and macrophage cell membrane were labeled with CF488A conjugated Cholera Toxin Subunit B (CTB) (shown in green). The white line in the first image shows the moving trajectory of a mag-*Toxoplasma*. Scale bars, 5 µm. (*D, E*) Fluorescence images show the assembly of Rab5 (D) and LAMP1 (E) on phagosomes encapsulating heat-killed *Toxoplasma* with or without magnetic pulling force. *Toxoplasma* is shown in red, magnetic nanoparticles in blue, and Rab5-GFP and LAMP1-GFP in green. Scale bars, 2 µm. Line plots show the average fluorescence intensity of Rab5-GFP and LAMP1-GFP on phagosomes with or without magnetic pulling force. Line curves are averages from 13 (no force) and 15 (with magnetic force) single phagosomes. Shaded areas represent the standard error of the mean. Scale bars, 2 µm.

In our experiments, we first allowed the mag-*Toxoplasma* to bind to the RAW 264.7 macrophage cells in a culture dish, and then used the magnetic tweezers to pull the membrane-bound mag-*Toxoplasma* towards the cell center by placing the tweezers across the cell body from the parasite of interest (Fig. 2 *A* and *C*). This magnetic pulling was intended to mimic the propulsion force exerted by live *Toxoplasma* during cell entry. We maintained the magnetic force after the internalization of mag-*Toxoplasma* throughout the imaging process, unless otherwise specified. Under the magnetic pulling force, the mag-*Toxoplasma* initially adhered to the cell membrane and then moved rapidly and directionally towards the tip of the magnetic tweezers upon internalization (Fig. 2*C*). Using gangliosides GM1 marker CTB-CF488 to label the cell plasma membrane, we confirmed that the mag-*Toxoplasma* was internalized into membrane vacuoles marked by CTB-CF488 fluorescence (Fig. 2*C*). This observation also confirmed that the magnetic force we applied was appropriate to allow membrane engulfment of the heat-killed parasite without rupturing the plasma membrane (68, 69).

### Heat-killed *Toxoplasma* with induced propulsive cell entry enters vacuoles that do not fuse with endosomes or lysosomes

We next sought to investigate whether the vacuoles containing mag-*Toxoplasma* after propulsive cell entry undergo the same membrane transformation as needed to mature into phagolysosomes. We have shown that when heat-killed *Toxoplasma* was internalized without magnetic force, the endocytic markers Rab5 and Rab7, and lysosomal marker LAMP1, were present on phagosomes as expected for typical phagosome maturation (Fig. 1 *B-D*). In live cell imaging, we observed that Rab5-GFP was recruited transiently to the phagosome membrane containing heat-killed *Toxoplasma*, reaching an intensity peak shortly after the complete formation of phagosomes and then gradually decreasing afterwards (Fig. 2*D*, N =13; single phagosome data in SI Appendix, Fig. S4*A*). LAMP1-GFP was continuously recruited to the phagosome (Fig. 2*E*, N =11; single phagosome data in SI Appendix, Fig. S5*A*). In contrast, when magnetic force was applied during internalization, the vacuoles containing mag-*Toxoplasma* showed significantly impaired recruitment of Rab5 and LAMP1, compared to that for live *Toxoplasma*. The level of Rab5-GFP remained consistently low on the vacuoles, in contrast to the typical process of Rab 5 recruitment and association (Fig. 2*D*, N =15; single phagosome data in SI Appendix, Fig. S4*B*). Further, the level of LAMP1-GFP was also significantly lower than that without magnetic force (Fig. 2*E*, N =10; single phagosome data in SI Appendix, Fig. S5*B*). The results suggest that the inactivated *Toxoplasma* with propulsive cell entry was internalized into vacuoles that fail to recruit the endocytic markers, an essential step in the maturation of nascent phagosomes to phagolysosomes. Next, we sought to determine whether this observation is specific to *Toxoplasma* or it generalizes to other types of internalized particles. For simplicity, from here on, we will refer to the vacuoles formed after propulsive cell entry as phagosomes, even though they may not function as normal phagosomes.

To test the generality of our findings, we performed experiments using 1 µm magnetic beads opsonized with immunoglobulin G (IgG) to trigger Fc gamma receptor (Fc*γ*R)-mediated phagocytosis (Fig. 3*A*) (70, 71). As in the mag-*Toxoplasma* experiments, we applied magnetic force to the beads after they bound to cells and maintained the force until the end of imaging. The propulsive force exerted by a 1 µm magnetic bead was estimated to be about 25 pN at the beginning of magnetic manipulation and 30 pN at the end (Fig. 3*B* and SI Appendix, Fig. S6). We confirmed the internalization of the beads by using a trypan blue quenching assay that quenches fluorescence of non-internalized beads and by directly imaging the phagosome membrane using a plasma membrane marker PM-RFP (SI Appendix, Fig. S7). Without magnetic force, Rab5-GFP was recruited and then dissociated from the bead-containing phagosomes, whereas Rab7-GFP and LAMP1-GFP were recruited gradually (Fig. 3*C*, N = 15 and SI Appendix, Fig. S8*A*; Fig. 3*D*, N = 8 and SI Appendix, Fig. S9*A*; Fig. 3*E*, N = 9 and SI Appendix, Fig. S10*A*). However, when magnetic force was applied during internalization, there was no significant recruitment of these markers to the phagosomes. This is consistent with the results from mag-*Toxoplasma* experiments described above (Fig. 3*C*, N = 15 and SI Appendix, Fig. S8*B*; Fig. 3*D*, N = 10 and SI Appendix, Fig. S9*B*; Fig. 3*E*, N = 11 and SI Appendix, Fig. S10*B*).

**Fig. 3.**
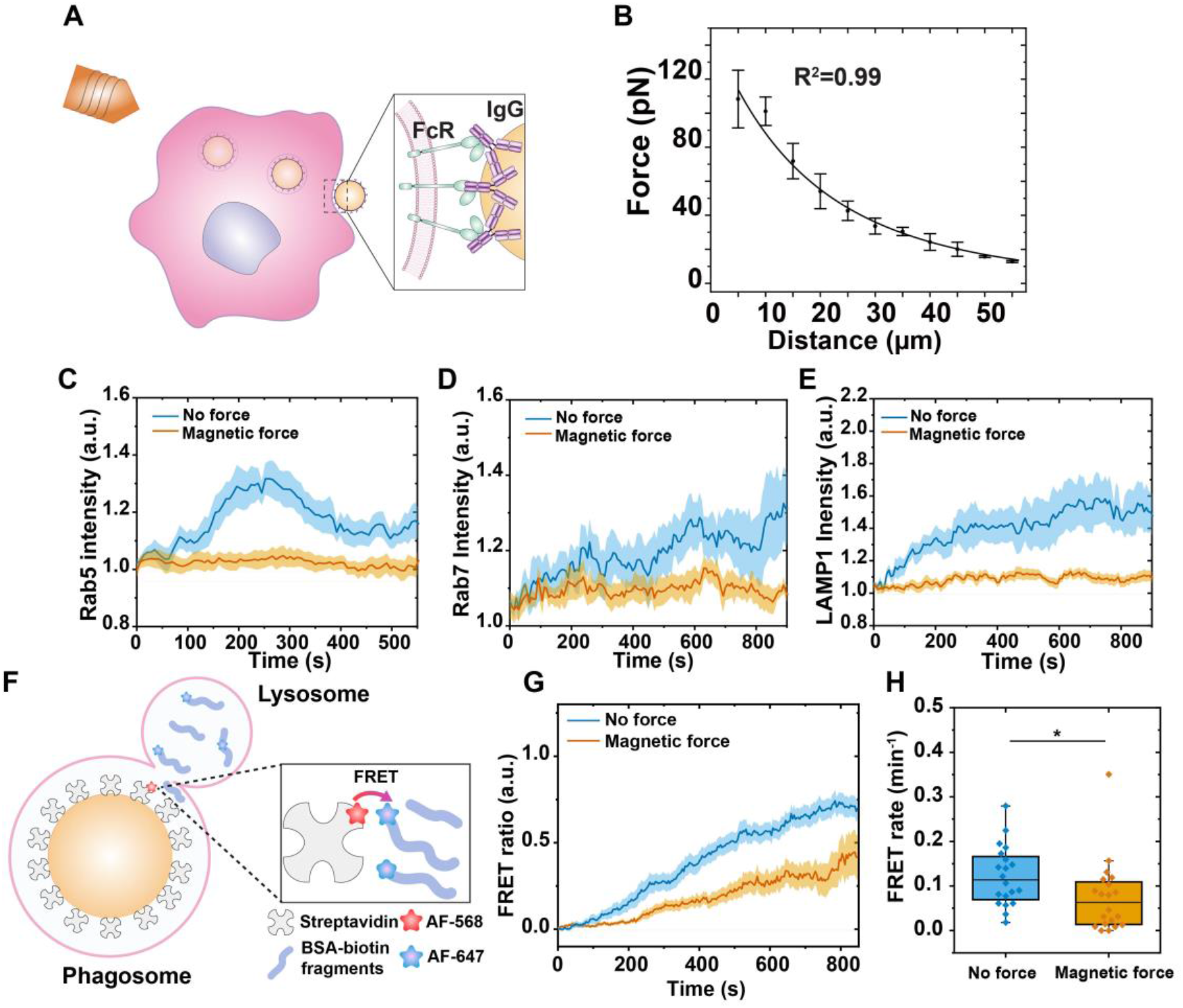
Propulsion force during cell entry causes opsonized beads to enter vacuoles that do not fuse with endocytic compartments. (*A*) Schematic illustration of the experimental design. The 1 µm magnetic beads were physically absorbed with IgG. Magnetic pulling force was applied on the beads after they bound to the plasma membrane of RAW 264.7 cells. (*B*) Calibration plot showing the magnetic force exerted on a single bead as a function of distance between the bead and the tip of the magnetic tweezers solenoid. Error bars are standard deviation from 5 samples. (*C*-*E*) Line plots showing the average time-dependent intensity of assembly of Rab5-GFP (no force, N = 15; with magnetic force, N = 15), Rab7-GFP (no force, N = 8; with magnetic force, N = 10), and LAMP1-GFP (no force, N = 9; with magnetic force, N = 11) on bead-containing phagosomes. Shaded areas represent standard error of the mean. (*F*) Schematic illustration showing the working principle of phagosome-lysosome fusion measurements based on Förster resonance energy transfer (FRET). The 1 μm FRET-MagSensors were functionalized with streptavidin-AF568 (FRET-donor) and IgG. Biotin-BSA-AF647 were incubated with RAW 264.7 cells overnight and chased for 2 h into lysosome compartments. (*G*) Line plots show the average FRET ratio as a function of time with or without magnetic pulling force as indicated. The line curves are averages from 20 single phagosomes for each experimental condition. Shaded areas represent standard error of the mean. (*H*) Box graph shows the average FRET rate under different experiment conditions as indicated. The average FRET rate is 0.12 ± 0.07 a.u. without magnetic force (N = 20) and 0.07 ± 0.08 a.u. with magnetic pulling force (N = 20). Each box plot indicates the mean (horizontal line) and the interquartile range from 25% to 75% of the corresponding data set. Statistical significance is highlighted by p-values (student’s t-test) as follows: *p < 0.05.

Because the recruitment of Rab5, Rab7, and LAMP1 are prerequisites for phagosomes to fuse with lysosomes and acquire digestive enzymes, we next directly quantified how propulsive entry into the cell affects the phagosome fusion with lysosomes using a Förster Resonance Energy Transfer (FRET) based assay (Fig. 3*F*). Details of this assay has been described in our recent study (72). In the FRET fusion experiments, we first biotinylated 1 µm magnetic beads and conjugated them with streptavidin labeled with donor fluorophore Alexa 568. We referred to those magnetic beads as FRET-magSensors. We also loaded lysosomes in cells with biotinylated bovine serum albumin (BSA) that was labeled with acceptor fluorophore Alexa 647. Upon phagosome fusion with lysosomes, streptavidin-Alexa 568 on the surface of the FRET-magSensors mixes with fragmented BSA-biotin-Alexa 647 in lysosomes, resulting in FRET signals (SI Appendix, Fig. S11*A*). Since each phagosome can fuse simultaneously with multiple lysosomes, the fusion events collectively resulted in a gradual decrease in donor emission (Alexa 568, ex/em: 561/586 nm) and concurrently an increase in acceptor emission in donor excitation (Alexa 647, ex/em: 561/680 nm) (SI Appendix, Fig. S11*C*). We quantified the FRET ratio of individual phagosomes using the following equation:

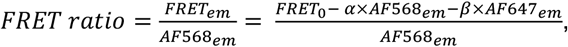

In which *FRET*_*em*_ is acceptor emission upon donor excitation (ex/em 561/680 nm) after correction for spectral bleed-through, and *AF*568_*em*_ is donor emission (ex/em 561/586 nm). *FRET*_0_ represents the acceptor emission under donor excitation (ex/em 561/680 nm) before correction. The coefficient *α* corrects the bleed-through contribution from the donor (Alexa568) due to the overlap of the donor emission in the FRET channel. The coefficient *β* corrects the non-FRET bleed-through from the acceptor (Alexa647) due to the excitation of the acceptor at the donor excitation wavelength. Correction factors *α* and *β* were determined for each independent experiment (details in SI Appendix, Materials and Methods).

Without magnetic force, the FRET ratio of single phagosomes during maturation followed a sigmoidal relationship with time (Fig. 3*G*, N = 20; SI Appendix, Fig. S11*E*), which is consistent with our previous finding (72). By comparison, phagosomes formed under magnetic force exhibited a significantly slower increase in FRET ratio (Fig. 3*G*, N = 20; SI Appendix, Fig. S11 *B*-*F*). We further quantified the kinetic rate of phagosome-lysosome fusion (referred to as FRET rate) by calculating the slope of the FRET ratio increase, based on the sigmoidal-Boltzmann fitting of single phagosome data (SI Appendix, Fig. S11*E*). We found that phagosomes fused with lysosomes at an average rate of 0.07 ± 0.08 a.u. under magnetic force (Fig. 3*H*, N = 20), in contrast to an average rate of 0.12 ± 0.07 a.u. without magnetic force (Fig. 3*H*, N = 20).

To confirm whether the reduced fusion of phagosomes with lysosomes was caused by propulsive cell entry, we performed control experiments in which we pulled FRET-magSensors during internalization but then turned off the magnetic force after bead internalization (Fig. S12*A*). This did not result in any noticeable recovery of phagosome-lysosome fusion (Fig. S12*B*). The results altogether indicate that propulsive cell entry, induced by magnetic force, disrupts the fusion of parasite-residing vacuoles with endosomes and lysosomes. This is the consequence of the impaired recruitment of endocytic and lysosomal markers including Rab5, Rab7, and LAMP1. This is a general finding regardless of the nature of the phagocytic cargo, as dysfunctional membrane remodeling caused by the exertion of propulsive force during internalization was consistently observed using both heat-killed *Toxoplasma* and IgG-coated beads as cargos.

### Heat-killed *Toxoplasma* with induced propulsive cell entry resides in vacuoles lacking acidification and degradative function

During normal maturation, phagosomes fuse with lysosomes to acquire proton pump vacuolar-type H+-ATPase (V-ATPase) (73, 74) and hydrolytic enzymes (7-10). The proton pumps maintain an acidic lumen environment, which prohibits the proliferation of pathogens and is required for the activation of proteolysis and the generation of reactive oxygen species for content degradation (8, 75-77). With the observation that both mag-*Toxoplasma* and beads under magnetic force were internalized into phagosomes that failed to fuse with lysosomes, we then examined if those phagosomes lost their capacity to acidify and degrade. To monitor the acidification of the phagosomes, we conjugated a pH sensitive dye, pHrodo Red, on the surface of the mag-*Toxoplasma* via amine-NHS ester reaction (Fig. 4*A*). All parasites were labeled with pHrodo Red (Fig. S2 *A* and *D*), and the dye labeling did not block the major surface antigen 1 (SAG-1) from binding to its antibody (SI Appendix, Fig. S2*E*). The fluorescence intensity of pHrodo Red (*I*_*pHrodo*_) of single mag-*Toxoplasma* increased linearly as the buffer pH decreased (average plot in Fig. 4*B*; single mag-*Toxoplasma* plots in Fig. S13).

**Fig. 4.**
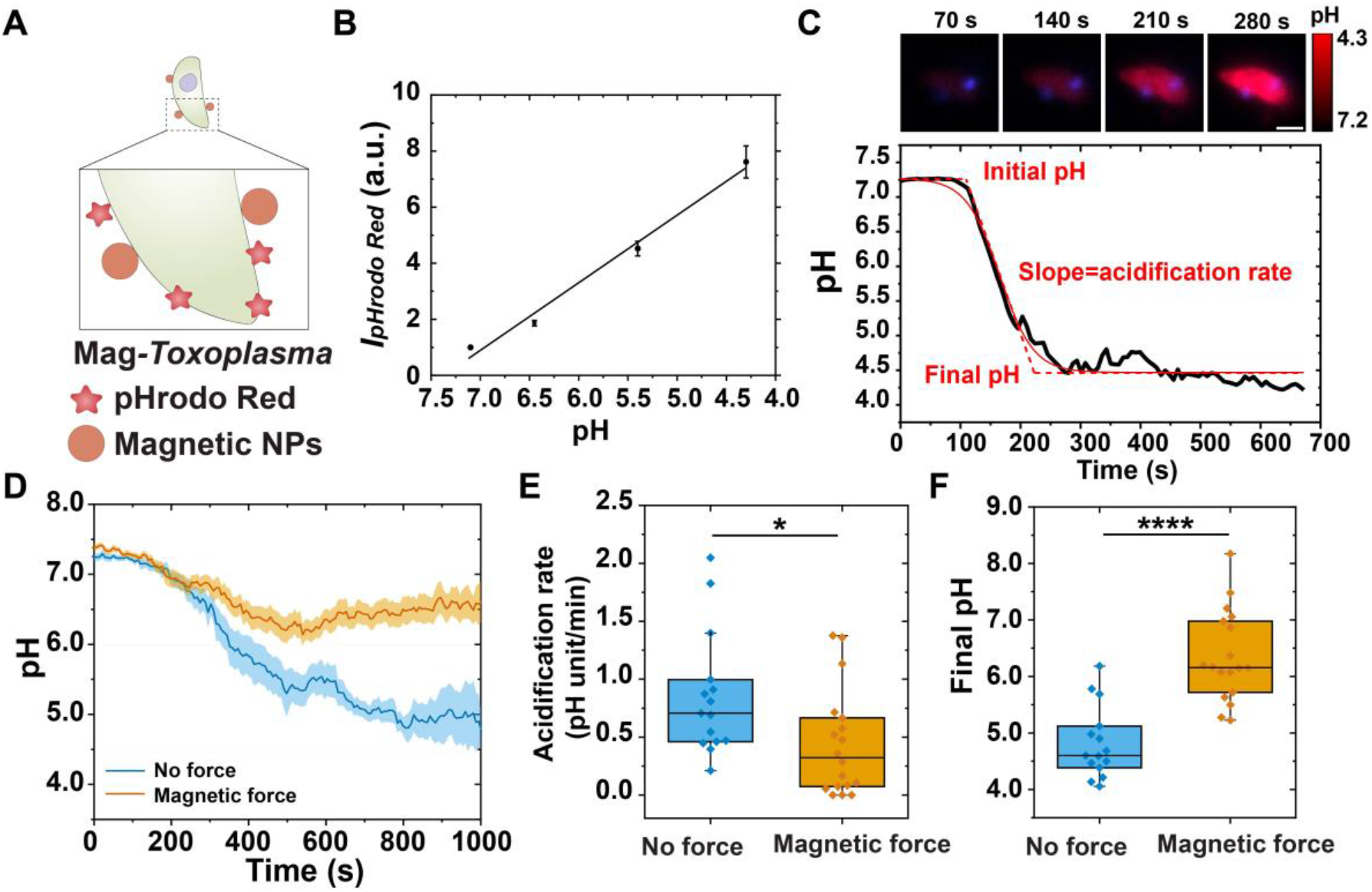
Propulsion force during cell entry causes heat-killed *Toxoplasma* to enter vacuoles lacking acidification and degradative function. (*A*) Schematic illustration showing the surface functionalization of pH sensitive mag-*Toxoplasma*. Heat-killed *Toxoplasma* was first conjugated with magnetic nanoparticles and then the pH indicator pHrodo Red. (*B*) Extracellular pH calibration plot showing the fluorescence intensity change of pHrodo Red labeled mag-*Toxoplasma* in Ringer’s imaging buffer at different pH values. The line plot is an average of 14 single mag-*Toxoplasma* and error bars represent standard deviations. (*C*) Fluorescence images and line curves showing the pH change of a representative mag-*Toxoplasma*-containing phagosome as a function of time without magnetic manipulation. Mag-*Toxoplasma* is shown in red and CF-640R labeled magnetic nanoparticles in blue. Scale bar, 2 µm. The pH-time line plot is fitted with sigmoidal-Boltzmann function (red solid line) to determine the initial pH, final pH, and acidification rate. The red dotted line indicates the determination of slope. Scale bar, 2 µm. (*E*) Box graph showing the average acidification rate of individual mag-*Toxoplasma*-containing vacuoles with or without magnetic pulling force. The average acidification rate is 0.85 ± 0.53 pH unit/min without magnetic manipulation (N = 15) and 0.44 ± 0.46 pH unit/min with magnetic pulling (N = 18). (*F*) Box graph showing the average phagosome final pH under different experiment conditions as indicated. The average final pH is 4.8 ± 0.6 without magnetic manipulation (N = 15) and 6.3 ± 0.8 with magnetic pulling (N = 18). In (*E* and *F*), each box plot indicates the mean (horizontal line) and the interquartile range from 25% to 75% of the corresponding data set. Statistical significance is highlighted by p-values (student’s t-test) as follows: ****p <0.0001 and *p < 0.05.

Without magnetic force, phagosomes containing mag-*Toxoplasma* acidified rapidly within a few minutes after internalization and the acidity eventually reached a plateau around pH 4.5 (Fig. 4*C*; more example plots in Fig. S14 *A*-*C*). This three-stage acidification profile is consistent with our previous observations of phagosome acidification using synthetic beads (72, 76). We obtained the initial and final pH, as well as the rate of acidification, of single phagosomes by fitting their pH vs. time plots with a sigmoidal-Boltzmann function (Fig. 4*C*):

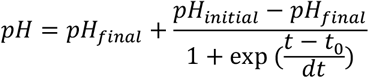

We found that the application of magnetic propulsive force on mag-*Toxoplasma* during internalization significantly impaired phagosome acidification (Fig. 4 *D*-*F*, N =15; single vacuole data in SI Appendix, Fig. S14 *D*-*F*). With propulsive cell entry, acidification of the phagosomes slowed down, at an average rate of 0.44 ± 0.46 pH unit/min (N =18) compared to 0.85 ± 0.53 pH unit/min for normal phagosomes (N = 15) (Fig. 4*E*). Meanwhile, the final pH of the phagosomes (6.3 ± 0.8, N =16) was higher than that of normal phagosomes without forces applied (4.8 ± 0.6, N = 15) (Fig. 4*F*). Further, the dysfunctional acidification of phagosomes did not recover to normal when the magnetic force was turned off (SI Appendix, Fig. S15), which confirmed that phagosome acidification was perturbed by the propulsive force applied during internalization.

To test the generality of this observation, we performed the experiments using pH-responsive magnetic beads (referred to as pH-magSensors). We first biotinylated 1 µm magnetic beads, then conjugated them with a mixture of pHrodo Red-labeled streptavidin and CF640R-labeled streptavidin, and finally coated the beads with IgG via physical adsorption (Fig. 5*A*), following a protocol we reported previously (72). The pHrodo Red dye is the pH indicator, whereas the CF640R dye is the reference because it is pH insensitive and photostable. The ratiometric fluorescence between pHrodo Red and CF640R (*I*_*pHrodo*_/*I*_*ref*_) increases linearly with decreased pH in aqueous buffers (SI Appendix, Fig. S16*A*) and inside phagosomes in cells (SI Appendix, Fig. S16*B*). As with the mag-*Toxoplasma* results, magnetic force applied during the internalization of beads caused a higher final pH and the slower acidification of phagosomes (Fig. 5*B-D*; single phagosome plots in SI Appendix, Fig. S17). Phagosomes without magnetic force reached a final pH of 4.7 ± 0.3 (N = 33) at an average rate of 0.48 ± 0.34 pH unit/min (N = 33). In contrast, phagosomes formed under magnetic force reached a significantly less acidic pH of 6.3 ± 0.8 (N = 26) at an average rate of 0.18 ± 0.20 pH unit/min (N = 26) (Fig. 5 *B-D*).

**Fig. 5.**
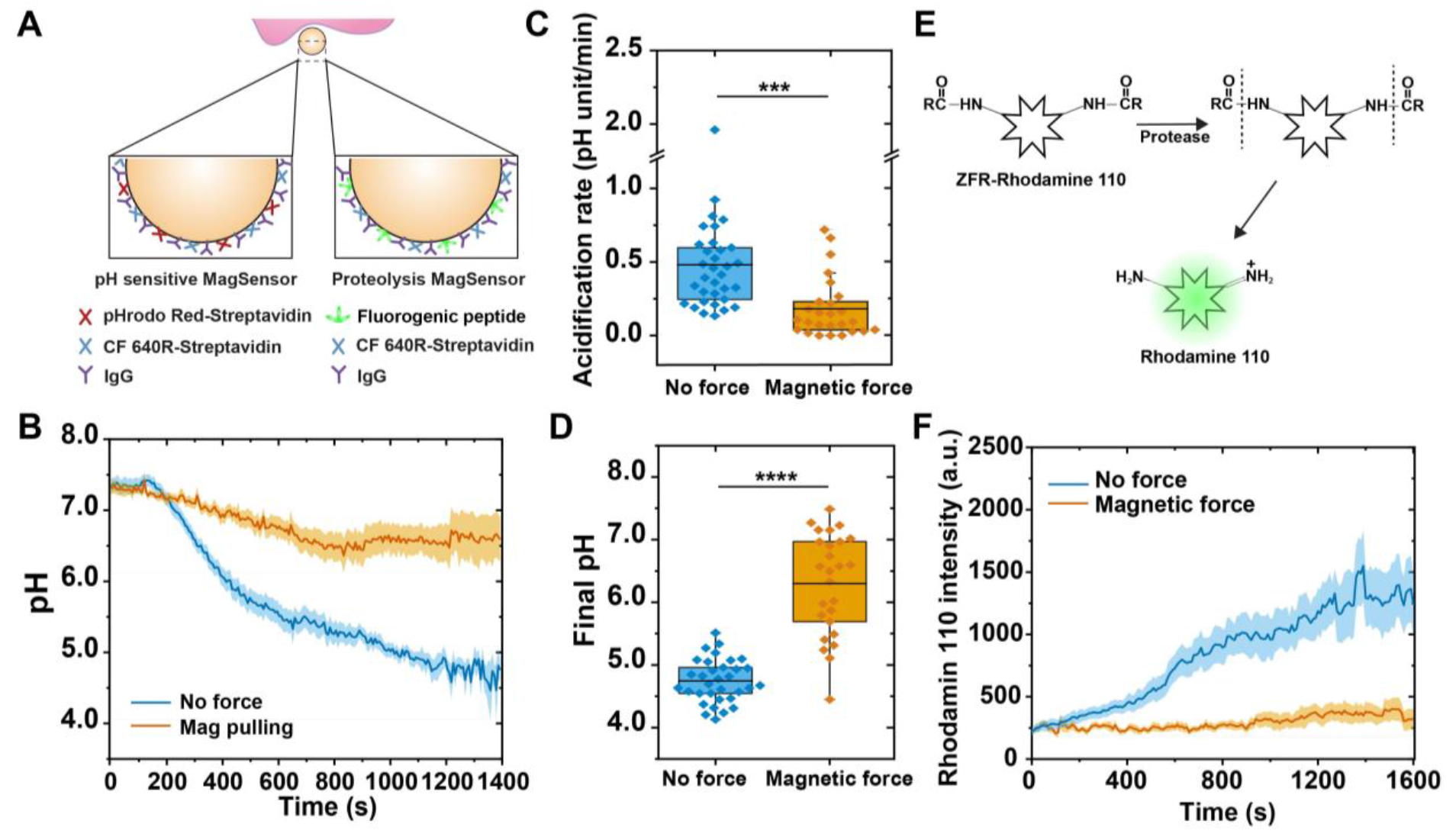
Propulsion force during cell entry causes opsonized beads to enter vacuoles lacking acidification and degradative function. (*A*) Schematic illustration showing the design of pH sensitive MagSensor (pH-MagSensor) and proteolysis sensitive MagSensor (proteolysis-MagSensor). The 1 μm pH-MagSensors were functionalized with streptavidin-pHrodo Red (pH indicator) and streptavidin-CF 640 (reference dye), and physically adsorbed with IgG. The 1 µm proteolysis-MagSensors were functionalized with a fluorogenic peptide ZFR-R110 (Rhodamine 110, bis-(N-CBZ-L-phenylalanyl-Larginine amide)) and IgG. (*B*) Line plots showing the average phagosome pH as a function of time with no force applied and with magnetic pulling during cell entry. The line curves are averages from 25 (no force) and 20 (with magnetic force) individual phagosomes. Shaded areas represent standard error of the mean. (*C*) Statistic result showing the average acidification rate in different experiments as indicated. The rates are: 0.48 ± 0.34 pH unit/min (without magnetic manipulation, N = 33) and 0.18 ± 0.20 pH unit/min (with magnetic pulling, N = 26) (*D*) Statistic result showing the average final pH of bead-containing vacuoles with and without magnetic force. The average final pH is 4.7 ± 0.3 (without magnetic force, N = 33) and 6.3 ± 0.8 (with magnetic force, N = 26). In both scattered plots, each box plot indicates the mean (horizontal line) and the interquartile range from 25% to 75% of the corresponding data set. Statistical significance is highlighted by p-values (student’s t-test) as follows: ****p < 0.0001, ***p < 0.001. (*E*) Schematic showing the working principle of fluorogenic peptide Z-FR-R110 (Rhodamine 110, bis-(N-CBZ-L-phenylalanyl-L-arginine amide) dihydrochloride) for phagosome proteolytic activity measurements. The Z-FR-R110 peptide releases the Rhodamine 110 dye once peptide bonds are cleaved by the cysteine proteases. (*F*) Line plots show the average rhodamine 110 intensity (no force, N = 9; with magnetic force, N = 6) indicating proteolysis within bead-containing vacuoles as a function of time with or without magnetic pulling. Shaded areas represent standard error of the mean.

We next quantified proteolytic activity of phagosomes, another indicator of degradative function. In these experiments, we conjugated biotinylated magnetic beads (1 µm) with CF640R-labeled streptavidin and then coated them with IgG and fluorogenic peptide Z-FR-R110 (Rhodamine 110, bis-(N-CBZ-L-phenylalanyl-Larginine amide) dihydrochloride) (Fig. 5*A*). The Z-FR-R110 peptide is a substrate for cysteine proteases, including cathepsins B and L, which are key digestive enzymes for cargo degradation in phagosomes (78, 79). Upon cleavage of the peptide bonds, the Rhodamine 110 dye “caged” in this fluorogenic peptide is released, resulting in strong fluorescence emission (Fig. 5*E*). We referred to these magnetic beads as proteolysis-magSensors. Without magnetic force, the Rhodamine 110 intensity remained low for a few minutes after internalization, but increased rapidly afterwards (Fig. 5*F*, N = 9; SI Appendix, Fig. S18*A*). This late start of proteolysis is because activation of the proteases requires acidic pH in the phagosome lumen (76, 80). However, when magnetic force was applied, proteolytic activity in phagosomes was significantly impaired, as evidenced by the low level of Rhodamine 110 intensity (Fig. 5*F*, N = 6 and SI Appendix, Fig. S18*B*). The results from phagosome acidification and proteolysis both demonstrate that the degradative function of phagosomes was compromised by the propulsive force during internalization. The force applied disrupted the cascade of activities in phagosome maturation. Without the proper recruitment of endocytic machineries like Rab5, phagosomes cannot fuse with endosomes and lysosomes, and thus cannot acquire either the proton pump V-ATPase for lumen acidification or proteolytic enzymes for pathogen degradation (74).

### Propulsive force alters protein spatial organization at the phagocytic synapse

After elucidating how propulsive cell entry perturbs phagosome maturation, we next sought to identify the cause of this disruptive effect. When microbes are phagocytosed, ligand-bound receptors and signaling proteins reorganize spatially to form a phagocytic synapse at the contact area between the microbe and the host cell (1, 3, 81, 82). A key characteristic of this phagocytic synapse is that inhibitory phosphatases like CD45 are spatially excluded from the pathogen-host contact area where activated receptors, such as Dectin-1 and Fc*γ*Rs, are concentrated (1-3). Based on studies of T cells and B cells (1, 83-85), microscale protein segregation occurring in the phagocytic synapse is likely to be a critical mechanism for sustaining productive signaling from phagocytic receptors, an event preceding the proper assembly of the nascent phagosome. Therefore, we hypothesized that propulsive force exerted during cell entry could affect the formation of the phagocytic synapse formation and thereby trigger a domino effect of disruption to the degradative function of phagosomes that we observed. Because it is unclear which receptors on macrophage cells are responsible for the phagocytosis of *Toxoplasma*, we examined the spatial re-organization of the tyrosine phosphatase CD45. We labeled CD45 on the live cell membrane using a low concentration of Alexa647 anti-CD45 antibody, following a previously reported procedure (2). During the internalization of heat-killed *Toxoplasma* without magnetic force, CD45 was excluded from the parasite-cell contact area, as indicated by its lower fluorescence intensity at the contact site compared to that in the membrane outside the contact site (Fig. 6 *A* and *C*; SI Appendix Fig. S19*A*). However, when magnetic force was applied, no exclusion of CD45 was observed at the parasite-cell contact area (Fig. 6 *B* and *D*; SI Appendix, Fig. S19*B*). Interestingly, the magnetic force had no effect on the distribution of the cell membrane glycosphingolipid GM1, which was labeled with CTB-CF488 (SI Appendix Fig. S20 *A*-*B*). Because CD45 has a large, rigid extracellular domain, this suggests that the propulsion force during internalization selectively hindered the re-organization of bulky proteins like CD45 during phagocytic synapse formation (schematic illustration in Fig. 6*E*).

**Fig. 6.**
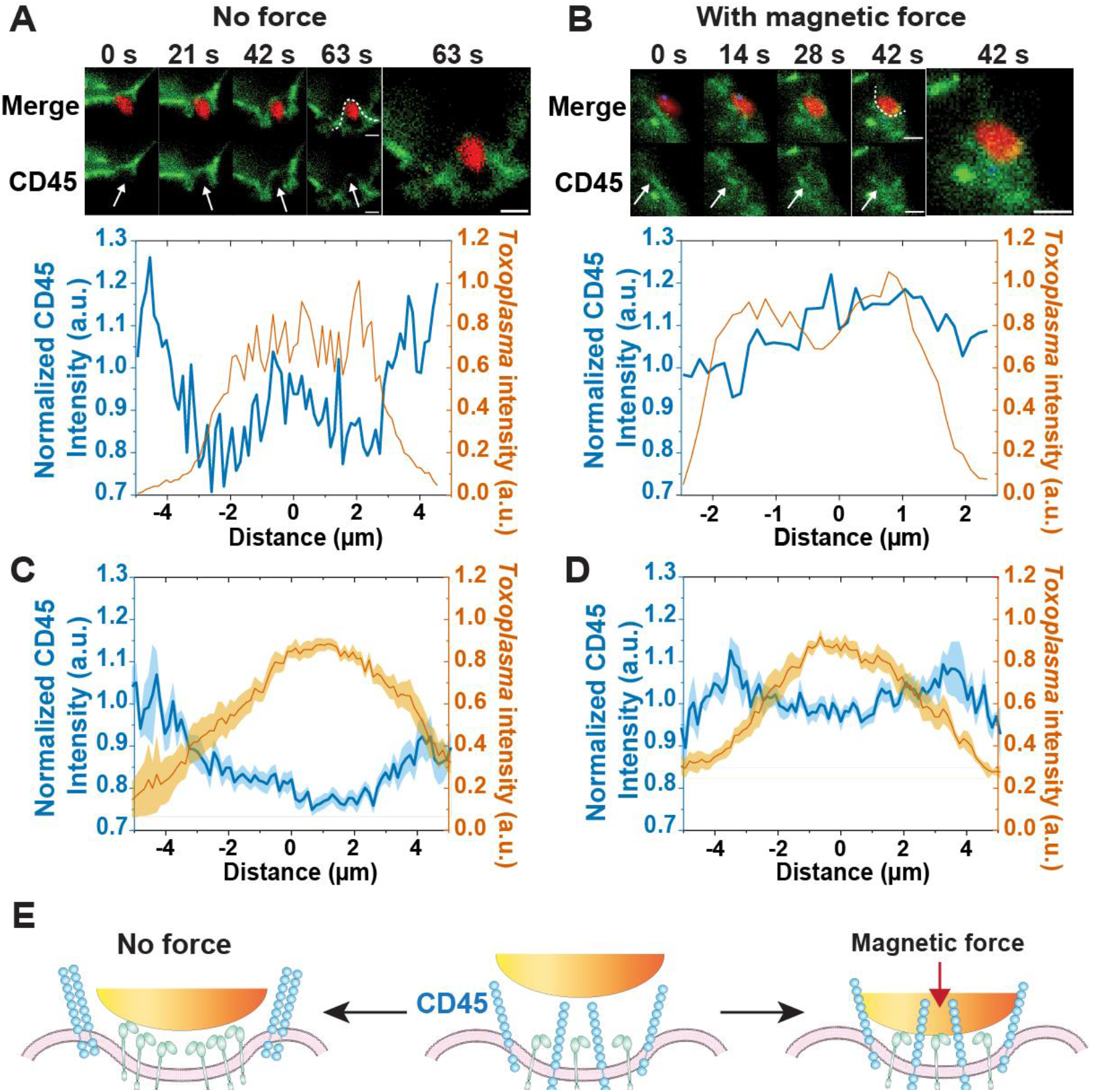
Propulsive force alters the distribution of phosphatase CD45 at the contact site between a heat-killed *Toxoplasma* and a RAW 264.7 cell. (*A* and *B*) Fluorescence images and line-scan intensity plots showing the distribution of CD45 at the phagocytic synapse, without or with magnetic pulling force. White doted lines in fluorescence images indicate the line scans along which the fluorescence intensity of CD45 is plotted. The line-scan intensity plots of *Toxoplasma* are used to indicate the periphery of the cell-parasite contact area. Scale bars, 2 µm. (*C* and *D*) Average line-scan intensity plots showing the average fluorescence intensity of CD45 at the phagocytic synapse, without or with magnetic pulling force. The line curves are averaged from 18 (no force) and 10 (with magnetic force) single phagosome measurements. Shaded areas represent standard error of the mean. (*E*) Schematic illustration showing CD45 distribution at the phagocytic synapse with no force applied and with magnetic force applied.

To further test whether the disruption of phagocytic synapse formation was caused by the physical force exerted or by potential biochemical effects of ligands on *Toxoplasma*, we examined the phagocytic synapse using magnetic beads that were coated with IgG for FcγR-mediated phagocytosis (70, 71). Beads of 2.8 µm in diameter were chosen for better visualization and quantification of the protein distribution at the bead-cell contact site here (force-distance calibration shown in SI Appendix, Fig. S21). The magnetic force applied was estimated to be 100-200 pN (SI Appendix, Fig. S22). We first imaged the distribution of FcγRs by labeling them with a low concentration of F(ab) fragments specific to mouse FcγIII (CD16) and FcγII (CD32) receptors, both of which are expressed on RAW 264.7 cells (86). The F(ab) fragments were fluorescently labeled with CF640R. Without magnetic force, FcγRs clustered at the phagocytic synapse, as indicated by its increased fluorescence intensity (Fig 7 *A* and *E*; SI Appendix, Fig. 23*A*). The clustering of ligand-bound FcγRs is expected for normal phagocytosis. This also confirmed that labeling the receptors with F(ab) fragments did not block their binding to IgG. In contrast, with magnetic pulling, there was no obvious clustering of FcγRs. The fluorescence intensity of FcγRs at the phagocytic synapse remained similar to that in the plasma membrane (Fig 7 *B* and *F*; SI Appendix, Fig. 23*B*). To confirm whether IgG-FcγR bonds formed between the beads and the cell surface, we tried to pull the beads away from the cell surface by reversing the direction of the magnetic force applied. We were unable to pull the beads away, which indicates that FcγRs bound to the IgG on the beads even in the absence of clustering. This suggests that the propulsion force at cell entry did not affect the FcγR recognition of ligands, but blocked receptors outside the bead-cell contact site from moving into the contact site to coalescence. Meanwhile, we found that the spatial exclusion of CD45 at the phagocytic synapse was also hindered under magnetic force (Fig. 7 *C*-*D*, and *G*-*H*). The fact that this phenomenon was observed for both mag-*Toxoplasma* and beads indicates that it is a general observation independent of the specific type of receptor-ligand interaction involved. The results suggest that the propulsive force exerted during internalization does not prevent receptor-ligand binding, but hinders the diffusion of membrane receptors and phosphatases into or out of the site of contact.

**Fig. 7.**
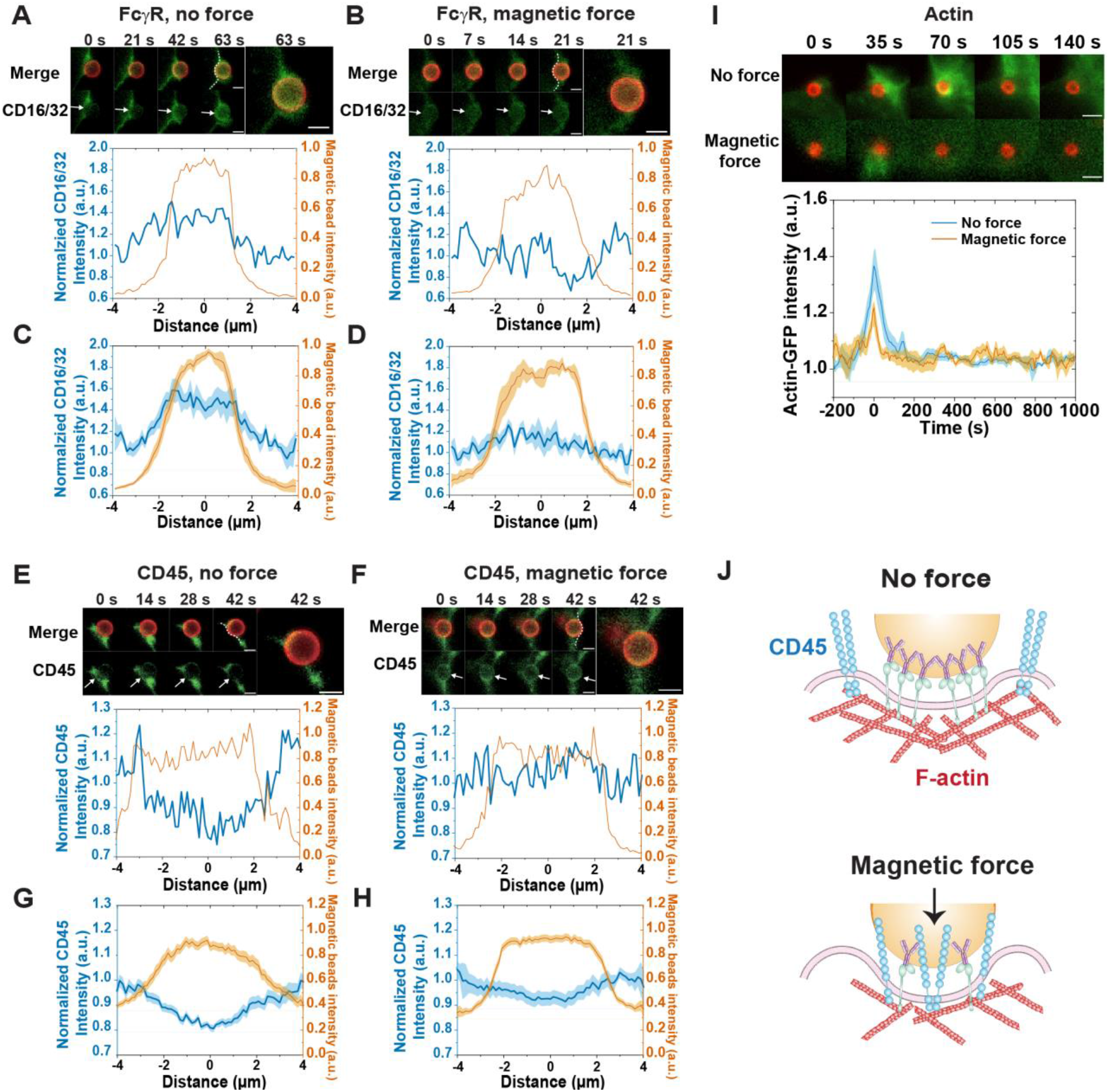
Propulsive force alters the organization of Fc*γ*Rs (CD16/32), CD45, and actin, at the phagocytic synapse. (*A-H*) Fluorescence images and line-scan intensity plots showing the distribution of Fc*γ*Rs (CD16/32) (*A-D*) and CD45 (*E-H*) at the phagocytic synapse during the internalization of IgG opsonized magnetic beads (red), without or with magnetic pulling force. Scale bars, 2 µm. White dotted lines in fluorescence images indicate the line scans along which the fluorescence intensity of receptors is plotted. *A, B, E*, and *F* are single phagosome data. *C, D, G*, and *H* are average line-scan intensity plots showing the average fluorescence intensity of Fc*γ*Rs (CD16/32) (*C, D*) and CD45 (*G, H*) at the phagocytic synapse. Each average line plot is an average of N single phagosome measurements, where N = 10 (C, Fc*γ*R no force), 7 (D, Fc*γ*R with magnetic force), 30 (G, CD45 no force), and 39 (H, CD 45 with magnetic force). The line-scan intensity plots of magnetic beads are used to indicate the periphery of the cell-bead contact area. Shaded areas represent standard error of the mean. (*I*) Fluorescence images showing dynamic remodeling of actin (green) during the internalization of opsonized magnetic beads (red). Scale bars, 2 µm. Line plots show the average actin-GFP accumulation during phagosome formation as a function of time with no force or without magnetic force. The line curves are averaged from N = 12 (no force) and 13 (with magnetic force) single phagosome measurements. Shaded areas represent standard error of the mean. To align the line plots, time zero is defined at the point when actin intensity reaches the peak value. (*J*) Schematic illustrations showing the organization of Fc*γ*Rs (CD16/32), CD45, and actin, at the phagocytic synapse, without or with magnetic pulling force.

Activation of FcγRs during phagocytosis is known to trigger polymerization of actin that pushes the membrane protrusion for engulfment (87, 88). Does CD45 remaining in the bead-cell contact site result in reduced signaling from the FcγRs and subsequently reduced remodeling of actin? We confirmed this by imaging the dynamics of actin-GFP during internalization of beads in live cells. As expected for typical phagocytosis, actin rapidly polymerized around nascent phagosomes and then disassembled (Fig. 7*I*, N = 12; SI Appendix Fig. S24*A*). This peak of actin intensity reportedly coincides with the closure of the phagosome cup (89). However, with applied magnetic force, a significantly reduced amount of actin was observed to polymerize around phagosomes and its presence was more transient, as shown by the decrease in both actin peak intensity and width (Fig. 7*I*, N=13; SI Appendix, Fig. S24*B*). Previous studies have noted that the assembly and disassembly of actin around nascent phagosomes tightly controls the maturation of phagosomes, including the recruitment of Rab5 and phagosome fusion with endosomes and lysosomes (90-92). It is therefore plausible that following the propulsive cell entry of a phagocytic target, the physical force hinders the spatial segregation of activated receptors from phosphatases at the phagocytic synapse, leading to reduced signaling from the receptors and thereby reduced remodeling of actin (schematics shown in Fig. 7*I*). This affects the maturation of nascent phagosomes and diverts them from the canonical phagolysosomal maturation pathway.

### Computational modeling of the impact of propulsive force on the phagocytic synapse formation

We next developed a computational model to understand the mechanism by which propulsion forces impact the organization of receptors and CD45 at the pathogen-cell contact site. We considered apposed regions of the pathogen surface and the macrophage cell surface, and implemented a hybrid simulation framework to characterize the spatiotemporal distribution of surface proteins and the shape of the macrophage membrane. The framework utilized a time-dependent Ginzburg-Landau approach to characterize deformations of the macrophage membrane and the Gillespie method to account for protein diffusion and receptor-ligand binding (detail in SI Appendix) (93, 94). In this model, receptors and CD45 were mobile on the macrophage membrane, receptors can bind to immobile ligands on the pathogen surface, and the macrophage membrane can deform in response to a propulsion force exerted by the rigid pathogen surface and the dynamic distribution of surface proteins. An important feature of the model was that a receptor-ligand bond was smaller in size than CD45, which has a relatively large and rigid extracellular domain. This size difference is the basis of the kinetic segregation model first proposed to understand antigen recognition by T cell receptors (95). To capture the impact of the size difference, we imposed an energy penalty when CD45 was compressed in regions where the surfaces were closer than its natural length (40 nm), or when a receptor-ligand bond was in a region where the distance between surfaces was either larger or smaller than its natural length (15 nm) (96). Mobile surface molecules were more likely to migrate to energetically favorable states than to energetically unfavorable states. In the model, the surface density of CD45 was 54 per µm^2^, based on our experimental estimation of 54 ± 14 CD45 per µm^2^ on RAW264.7 cell surfaces (SI Appendix, Fig. S25). The density of FcγR was 325 per µm^2^, estimated based on measurements reported in the literature (97). We tuned the magnitude of the propulsion force by changing the force constant of a harmonic potential acting on surface elements of the pathogen. Other parameters were obtained from the literature or estimated based on measurements in similar systems (SI Appendix, Table S1).

Our simulations revealed that the spatial organization of surface molecules was modulated by the magnitude of the applied force. For small forces (≤ 7 pN), a central cluster of receptor-ligand bonds formed around the initial cell-pathogen contact site (Fig. 8*A*; SI Appendix, Video S1). The high density of receptor-ligand bonds in this region brought the apposed surfaces to be approximately 15 nm apart. As a result, CD45 molecules were mostly excluded from this region because of their larger, rigid extracellular domain (≈ 40 nm). This spatial organization of surface proteins was a consequence of the growth of the receptor-ligand cluster (SI Appendix, Fig. S26*A*). The initial small number of receptor-ligand bonds held the cell-pathogen interface close together, which facilitated further receptor binding to ligands and caused nearby CD45 to migrate away due to the size discrepancy. The cluster of receptor-ligand bonds grew as unbound receptors diffused into the closely apposed region and bound to ligands near the periphery of the cluster. This growth continued to drive CD45 molecules to the periphery of the central cluster, where the apposed cell surfaces were farther apart. The resulting organization mimics the classic structure of the phagocytic synapse and is consistent with the kinetic segregation model proposed in the context of T-cell activation (96). Weak applied forces (≤ 7 pN) facilitated the formation of additional receptor-ligand bonds by pushing the apposed cell surfaces to be closer near the periphery of the central cluster of ligand-bound receptors. We found that within this weak force regime, larger forces promoted faster growth of the receptor-ligand cluster and increased exclusion of CD45 (Fig. 8*A*; SI Appendix, Videos S1 and S2).

**Fig. 8.**
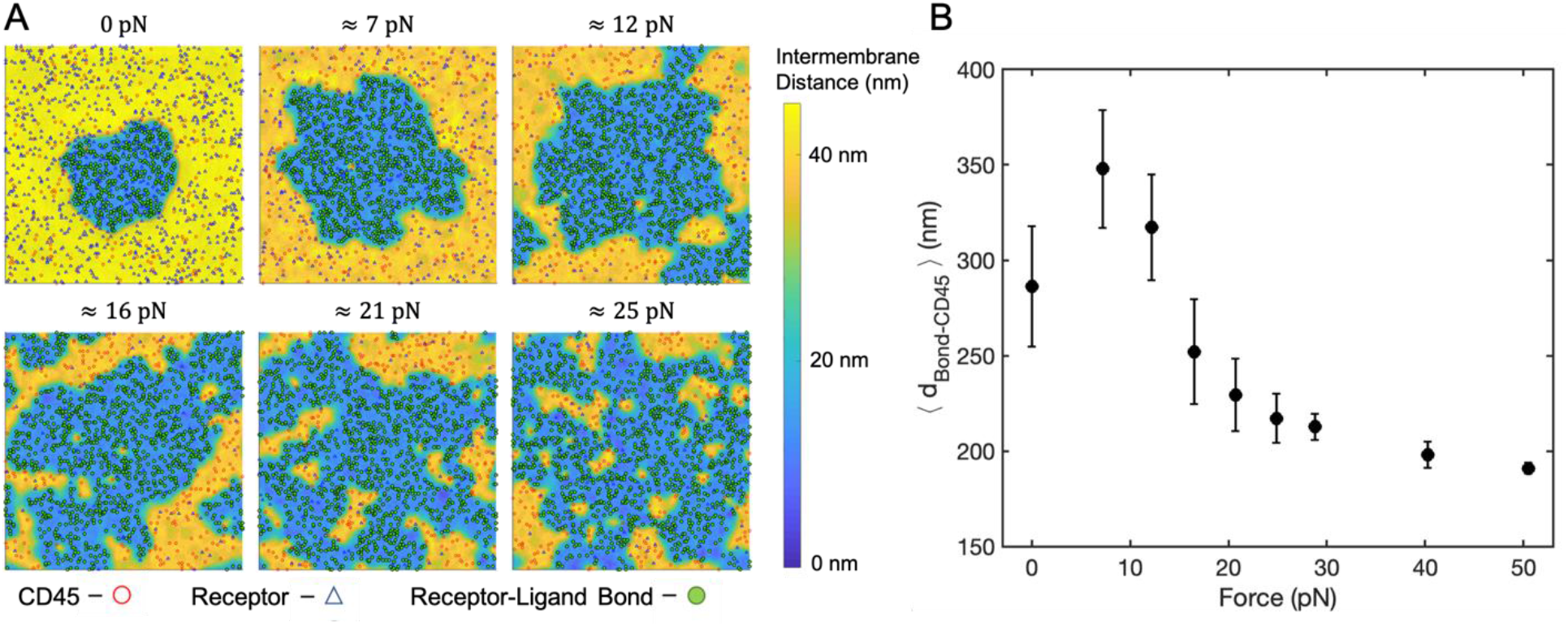
Applied physical force disrupts protein organization of the early phagocytic synapse *in silico*. (*A*) Representative snapshots of simulation trajectories 60 s after initial contact for different magnitudes of applied force. Locations of free receptors (blue triangles), CD45 (red circles), and receptor-ligand bonds (filled green circles) are superimposed on a heat map representing the local distance between the two surfaces. Areas where the distances between the two surfaces are greater than 45 nm are shown in light yellow. Each frame represents a 2 *μ*m × 2 *μ*m patch of membrane. (*B*) The average distance from each receptor-ligand bond to the nearest CD45 molecule (⟨*d*_Bond−CD45_⟩) is plotted as a function of force. Each data point represents the average and standard deviation of 10 independent trajectories.

For larger applied forces (> 7 pN), we observed a change in the organization of surface molecules at the cell-pathogen contact site. Instead of a single, central cluster of receptor-ligand bonds surrounded by spatially excluded CD45 in the peripheral area, the contact region was characterized by multiple CD45-rich regions coexisting with dense clusters of receptor-ligand bonds (Fig. 8*A*). In this large force regime, the applied force rapidly pushed the cell surface and pathogen surface into close contact (SI Appendix, Video S1). This facilitated rapid receptor-ligand binding over a large area extending beyond the initial cluster of bonds (SI Appendix, Fig. S26 *A* and *B*). This caused CD45-rich domains, owing to their large intermembrane distance (≈ 40 nm), to become trapped within the regions of receptor-ligand bonds that were closely apposed (≈ 15 nm). This resulted in irregular and interdigitated boundaries. Differences in the sizes of the surface molecules also resulted in coexisting regions with different intermembrane distances (SI Appendix, Video S2), which restrained reorganization into larger domains because of the energetic cost associated with CD45 crossing the closely apposed region of receptor-ligand bonds. The organization with larger applied forces is consistent with CD45 remaining in the region of contact between the cells, and the coexisting domains appear mixed from a macroscopic perspective.

We quantified the extent of spatial exclusion of CD45 from ligand-bound (activated) receptors by calculating the distance from each receptor-ligand bond to the nearest CD45 after 60 s of simulation time (Fig. 8*B*). Within the small force regime (≤ 7 pN), the spatial separation between the activated receptors and CD45 increased with larger forces (Fig. 8*B*) and increased with time at each given force (SI Appendix, Fig.S26*C*). Both results quantitatively support the observations that the ligand-bound receptors formed a central cluster with CD45 excluded to the periphery, and that larger applied forces caused faster growth of the receptor cluster. Within the large force regime (> 7 pN), the average distance between each ligand-bound receptor and its nearest CD45 was significantly reduced compared to the small force regime, and it decreased with increasing force. This is consistent with the observation of altered surface organization, where multiple CD45-rich domains remained immersed with clusters of receptor-ligand bonds. The time dependence of the average distance (SI Appendix, Fig. S26*C*) showed the rapid formation of coexisting domains of activated receptors and CD45, which then slowly coalesced. Because CD45 is a phosphatase, the smaller distance between CD45 and receptor-ligand bonds at larger forces is consistent with reduced levels of receptor phosphorylation and signaling. Taken together, the simulation results demonstrate that applied force can lead to the reorganization of the surface molecules that control downstream signaling.

## Discussion

In this study, we demonstrate how propulsive cell entry – the physical process by which motile pathogens actively penetrate host cell membranes – redirects the intracellular trafficking of internalized pathogens away from the degradation pathway. This study was motivated by the emerging evidence that the motility of pathogens plays important roles in their invasion of the host. However, how the physical process of pathogen motility might affect the pathogen-host interaction was previously unknown. The major challenge is to differentiate the physical effect of motile pathogens from the biochemical effect of the virulence factors associated with pathogen motility. A widely used approach to study the role of pathogen motility in infection was to generate pathogen mutants with motility impairments. This approach yielded inconclusive results about the role of pathogen motility because the mutants produced still sometimes had residual motility and the mutations could have side effects that influence the pathogen’s efficacy in other ways than through motility.

In this study, we took a new approach by using external magnetic force fields to induce propulsive cell entry in a manner similar to how motile pathogens actively penetrate host cell membranes. We engineered magnetically responsive *Toxoplasma* by conjugating a small number of magnetic nanoparticles onto the surface of the heat-killed parasite. This allowed us to use magnetic tweezers to propel the inactive *Toxoplasma*, like a motile live parasite, during its receptor-mediated internalization in macrophage cells. Using this method, we determined how physical forces exerted during cell entry affect the intracellular fate of pathogens. Because the inactive parasite preserves its surface antigens, this magnetic manipulation approach, combined with computer simulations, allowed us to elucidate the mechanism by which physical forces, as distinct from the effect of the associated virulence factors, contribute to modulating the intracellular trafficking fate of pathogens. We demonstrated the generality of these findings by using both *Toxoplasma* and opsonized beads as phagocytic targets.

A key finding of this study is that physical forces applied during cell entry altered the spatial organization of proteins at the phagocytic synapse. This turned out to be the decision-making point where the internalized pathogens were redirected into vacuoles that failed to mature into degradative units. Through both experiments and simulation, we showed that large propulsion forces, by pushing the pathogen surface close to the host cell membrane, hinder the spatial exclusion of the bulky phosphatase CD45 from activated receptors and thereby limit the productive signaling from the receptors. Previous studies have suggested that the spatial segregation of receptors from phosphatases at the phagocytic synapse is critical for sustaining the activation of receptors (1-3) and promoting phagocytosis (98). Our results here indicate that the formation of a phagocytic synapse is more than just a necessity for regulating receptor signaling at the membrane. It also contributes to determining the intracellular trafficking fate of phagosomes in two plausible ways. First, the phagocytic synapse eventually becomes part of a nascent phagosome. The molecular composition of the phagocytic synapse – the accumulation of receptors and exclusion of phosphatases – determines the membrane composition of the nascent phagosome and likely its destination in intracellular trafficking. Second, activation of receptors at the phagocytic synapse triggers downstream signaling events including the remodeling of actin, which is known to affect the recruitment of the endocytic marker Rab5 to phagosomes and the fusion of phagosomes with lysosomes (90-92, 99). In support of this idea, our results showed that when the spatial exclusion of phosphatase CD45 at the phagocytic synapse was hindered by large propulsion forces, the intensity of transient actin assembly around the nascent phagosomes was reduced. This force-induced reduction in actin remodeling likely resulted in reduced recruitment of signaling proteins, such as the endocytic marker Rab5 that we observed. These signaling proteins are necessary for the maturation of nascent phagosomes. There are no prior studies on how physical forces affect the formation of phagocytic synapses. We elucidated the mechanism within the framework of the kinetic segregation model, which was originally proposed to explain the formation of T cell immune synapse (96). This mechanism focuses on the energetic landscape shaped by the drastic size differences between ligand-bound receptors and the phosphatase CD45. However, it is possible that the applied physical forces can affect the formation of the phagocytic synapse and the assembly of nascent phagosomes via other mechanisms. This will require further exploration. For example, the propulsion force might increase the cell membrane tension at the cell entry site, which has been suggested to affect endocytosis (100).

Along with the force-induced alteration of the phagocytic synapse, a surprising observation is that while inactive parasites that were passively internalized were degraded in phagolysososmes, the same inactive parasites that were actively propelled by magnetic force during cell internalization entered an alternative non-degradative intracellular trafficking pathway. The altered organization at the phagocytic synapse due to propulsion force during cell entry causes a domino effect in preventing the sequential biochemical events that are required to transform a nascent vacuole into a degradative phagolysosome. It disrupted the assembly of endocytic markers, including Rab5 and Rab7, and the lysosomal marker LAMP1 on the surface of the parasite-residing vacuoles. Without those markers, the vacuoles cannot fuse with endosomes and lysosomes. This subsequently prevents the vacuoles from acquiring degradative capacity, which we observed. Without fusion with lysosomes, the vacuoles cannot acquire either the hydrolytic enzymes needed to digest their content (11) nor the ATPases needed to continuously acidify the vacuole lumen (4). We have shown previously that the phagosomal lumen must acidify to an acidic pH of < 5.8 to activate proteolysis and maintain the acidity needed for continuous proteolysis (76). However, the vacuoles formed under propulsive force only reached a final lumen pH of ∼6.5, which is significantly higher than the threshold pH. This chain of disruptive effects highlights the tight regulation of phagosome maturation and the decisive role of cell entry on the intracellular fate of internalized cargos.

We showed that magnetically propelled inactive *Toxoplasma* entered a non-degradative intracellular route strikingly similar to that of live *Toxoplasma*, which is known to actively penetrate host cell membranes during invasion (43-45). Further, by using both the inactivated parasite and beads opsonized with ligands for the Fc*γ*Rs, we showed that this is a general phenomenon, not specific to features of the parasite. This finding suggests that physical forces exerted by motile pathogens during invasion contribute to their evasion of immune degradation. It doesn’t contradict the established evidence that pathogens use virulent factors that perturb one or more steps in the phagosome maturation process to evade degradation (23-26). However, our results suggest a physical effect from the motility of pathogens that can augment the biochemical effect of virulence factors. It is highly plausible that multiple evasion mechanisms are used by a single type of pathogen. Interestingly, we showed in computer simulations that the effect of applied forces can vary depending on the magnitude of the force. In contrast to large forces (> 7 pN) that hinder the protein segregation at the phagocytic synapse, small forces (< 7 pN) can enhance the receptor signaling by promoting receptor-ligand binding and the spatial exclusion of CD45. Forces exerted by motile pathogens are mostly within the large-force range, but the magnitude of forces can vary greatly between different pathogens or even the same pathogens in different growth phases (66, 67). It is therefore possible that the motility of some pathogens makes them more vulnerable to immune degradation. Many pathogens, including the parasite *Toxoplasma* used in our study here, have been shown to use motility to promote their invasion of host cells (27, 28). In contrast to previous studies that focused on how pathogen motility increases their binding to host cells and penetration into host cells, our study provides a new perspective about how the physical process of pathogen motility might affect its intracellular fate after cell entry. This underscores the importance of targeting pathogen movement to combat intracellular pathogens.

## Materials and Methods

Details of reagents, cells, and experimental methods are provided in *SI Appendix*.

### Fabrication of Phagosome Sensors

To fabricate Mag-*Toxoplasma*, heat-killed *Toxoplasma* tachyzoites were first conjugated with fluorescently labeled 200 nm magnetic nanoparticles via EDC reaction. Subsequently, magnetic particles labeled *Toxoplasma* were conjugated with pH sensitive dye pHrodo Red. To prepare FRET-MagSensors for the FRET fusion assay, streptavidin labeled with the FRET donor (ex/em 561/586 nm) Alexa 568 was conjugated on 1 µm Dynabeads.To fabricate pH-MagSensors, Dynabeads with various sizes were conjugated with either pHrodoRed or reference dye CF640R labeled streptavidin. Proteolysis-MagSensors were prepared by labeling 1 µm Dynabeads with streptavidin-CF640R and Z-FR-R110 peptide.

### Live cell fluorescence imaging

Simultaneous imaging of events of phagosome maturation was done using a Nikon Eclipse-Ti inverted microscope equipped with a 1.49 N.A. ×100 TIRF objective and an Andor iXon3 EMCCD camera. RAW264.7 cells were used in magnetic tweezers experiments and control experiments. In FRET-fusion assays, lysosomes were labelled by incubating macrophage cells with biotin-BSA-Alexa 647 in full DMEM overnight followed by a 2h chasing period.

### Magnetic tweezers setup

A magnetic tweezers setup was built on a Nikon Eclipse-Ti inverted microscope. The tip of the magnetic solenoid had a diameter of ≈ 1 µm and the position of the solenoid was controlled using a manual micromanipulator. In experiments, the magnetic force was turned on before phagosome sensors internalization and was kept on during imaging of each sample unless otherwise indicated.

### Image analysis

The localization and fluorescence intensity of phagosome sensors was done using MATLAB algorithms as described previously (101). Analysis of the dynamic distributions of actin filaments and endocytic proteins were performed using custom MTALAB algorithms. Analysis of Fcγ receptor clustering and CD45 exclusion during phagosome formation were analyzed using Image J.

### Simulation

A computational framework was built in which the pathogen surface was in close proximity to a portion of the macrophage surface. Surface proteins were represented as particles that could diffuse; receptors on the macrophage surface could reversibly bind immobile ligands on the pathogen surface. Dynamics of the particles were governed by a stochastic reaction-diffusion process. The shape of the membrane was characterized by a time-dependent Ginzburg-Landau (TDGL) model that accounted for the membrane bending energy and protein deformation energies when surface proteins were located in regions inconsistent with their natural length. The total magnitude of the force by which the pathogen surface was pushed against the macrophage surface was systematically varied.

## ACKNOWLEDGMENTS

We thank the following scientists who have generally contributed reagents and time to make this study possible: Drs. Ke Hu and John Murray at Arizona State University for providing *Toxoplasma* strains, assistance with the parasite culture, and insightful discussion; Dr. Sergio Grinstein at The Hospital for Sick Children Research Institute (Canada) for providing plasmids of endocytic markers and insightful discussion. We thank the following staff members at Indiana University: Andrew Alexander at the Electronic Instrument Services for helping with the setup of magnetic tweezers, Dr. Yi Yi at the Nanoscale Characterization Center for assistance with scanning electron microscopy, and Dr. Giovanni Gonzalez-Gutierrez at the Physical Biochemistry Instrumentation Facility for assistance with Nanodrop. This work was supported by National Institutes of Health award R35GM124918 to Y. Y. and National Science Foundation award PHY1753017 to S.M.A. The content is solely the responsibility of the authors and does not necessarily represent the official views of the National Institutes of Health.

## Notes

### Competing Interest Statement

The authors have declared no competing interest.

## References

1. H. S. Goodridge et al., Activation of the innate immune receptor Dectin-1 upon formation of a ‘phagocytic synapse’. Nature 472, 471–475 (2011).

2. M. H. Bakalar et al., Size-Dependent Segregation Controls Macrophage Phagocytosis of Antibody-Opsonized Targets. Cell 174, 131–142 e113 (2018).

3. S. A. Freeman et al., Integrins Form an Expanding Diffusional Barrier that Coordinates Phagocytosis. Cell 164, 128–140 (2016).

4. B. Steinberg, K. Huynh, S. Grinstein (2007) Phagosomal acidification: measurement, manipulation and functional consequences. (Portland Press Ltd.).

5. S. Duclos et al., Rab5 regulates the kiss and run fusion between phagosomes and endosomes and the acquisition of phagosome leishmanicidal properties in RAW 264.7 macrophages. Journal of Cell Science 113, 3531–3541 (2000).

6. M. C. Kielian, Z. A. Cohn, Phagosome-Lysosome Fusion - Characterization of Intracellular Membrane-Fusion in Mouse Macrophages. Journal of Cell Biology 85, 754–765 (1980).

7. K. K. Huynh et al., LAMP proteins are required for fusion of lysosomes with phagosomes. EMBO J 26, 313–324 (2007).

8. V. Claus et al., Lysosomal enzyme trafficking between phagosomes, endosomes, and lysosomes in J774 macrophages - Enrichment of cathepsin H in early endosomes. Journal of Biological Chemistry 273, 9842–9851 (1998).

9. H. J. Ullrich, W. L. Beatty, D. G. Russell, Direct delivery of procathepsin D to phagosomes: Implications for phagosome biogenesis and parasitism by Mycobacterium. European Journal of Cell Biology 78, 739–748 (1999).

10. R. M. Yates, A. Hermetter, D. G. Russell, The kinetics of phagosome maturation as a function of phagosome/lysosome fusion and acquisition of hydrolytic activity. Traffic 6, 413–420 (2005).

11. A. M. Lennon-Dumenil et al., Analysis of protease activity in live antigen-presenting cells shows regulation of the phagosomal proteolytic contents during dendritic cell activation. J Exp Med 196, 529–540 (2002).

12. C. Persson et al., Cell-surface-bound Yersinia translocate the protein tyrosine phosphatase YopH by a polarized mechanism into the target cell. Molecular microbiology 18, 135–150 (1995).

13. S. Håkansson, E. E. Galyov, R. Rosqvist, H. Wolf-Watz, The Yersinia YpkA Ser/Thr kinase is translocated and subsequently targeted to the inner surface of the HeLa cell plasma membrane. Molecular microbiology 20, 593–603 (1996).

14. E. Frithz-Lindsten, Y. Du, R. Rosqvist, Å. Forsberg, Intracellular targeting of exoenzyme S of Pseudomonas aeruginosa via type III-dependent translocation induces phagocytosis resistance, cytotoxicity and disruption of actin microfilaments. Molecular microbiology 25, 1125–1139 (1997).

15. L. Garrity-Ryan et al., The arginine finger domain of ExoT contributes to actin cytoskeleton disruption and inhibition of internalization of Pseudomonas aeruginosa by epithelial cells and macrophages. Infection and immunity 68, 7100–7113 (2000).

16. N. W. Andrews, C. K. Abrams, S. L. Slatin, G. Griffiths, A T. cruzi-secreted protein immunologically related to the complement component C9: evidence for membrane pore-forming activity at low pH. Cell 61, 1277–1287 (1990).

17. B. F. Hall, P. Webster, A. K. Ma, K. A. Joiner, N. W. Andrews, Desialylation of lysosomal membrane glycoproteins by Trypanosoma cruzi: a role for the surface neuraminidase in facilitating parasite entry into the host cell cytoplasm. The Journal of experimental medicine 176, 313–325 (1992).

18. M. J. McConville, S. J. Turco, M. Ferguson, D. L. Sacks, Developmental modification of lipophosphoglycan during the differentiation of Leishmania major promastigotes to an infectious stage. The EMBO journal 11, 3593–3600 (1992).

19. B. S. McGwire, K.-P. Chang, D. M. Mosser, Role of the Leishmania Surface Protease gp63 in Complement Fixation, Cell Adhesion, and Resistance to Complement-Mediated Lysis’. (1995).

20. K. A. Mcdonough, Y. Kress, B. Bloom, Pathogenesis of tuberculosis: interaction of Mycobacterium tuberculosis with macrophages. Infection and immunity 61, 2763–2773 (1993).

21. I. Vergne et al., Mechanism of phagolysosome biogenesis block by viable Mycobacterium tuberculosis. Proceedings of the National Academy of Sciences of the United States of America 102, 4033–4038 (2005).

22. L. Staali, S. Bauer, M. Mörgelin, L. Björck, H. Tapper, Streptococcus pyogenes bacteria modulate membrane traffic in human neutrophils and selectively inhibit azurophilic granule fusion with phagosomes. Cellular microbiology 8, 690–703 (2006).

23. E. Hertzén et al., M1 protein-dependent intracellular trafficking promotes persistence and replication of Streptococcus pyogenes in macrophages. Journal of innate immunity 2, 534–545 (2010).

24. D. Wong, H. Bach, J. Sun, Z. Hmama, Y. Av-Gay, Mycobacterium tuberculosis protein tyrosine phosphatase (PtpA) excludes host vacuolar-H+-ATPase to inhibit phagosome acidification. P Natl Acad Sci USA 108, 19371–19376 (2011).

25. A. Mehra et al., Mycobacterium tuberculosis Type VII Secreted Effector EsxH Targets Host ESCRT to Impair Trafficking. Plos Pathogens 9 (2013).

26. A. Gallois, J. R. Klein, L.-A. H. Allen, B. D. Jones, W. M. Nauseef, Salmonella pathogenicity island 2-encoded type III secretion system mediates exclusion of NADPH oxidase assembly from the phagosomal membrane. The journal of immunology 166, 5741–5748 (2001).

27. C. Josenhans, S. Suerbaum, The role of motility as a virulence factor in bacteria. International Journal of Medical Microbiology 291, 605–614 (2002).

28. D. Sacks, A. Sher, Evasion of innate immunity by parasitic protozoa. Nat Immunol 3, 1041–1047 (2002).

29. C. Dietrich, K. Heuner, B. C. Brand, J. Hacker, M. Steinert, Flagellum of Legionella pneumophila positively affects the early phase of infection of eukaryotic host cells. Infection and Immunity 69, 2116–2122 (2001).

30. J. A. Arias-del-Angel, J. Santana-Solano, M. Santillan, R. G. Manning-Cela, Motility patterns of Trypanosoma cruzi trypomastigotes correlate with the efficiency of parasite invasion in vitro. Sci Rep-Uk 10 (2020).

31. S. L. Hoffman, G. M. Subramanian, F. H. Collins, J. C. Venter, Plasmodium, human and Anopheles genomics and malaria. Nature 415, 702–709 (2002).

32. J. K. Griffiths, Human cryptosporidiosis: Epidemiology, transmission, clinical disease, treatment, and diagnosis. Adv Parasit 40, 37–85 (1998).

33. D. Soldati, B. J. Foth, A. F. Cowman, Molecular and functional aspects of parasite invasion. Trends in Parasitology 20, 567–574 (2004).

34. M. B. Heintzelman, Gliding motility in apicomplexan parasites. Seminars in Cell & Developmental Biology 46, 135–142 (2015).

35. K. Frenal, J. F. Dubremetz, M. Lebrun, D. Soldati-Favre, Gliding motility powers invasion and egress in Apicomplexa. Nature Reviews Microbiology 15, 645–660 (2017).

36. M. M. Shimogawa et al., Parasite motility is critical for virulence of African trypanosomes. Scientific Reports 8, 9122 (2018).

37. J. M. Dobrowolski, V. B. Carruthers, L. D. Sibley, Participation of myosin in gliding motility and host cell invasion by Toxoplasma gondii. Mol Microbiol 26, 163–173 (1997).

38. S. Mehta, L. D. Sibley, Actin depolymerizing factor controls actin turnover and gliding motility in Toxoplasma gondii. Mol Biol Cell 22, 1290–1299 (2011).

39. J. A. Whitelaw et al., Surface attachment, promoted by the actomyosin system of Toxoplasma gondii is important for efficient gliding motility and invasion. Bmc Biol 15 (2017).

40. L. M. Weiss, J. P. Dubey, Toxoplasmosis: A history of clinical observations. Int J Parasitol 39, 895–901 (2009).

41. C. Contini, Clinical and diagnostic management of toxoplasmosis in the immunocompromised patient. Parassitologia 50, 45–50 (2008).

42. G. Saadatnia, M. Golkar, A review on human toxoplasmosis. Scand J Infect Dis 44, 805–814 (2012).

43. D. G. Mordue, N. Desai, M. Dustin, L. D. Sibley, Invasion by Toxoplasma gondii establishes a moving junction that selectively excludes host cell plasma membrane proteins on the basis of their membrane anchoring. J Exp Med 190, 1783–1792 (1999).

44. L. D. Sibley, Intracellular parasite invasion strategies. Science 304, 248–253 (2004).

45. J. M. Dobrowolski, L. D. Sibley, Toxoplasma invasion of mammalian cells is powered by the actin cytoskeleton of the parasite. Cell 84, 933–939 (1996).

46. L. D. Sibley, E. Weidner, J. L. Krahenbuhl, Phagosome Acidification Blocked by Intracellular Toxoplasma-Gondii. Nature 315, 416–419 (1985).

47. K. A. Joiner, S. A. Fuhrman, H. M. Miettinen, L. H. Kasper, I. Mellman, Toxoplasma-Gondii - Fusion Competence of Parasitophorous Vacuoles in Fc-Receptor Transfected Fibroblasts. Science 249, 641–646 (1990).

48. D. G. Mordue, S. Hakansson, I. Niesman, L. D. Sibley, Toxoplasma gondii resides in a vacuole that avoids fusion with host cell endocytic and exocytic vesicular trafficking pathways. Exp Parasitol 92, 87–99 (1999).

49. I. Coppens et al., Toxoplasma gondii sequesters lysosomes from mammalian hosts in the vacuolar space. Cell 125, 261–274 (2006).

50. D. G. Mordue, L. D. Sibley, Intracellular fate of vacuoles containing Toxoplasma gondii is determined at the time of formation and depends on the mechanism of entry. Journal of Immunology 159, 4452–4459 (1997).

51. B. A. Butcher, E. Y. Denkers, Mechanism of entry determines the ability of Toxoplasma gondii to inhibit macrophage proinflammatory cytokine production. Infect Immun 70, 5216–5224 (2002).

52. N. K. Kisalu, G. Langousis, L. A. Bentolila, K. S. Ralston, K. L. Hill, Mouse infection and pathogenesis by Trypanosoma brucei motility mutants. Cellular Microbiology 16, 912–924 (2014).

53. R. A. Fratti, J. M. Backer, J. Gruenberg, S. Corvera, V. Deretic, Role of phosphatidylinositol 3-kinase and Rab5 effectors in phagosomal biogenesis and mycobacterial phagosome maturation arrest. J Cell Biol 154, 631–644 (2001).

54. O. V. Vieira et al., Modulation of Rab5 and Rab7 recruitment to phagosomes by phosphatidylinositol 3-kinase. Mol Cell Biol 23, 2501–2514 (2003).

55. A. Vonderheit, A. Helenius, Rab7 associates with early endosomes to mediate sorting and transport of Semliki forest virus to late endosomes. PLoS Biol 3, e233 (2005).

56. P. Cossart, A. Helenius, Endocytosis of viruses and bacteria. Cold Spring Harb Perspect Biol 6 (2014).

57. A. M. Pauwels, M. Trost, R. Beyaert, E. Hoffmann, Patterns, Receptors, and Signals: Regulation of Phagosome Maturation. Trends Immunol 38, 407–422 (2017).

58. A. Jahraus et al., In vitro fusion of phagosomes with different endocytic organelles from J774 macrophages. J Biol Chem 273, 30379–30390 (1998).

59. M. Desjardins, N. N. Nzala, R. Corsini, C. Rondeau, Maturation of phagosomes is accompanied by changes in their fusion properties and size-selective acquisition of solute materials from endosomes. Journal of Cell Science 110, 2303–2314 (1997).

60. Y. K. Oh et al., Rapid and complete fusion of macrophage lysosomes with phagosomes containing Salmonella typhimurium. Infection and Immunity 64, 3877–3883 (1996).

61. R. Garcia-Rodas, F. Gonzalez-Camacho, J. L. Rodriguez-Tudela, M. Cuenca-Estrella, O. Zaragoza, The Interaction between Candida krusei and Murine Macrophages Results in Multiple Outcomes, Including Intracellular Survival and Escape from Killing. Infection and Immunity 79, 2136–2144 (2011).

62. C. Zahn et al., Measurement of the magnetic moment of single Magnetospirillum gryphiswaldense cells by magnetic tweezers. Sci Rep 7, 3558 (2017).

63. A. R. Bausch, W. Moller, E. Sackmann, Measurement of local viscoelasticity and forces in living cells by magnetic tweezers. Biophysical Journal 76, 573–579 (1999).

64. P. Kollmannsberger, B. Fabry, High-force magnetic tweezers with force feedback for biological applications. Rev Sci Instrum 78, 114301 (2007).

65. C. Jiang, T. A. Lionberger, D. M. Wiener, E. Meyhofer, Electromagnetic tweezers with independent force and torque control. Rev Sci Instrum 87, 084304 (2016).

66. R. V. Stadler, L. A. White, K. Hu, B. P. Helmke, W. H. Guilford, Direct measurement of cortical force generation and polarization in a living parasite. Molecular Biology of the Cell 28, 1912–1923 (2017).

67. M. Mizutani, Y. Sasajima, M. Miyata, Force and Stepwise Movements of Gliding Motility in Human Pathogenic Bacterium Mycoplasma pneumoniae. Front Microbiol 12, 747905 (2021).

68. I. Obataya, C. Nakamura, S. Han, N. Nakamura, J. Miyake, Nanoscale operation of a living cell using an atomic force microscope with a nanoneedle. Nano Letters 5, 27–30 (2005).

69. D. Gonzalez-Rodriguez et al., Mechanical Criterion for the Rupture of a Cell Membrane under Compression. Biophysical Journal 111, 2711–2721 (2016).

70. K. Newton, V. M. Dixit, Signaling in Innate Immunity and Inflammation. Csh Perspect Biol 4 (2012).

71. E. Garcia-Garcia, C. Rosales, Signal transduction during Fc receptor-mediated phagocytosis. J Leukocyte Biol 72, 1092–1108 (2002).

72. Y. Yu, Z. Zhang, G. F. W. Walpole, Y. Yu, Kinetics of phagosome maturation is coupled to their intracellular motility. Communications Biology 5, 1014 (2022).

73. G. L. Lukacs, O. D. Rotstein, S. Grinstein, Phagosomal Acidification Is Mediated by a Vacuolar-Type H+-Atpase in Murine Macrophages. Journal of Biological Chemistry 265, 21099–21107 (1990).

74. G. H. Sun-Wada, H. Tabata, N. Kawamura, M. Aoyama, Y. Wada, Direct recruitment of H+-ATPase from lysosomes for phagosomal acidification. J Cell Sci 122, 2504–2513 (2009).

75. B. E. Steinberg, K. K. Huynh, S. Grinstein, Phagosomal acidification: measurement, manipulation and functional consequences. Biochem Soc T 35, 1083–1087 (2007).

76. S. Lee, Z. Zhang, Y. Yu, Real-time Simultaneous Imaging of Acidification and Proteolysis in Single Phagosomes Using Bifunctional Janus Particle Probes. Angew Chem Int Ed Engl 10.1002/anie.202111094 (2021).

77. A. Savina et al., NOX2 controls phagosomal pH to regulate antigen processing during crosspresentation by dendritic cells. Cell 126, 205–218 (2006).

78. C. G. Zou, Y. C. Ma, L. L. Dai, K. Q. Zhang, Autophagy protects C. elegans against necrosis during Pseudomonas aeruginosa infection. P Natl Acad Sci USA 111, 12480–12485 (2014).

79. C. J. Luke et al., An intracellular serpin regulates necrosis by inhibiting the induction and sequelae of lysosomal injury. Cell 130, 1108–1119 (2007).

80. P. L. McNeil, L. Tanasugarn, J. B. Meigs, D. L. Taylor, Acidification of phagosomes is initiated before lysosomal enzyme activity is detected. J Cell Biol 97, 692–702 (1983).

81. R. K. Tsai, D. E. Discher, Inhibition of “self” engulfment through deactivation of myosin-II at the phagocytic synapse between human cells. Journal of Cell Biology 180, 989–1003 (2008).

82. M. S. Itano et al., Super-resolution imaging of C-type lectin spatial rearrangement within the dendritic cell plasma membrane at fungal microbe contact sites. Front Phys 2 (2014).

83. E. D. C. Mcfarland et al., Correlation between Src Family Member Regulation by the Protein-Tyrosine-Phosphatase Cd45 and Transmembrane Signaling through the T-Cell Receptor. P Natl Acad Sci USA 90, 1402–1406 (1993).

84. P. Shrivastava, T. Katagiri, M. Ogimoto, K. Mizuno, H. Yakura, Dynamic regulation of Src-family kinases by CD45 in B cells. Blood 103, 1425–1432 (2004).

85. M. L. Hermiston, J. Zikherman, J. W. Zhu, CD45, CD148, and Lyp/Pep: critical phosphatases regulating Src family kinase signaling networks in immune cells (vol 228, pg 288, 2009). Immunological Reviews 229, 387–387 (2009).

86. S. A. Freeman et al., Transmembrane Pickets Connect Cyto- and Pericellular Skeletons Forming Barriers to Receptor Engagement. Cell 172, 305–317 e310 (2018).

87. H. Park, D. Cox, Cdc42 Regulates Fc(gamma) Receptor-mediated Phagocytosis through the Activation and Phosphorylation of Wiskott-Aldrich Syndrome Protein (WASP) and Neural-WASP. Molecular Biology of the Cell 20, 4500–4508 (2009).

88. S. A. Freeman, S. Grinstein, Phagocytosis: receptors, signal integration, and the cytoskeleton. Immunological Reviews 262, 193–215 (2014).

89. C. C. Scott et al., Phosphatidylinositol-4,5-bisphosphate hydrolysis directs actin remodeling during phagocytosis. J Cell Biol 169, 139–149 (2005).

90. I. Guérin, C. de Chastellier, Disruption of the actin filament network affects delivery of endocytic contents marker to phagosomes with early endosome characteristics: the case of phagosomes with pathogenic mycobacteria. European journal of cell biology 79, 735–749 (2000).

91. M. Kitano, M. Nakaya, T. Nakamura, S. Nagata, M. Matsuda, Imaging of Rab5 activity identifies essential regulators for phagosome maturation. Nature 453, 241–245 (2008).

92. Y. Yu, Z. Zhang, Y. Yu, Maturation kinetics of phagosomes depends on their transport switching from actin to microtubule tracks. bioRxiv, 2022.2005. 2013.491905 (2022).

93. D. T. Gillespie, Stochastic Simulation of Chemical Kinetics. Annu. Rev. Phys. Chem. 58, 35–55 (2007).

94. S. Raychaudhuri, A. K. Chakraborty, M. Kardar, Effective Membrane Model of the Immunological Synapse. Phys. Rev. Lett. 91, 208101 (2003).

95. S. J. Davis, P. A. van der Merwe, The kinetic-segregation model: TCR triggering and beyond. Nature Immunology 7, 803–809 (2006).

96. R. H. Pullen, S. M. Abel, Catch Bonds at T Cell Interfaces: Impact of Surface Reorganization and Membrane Fluctuations. Biophys. J. 113, 120–131 (2017).

97. F. Krombach et al., Cell size of alveolar macrophages: An interspecies comparison. Environ Health Persp 105, 1261–1263 (1997).

98. M. H. Bakalar et al., Size-Dependent Segregation Controls Macrophage Phagocytosis of Antibody-Opsonized Targets. Cell 174, 131–142.e113 (2018).

99. D. Liebl, G. Griffiths, Transient assembly of F-actin by phagosomes delays phagosome fusion with lysosomes in cargo-overloaded macrophages. J Cell Sci 122, 2935–2945 (2009).

100. U. Djakbarova, Y. Madraki, E. T. Chan, C. Kural, Dynamic interplay between cell membrane tension and clathrin-mediated endocytosis. Biol Cell 113, 344–373 (2021).

101. Y. Q. Yu, Y. Gao, Y. Yu, “Waltz” of Cell Membrane-Coated Nanoparticles on Lipid Bilayers: Tracking Single Particle Rotation in Ligand-Receptor Binding. Acs Nano 12, 11871–11880 (2018).

