## Supplemental Information for "Propulsive cell entry diverts pathogens from immune degradation by remodeling the phagocytic synapse"

**This PDF file includes:**

- Materials and Methods
- Table S1
- Figures S1 to S26
- Caption for videos S1 and S2
- SI References

### Materials and Methods

#### 1. Materials

Carboxylate-modified superparamagnetic Dynabeads (diameter 1 µm), pentylamine-biotin, Alexa Fluor 568 NHS ester (succinimidyl ester), Alexa Fluor 647 NHS ester (succinimidyl ester), pHrodo iFL Red STP Ester, *Toxoplasma gondii* SAG1 Monoclonal Antibody (D61S), Alexa647 conjugated chicken anti-mouse IgG secondary antibody, Pierce Fab preparation kit were purchased from ThermoFisher (Waltham, MA). Immunoglobulin G from rabbit plasma, albumin from bovine serum (BSA), and biotin N-hydroxysuccinimide ester (biotin-NHS) were purchased from Sigma-Aldrich (St. Louis, MO). CF488 NHS ester, CF640R amine, CF640R NHS ester, CF488A labeled Cholera Toxin Subunit B, and fluorescent peptide Rhodamine 110, bis-(N-CBZ-L-phenylalanyl-L-arginine amide) dihydrochloride (Z-FR-R110) were purchased from Biotium (Fremont, CA). 200 nm Super Mag Carboxylic Acid Beads were purchased from Ocean NanoTech (San Diego, CA), Streptavidin was purchased from MyBioSource (San Diego, CA). Alexa647 conjugated rat anti-mouse CD45 antibody was purchased from BioLegend (San Diego, CA). Recombinant rabbit anti-mouse CD16/32 antibody (2.4G2) was purchased from Novus Biologicals (Centennial CO). Sylgard 184 PDMS base was from Dow Corning (Midland, MI). 1-(3-Dimethylaminopropyl)-3-ethylcarbodiimide hydrochloride (EDC) was purchased from Alfa Aesar (Haverhill, MA). Nigericin sodium salt was purchased from Tocris Bioscience (Minneapolis, MN). FuGENE HD transfection reagent was purchased from Promega (Madison, WI). GFP-Rab5, RFP-Rab5, GFP-Rab7, RFP-Rab7, GFP-LAMP1, and RFP-LAMP1 were prepared as previously described (1, 2). PM–RFP, which contains a ten amino acid sequence (MGCIKSKRKD) from the conserved palmitoylation and myristoylation sequences from Lyn at the N-terminus of RFP, has been previously described (3, 4). RAW264.7 macrophage cells were purchased from ATCC (Manassas, VA). RAW264.7 macrophages stably expressing EGFP-actin and RFP-actin have been previously described (5). RH strain and tubulin- GFP expressing *Toxoplasma gondii* and Human foreskin fibroblast (HFF) cells are cultured as described (6). Tubulin-GFP expressing *Toxoplasma gondii* were selected from RHΔHX stably transfected with phr6_mNeonGreenFP_TgTUBA1 with mycophenolic acid + xanthine. Ringer's imaging buffer (10 mM HEPES, 10 mM glucose, 155 mM NaCl, 2 mM $NaH_2PO_4$, 5 mM KCl, 2 mM $CaCl_2$, 1 mM $MgCl_2$, pH = 7.2) was used for live-cell imaging. Potassium-rich solution (135 mM KCl, 2 mM $K_2HPO_2$, 1.2 mM $CaCl_2$, 0.8 mM $MgSO_4$) was used for intracellular pH calibration.

#### 2. Cell culture and transfection

All RAW 264.7 macrophages cells were cultured in Dulbecco's Modified Eagle Medium (DMEM) complete medium supplemented with 10% fetal bovine serum (FBS), 2 mM L-Glutamine, 100 units/ml penicillin, and 100 µg/ml streptomycin at 37°C with 5% $CO_2$. For cell transfection, 1 million RAW 264.7 macrophage cells were seeded on cleaned glass coverslip for 3h prior to transfection. Transfection was performed according to manufacturer's instructions. Briefly, 3 µl Fugene HD transfection reagent and 1000 ng of plasmid were mixed in 100 µl DMEM and kept at room temperature for 15 min. After that, the mixture was added gently to cells and incubated with cells in a 37°C with 5% $CO_2$ for 18 h before live cell imaging.

#### 3. *Toxoplasma gondii* cell culture and isolation

*Toxoplasma gondii* were cultured and maintained by serial passage in human foreskin fibroblast (HFF) cells in DMEM supplemented with 1% fetal bovine serum (FBS), 2 mM L-Glutamine, 100 units/ml penicillin, and 100 µg/ml streptomycin at 37°C with 5% $CO_2$. HFF cells alone were cultured in DMEM supplemented with 10% fetal bovine serum (FBS), 2 mM L-Glutamine, 100 units/ml penicillin, and 100 µg/ml streptomycin at 37°C with 5% $CO_2$.

Parasites were isolated from lysed HFF cell suspension. In brief, the lysed suspension was first passed through a 25-G needle using a syringe and subsequently filtered using a 3-µm pore size polycarbonate filter (Whatman Nuclepore). For live *Toxoplasma* invasion experiment, freshly harvested parasites were used immediately after isolation. To prepare heat-killed *Toxoplasma* tachyzoites, parasites after filtration were heated at 61 °C for 10 min and then stored in 1xPBS for future use.

#### 4. Fabrication and characterization of phagosome sensors

### 4.1 Fluorescence labeling of proteins

Streptavidin-CF640R conjugates were synthesized by mixing streptavidin (reaction concentration: 2 mg/ml), CF640R amine (reaction concentration: 0.23 mg/ml), and EDC (reaction concentration: 1.5 mg/ml) in MES buffer (50mM, pH 4.5) for 3h. To prepare streptavidin-Alexa568 conjugates, streptavidin (reaction concentration: 2.1 mg/ml) and Alexa568 NHS ester (reaction concentration: 0.26 mg/ml) were mixed in sodium bicarbonate buffer (0.1 M, pH 8.25) for 1 h at room temperature. Streptavidin-pHrodo Red conjugates were prepared via amine-NHS coupling by mixing streptavidin (reaction concentration: 2 mg/ml) and pHrodo Red STP (reaction concentration: 0.25 mg/ml) in sodium bicarbonate buffer (0.1 M, pH 8.25) for 3 h. Streptavidin-CF488A conjugates were synthesized by mixing streptavidin (reaction concentration: 2 mg/ml) and CF488A NHS ester (reaction concentration: 0.25 mg/ml) in sodium bicarbonate buffer (0.1 M, pH 8.25) for 3 h at room temperature. In all protein labeling procedures described above, free dyes were removed by centrifugal filtration using Amicon Ultra filters (30K). To prepare bovine serum albumin-biotin-Alexa647 (BSA-biotin-Alexa647) conjugates BSA (reaction concentration: 2 mg/ml) and biotin-NHS (reaction concentration: 0.19 mg/ml) were mixed in sodium bicarbonate buffer (0.1 M, pH 8.2) at room temperature for 1 h. Unbound biotin-NHS was removed by centrifugal filtration using Amicon Ultra filters (30K). Subsequently, BSA-biotin (reaction concentration: 2 mg/ml) and Alexa647 NHS ester (reaction concentration: 0.78 mg/ml) were mixed in sodium bicarbonate buffer (0.1 M, pH 8.2) at room temperature for 3h. Free dyes were removed by centrifugal filtration using Amicon Ultra filters (30K). The F(ab) fragments of anti-CD16/32 antibody were prepared using the Pierce Fab preparation kit following manufacturer protocol and subsequently labeled with a fluorescent dye CF640R following the dye-labeling protocol described above.

### 4.2. Magnetically (Mag-) and pH responsive (pH-) heat-killed *Toxoplasma* tachyzoites

**Mag-*Toxoplasma* tachyzoites:** First, 200 nm magnetic nanoparticles were conjugated with fluorescently labeled streptavidin. To convert carboxylic groups on magnetic particles to biotin groups, magnetic particles (reaction concentration: 1 mg/ml) were rinsed three times with methanol and MES buffer (50 mM, pH 6.2) and resuspended in MES buffer (50 mM, pH 6.2) containing EDC (reaction concentration: 8 mg/ml) and of biotin pentylamine (reaction concentration: 1 mM). After the biotinylation, streptavidin-CF640R (reaction concentration: 50 µg/ml) or streptavidin-CF488A (reaction concentration: 20 µg/ml) and BSA (reaction concentration: 0.1 mg/ml) were mixed into the particle solution. The reaction was performed at room temperature for 1 h. Second, fluorescently labeled magnetic nanoparticles were conjugated on heat-killed *Toxoplasma* tachyzoites. In brief, heat-killed *Toxoplasma* tachyzoites (reaction concentration: 160 million/ml) were mixed with streptavidin-CF640R labeled magnetic particles at 1:400 molar ratio in MES buffer (50 mM, pH 6.2) containing 4 mg/ml EDC in room temperature for 1h. To subsequently label magnetically responsive *Toxoplasma* tachyzoites with pHrodo Red, magnetic particles labeled *Toxoplasma* tachyzoites (reaction concentration: 160 million/ml) were resuspended in sodium bicarbonate buffer (0.1 M, pH 8.25). Subsequently, pHrodo Red STP dissolved in DMSO (reaction concentration: 20 µg/ml) was added to solution and the reaction was carried out at room temperature for 2 h. Free dyes were removed by washing tachyzoites three times with 1xPBS. To label heat-killed *Toxoplasma* tachyzoites with Alexa568 NHS ester, Mag- *Toxoplasma* tachyzoites (reaction concentration: 160 million/ml) were resuspended in sodium bicarbonate buffer (0.1 M, pH 8.25) containing Alexa568 (reaction concentration: 20 µg/ml). The mixture was incubated at room temperature for 2h. Free dyes were removed by washing tachyzoites three times with 1xPBS.

### 4.3. Magnetically modulated phagosome sensors (MagSensors)

**FRET-MagSensors:** To convert carboxylic groups on 1 µm Dynabeads to biotin groups, stock Dynabeads (reaction concentration: 1 mg/ml) were washed three times with MES buffer (50 mM, pH 6.2) and resuspended in MES buffer (50 mM, pH 6.2) containing EDC (reaction concentration: 10 mg/ml) and of biotin pentylamine (reaction concentration: 1 mM). The mixture was incubated at room temperature for 1 h. After that, biotinylated Dynabeads were rinsed and resuspended into of 1×PBS buffer containing streptavidin-Alexa568 (reaction concentration: 28 µg/ml). The reaction was carried out for 1 h at room temperature. finally, streptavidin coated Dynabeads were opsonized by incubation with IgG (reaction concentration: 1 mg/ml) in 1×PBS for 1 h before live cell experiments.

**pH-MagSensors:** Biotinylated 1 μm Dynabeads were biotinylated following the same procedure described above. Streptavidin-pHrodo Red (reaction concentration: 50 μg/ml) and streptavidin-CF640R (reaction concentration: 50 μg/ml) were then mixed with the biotinylated Dynabeads for 1 h at room temperature. Finally, MagSensors were opsonized with IgG (reaction concentration: 1 mg/ml) in 1×PBS for 1 h before live cell experiments.

**Intracellular pH calibration of pH-MagSensors:** The synthesized MagSensors were incubated with cells seeded on glass coverslips at 37°C for 30 min to allow internalization. Cells were then washed with potassium-rich buffer containing 10 mM nigericin at each desired pH for calibration (pH = 4.60, 5.45, 6.45, and 7.25) (7). Z-stack fluorescence images of phagosomes encapsulating the pH MagSensors were acquired simultaneously in two channels: ex561/em586 nm for pHrodo Red and ex640/em680 nm for CF640R. The calibration experiments were repeated at each individual pH for 16 phagosomes each. The localization and fluorescence intensity of each particle were analyzed using a single-particle tracking MATLAB script as previously described (8). Fluorescence intensity ratios of two channels of each probe at different pH were analyzed using a custom MATLAB algorithm. The average intensity ratio, *I(pHrodoRed)*/*I(ref)*, at each pH was obtained from 16 phagosomes.

**Proteolysis-MagSensors:** Biotinylated 1 μm Dynabeads were washed and resuspended in 1×PBS buffer containing streptavidin-CF640R (reaction concentration: 50 μg/ml). The resulting dynabeads conjugates were incubated with Z-FR-R110 peptide (reaction concentration: 100 μM) in 1xPBS at room temperature for 1 h. Unbound peptides were removed by three times wash with 1xPBS. Finally, MagSensors were opsonized with IgG (reaction concentration: 1 mg/ml) in 1×PBS for 1 h before live cell experiments.

**2.8 μm IgG coated magnetic beads:** The carboxylic groups on 2.8 μm Dynabeads were converted to biotin groups by mixing Dynabeads (reaction concentration: 1 mg/ml), EDC (reaction concentration: 10 mg/ml), and biotin pentylamine (reaction concentration: 1 mM) in MES buffer (50 mM, pH 6.2) at room temperature for 1 h. Subsequently, biotinylated Dynabeads were rinsed and resuspended in 1×PBS buffer containing streptavidin-CF488A (reaction concentration: 20 μg/ml). The reaction was carried out for 1h at room temperature. 1 mg/ml IgG were incubated with MagSensors in 1XPBS for 1 h before live cell imaging.

### 5. Magnetic tweezers setup and force calibration

The magnetic tweezers were set on Nikon Eclipse Ti fluorescence microscope and includes a solenoid and a power supply (Fig.S1*A*). The high permeability HyMu-80 alloy rod (Carpenter Technology, Reading, PA) was inserted into an aluminum bobbin wrapped with 600 turns of copper coil to assemble solenoid (Fig. S1*B*) (9). The diameter of the tip is ≈ 1 μm (Fig. S1*C*). The displacement of magnetic tweezers tip was controlled in live imaging by a manual micromanipulator (Narishige NMN-21). Electromagnetic filed was generated by a power supply (Tekpower, Montclair, CA) which applied up to 5A current through solenoid.

Magnetic force was calibrated as a function of particle-to-solenoid tip distance by measuring the movements of single magnetic particles in calibration medium under magnetic pulling. Carboxylate acid functionalized magnetic particles with the size of 200 nm, 1 μm, and 3 μm were first conjugated with CF640R dye via EDC coupling. The labeled magnetic particles were resuspended in base solution of SYLGARD™ 184 Silicone Elastomer kit with viscosity of 5.1 Pa·s at a concentration of $3.0 \times 10^5$/ml. Low particle concentration was used to avoid aggregations and magnetic interaction between particles (10). High viscosity calibration medium was chosen to decrease the velocity of particles and make the tracking feasible in high time resolution.

The movements of particles of each size under magnetic force were captured with an interval time of 0.2 by a Andor iXon3 EMCCD camera (Andor Technology, Belfast, U.K.). The working current was set to 1 A in all calibration experiments. The magnetic force $F(r)$ generated on each magnetic particles at particle-to-solenoid tip distance $r$ was calculated using Stokes' law:

$$F(r) = 3 \cdot \pi \cdot \eta \cdot d \cdot v \quad (1)$$

where $\eta$ is the viscosity of Sylgard PDMS base, $d$ is the diameter of particle, and $v$ represents the velocity of particle. The relationship between magnetic force $F(r)$ and distance of particle-to-tip $r$ was fitted using equation：

$$F(r) = \frac{F_0}{[\frac{r}{2r_0}+\frac{1}{2}]^{C_F}} \quad (2)$$

in which $F_0$ is a force constant in pN, and $F_{(r0)} = F_0$ .; $r_0$ is a distance constant in μm; $C_F$ is unitless(9).

### 6. Fluorescence microscopy

**6.1 Measurements of Rab5, Rab7, and LAMP1 markers recruitment within phagosomes.** 1 million RAW 264.7 macrophage cells were seeded on glass coverslip in complete DMEM medium for 24 h before the addition of particles. To synchronize particle uptake, pH-MagSensors and Mag-*Toxoplasma* tachyzoites were added to macrophage monolayer expressing Rab5-GFP, Rab7-GFP, or LAMP1-GFP at 5:1 particle-to-cell ratio and centrifuged at 200xg at room temperature for 1 min. Epi-fluorescence imaging was performed using Nikon Eclipse-Ti inverted microscope equipped with a 1.49 N.A. ×100 TIRF objective (Nikon, Tokyo, Japan) and an ORCA-Fusion C14440 digital CMOS camera (Hamamatsu photonics K.K., Shizuoka, Japan) at 37°C. To magnetically manipulate the cell entry of Mag-*Toxoplasma* tachyzoites and MagSensors, magnetic pulling force was applied after the centrifugation when particles were bound with macrophage cell membrane. The force was kept on until the end of live cell imaging. Fluorescence emissions at three wavelengths (ex: 488, 561, and 640 nm; em: 515, 586 and 680 nm) were acquired with a time interval of 7 s in time-lapse multi-channel imaging to record the fluorescence of endocytic markers and the localization of MagSensors.

**6.2 Measurements of phagosome-lysosome fusion.** 0.25 million Raw 264.7 macrophage cells were seeded on glass coverslip in complete DMEM medium for 3 h before the addition of BSA-biotin-Alexa647 (final concentration: 10 μg/ml) for overnight incubation. After that, cells were washed three times with 1xPBS and kept in fresh complete medium to chase BSA-biotin-Alexa647 to lysosomal compartments for 2 h before live cell imaging. Particles were then centrifuged following the procedure described above to synchronize the cell membrane-MagSensor binding. Magnetic pulling force was applied during the cell entry of FRET-MagSensors and the force was kept on until the end of live cell imaging. Epi-fluorescence images were acquired to record FRET emission ($FRET_0$; ex/em 561/680 nm), donor emission ($AF568_{em}$; ex/em 561/586 nm), acceptor emission ($AF647_{em}$; ex/em 660/680 nm) with a time interval of 4 s.

The effect of two types of spectral crosstalk were considered and calibrated in the FRET ratio calculation: (i) the leakage of donor fluorophore Alexa568 emission into acceptor emission channel (680nm) under the excitation of the donor (561 nm) and (ii) the detection of acceptor fluorophore Alexa647 emission in acceptor emission channel (680 nm) under the excitation of donor (561 nm) without the FRET energy transfer. However, we found that the percentages of bleed-through from donor and acceptor fluorophores emission into FRET emission are constant. We then used correction factors $\alpha$ and $\beta$ to cancel out the spectral crosstalk in FRET experiment. The FRET ratio could be calculated using the equation (11)：

$$FRET\ ratio = \frac{FRET_{em}}{AF568_{em}} = \frac{FRET_0 - \alpha \times AF568_{em} - \beta \times AF647_{em}}{AF568_{em}}, \quad (3)$$

Where $\alpha$ was measured based on the FRET emission and donor emission of Alexa568 on FRET-MagSensors that stayed outside of cells, $\beta$ was measured based on the FRET emission and acceptor emission of BSA-Alexa647 in cells without FRET-MagSensors, $FRET_0$ is the measured FRET emission (ex/em 561/680 nm) before cross-talk correction, $FRET_{em}$ is the calibrated FRET emission, $AF568_{em}$ is the donor emission (ex/em 561/586 nm) and $AF647_{em}$ represents the acceptor emission (ex/em 660/680 nm).

**6.3 Measurements of single phagosome acidification.** 1 million of RAW 264.7 macrophage cells were seeded on clean glass coverslip in complete DMEM medium for 24 h before the addition of pH-MagSensors. MagSensors were then centrifuged following the procedure described above to synchronize the cell membrane-MagSensor binding. Magnetic pulling force was applied during the cell entry of pH-MagSensors and the force was kept on until the end of live cell imaging. Fluorescence emissions at one wavelength (ex: 561 nm; em: 586 nm) and two wavelengths (ex: 561 and 640 nm; em: 586 and 680 nm) were acquired with a time interval of 7 s in time-lapse multi-channel imaging to record the fluorescence of Mag-*Toxoplasma* tachyzoites and pH-MagSensors, respectively. At the end of each experiment, cells were washed and incubated in pH 4.6 potassium-rich buffer containing 10 mM nigericin for 5 min. Then, the emissions of mag-*Toxoplasma* and pH-MagSensors were recorded again.

**6.4 Measurements of single phagosome proteolytic activity.** 1 million actin-RFP expressing macrophage cells were seeded on clean glass coverslip in complete DMEM medium for 24 h before the addition of proteolysis-MagSensors. MagSensors were then centrifuged following the procedure described above to synchronize the cell membrane-MagSensor binding. Magnetic pulling force was applied during the cell entry of proteolysis-MagSensors and the force was kept on until the end of live cell imaging. Time-lapse multi-channel epi-fluorescence images were acquired to record emission of rhodamine 110 (ex/em 488/515 nm), actin-RFP (ex/em 561/586 nm) and CF640R (ex/em 640/680 nm) with a time interval of 7 s.

**6.5 Measurements of Fc receptor clustering during MagSensor cell entry.** Anti-mouse CD16/32 Fab fragments were prepared using Pierce Fab preparation kit following manufacturer's instruction. The generated Fab fragments were further conjugated with CF 640R NHS ester. In brief, Fab fragments (reaction concentration: 14 µg/ml) and CF640R (reaction concentration: 7 µg/ml) were mixed in sodium bicarbonate buffer (0.1 M, pH 8.25) for 3 h. Free dyes were removed by centrifugal filtration using Amicon Ultra filters (30K). To label FcγII/FcγIII receptors on cell membrane, confluent macrophage cells were incubated in Ringer's imaging buffer containing 2.3 µg/ml fluorescent Fab for 10 min. Subsequently, cells were washed three times with Ringer's imaging buffer, and 2.8 µm IgG coated MagSensors conjugated with streptavidin-CF488A were added. MagSensors were then centrifuged following the procedure described above to synchronize the cell membrane-MagSensor binding. Magnetic pulling force was applied during the cell entry of 2.8 µm MagSensors. Time-lapse multi-channel epi-fluorescence images were acquired to record emission of CF488A (ex/em 488/515 nm and CF640R (ex/em 640/680 nm) with a time interval of 7 s.

**6.6 Measurements of CD45 exclusion during MagSensor cell entry.** CD45 labeling on macrophage plasma membrane was carried out following a previously reported protocol (12). Briefly, cells were incubated in Ringer's imaging buffer containing Alexa647 conjugated rat anti-mouse CD45 (final concentration: 500 ng/ml) for 10 min in room temperature. After that, cells were washed with Ringer's imaging buffer and 2.8 µm MagSensors conjugated with streptavidin-CF488A were added. MagSensors were then centrifuged following the procedure described above to synchronize the cell membrane-MagSensor binding. Magnetic pulling force was applied during the cell entry of 2.8 µm MagSensors. Time-lapse multi-channel epi-fluorescence images were acquired to record emission of CF488A (ex/em 488/515 nm and Alexa647 (ex/em 640/680 nm) with a time interval of 7 s.

**6.7 Measurements of actin filaments assembly near nascent phagosomes.** 1 million actin-EGFP expressing macrophage cells were seeded on glass coverslip in complete DMEM medium for 24 h before the addition of pH-MagSensors. MagSensors were then centrifuged following the procedure described above to synchronize the cell membrane-MagSensor binding. Magnetic pulling force was applied during the cell entry of pH-MagSensors and the force was kept on until the end of live cell imaging. Time-lapse multi-channel epi-fluorescence images were acquired to record emission of actin-GFP (ex/em 488/515 nm), pHrodo Red (ex/em 561/586 nm) and CF640R (ex/em 640/680 nm) with a time interval of 7 s.

**6.8 Measurements of Rab5, Rab7, and LAMP1 recruitment within phagosomes or parasitophorous vacuole (PV) in fixed cells.** RAW 264.7 macrophage cells transfected Rab5-RFP, Rab7-RFP, or LAMP1-RFP following the procedure described above. Live and heat-killed (CF640R NHS labeled) *Toxoplasma* tachyzoites were added to transfected cell monolayer at 10:1 parasite-to-cell ratio in complete medium. The cell chambers were incubated at 37°C with 5% $CO_2$ and were subsequently fixed using 2% paraformaldehyde (PFA) after 7.5, 15, and 40 min of incubation.

**6.9 Macrophage plasma membrane GM-1 labeling and imaging.** Macropahge cell plasma membrane GM-1 was labeled by incubating confluent macrophage monolayer in Ringer's imaging buffer containing Cholera Toxin Subunit B CF488A (final concentration: 500 ng/ml) for 10 min at room temperature. After that, cells were rinsed three times with Ringer's imaging buffer. Alexa568 labeled Mag-*Toxoplasma* tachyzoites were centrifuged to synchronize the cell membrane-parasite binding. Magnetic pulling force was applied during the cell entry of Mag-*Toxoplasma* tachyzoites. Time-lapse multi-channel epi-fluorescence images were acquired to record emission of CF488A (ex/em 488/515 nm and Alexa568 (ex/em 561/586 nm) with a time interval of 7 s.

**6.10 Trypan blue quenching assay.** Trypan blue has been shown to quench Alexa488 (13), Alexa647 (14), and QDs with 605 nm (15) emission. To distinguish internalized MagSensors from those located extracellularly, trypan blue solution was added at the end of imaging (final concentration: 4.6 µM). After 10 min of incubation, cell samples were imaged and the fluorescence intensity of MagSensors was used to distinguish internalized particles from extracellular ones.

**6.11 Imaging the plasma membrane encapsulation of MagSensors.** RAW 264.7 macrophages were transfected with the plasma membrane-RFP as a membrane marker following the protocol mentioned above. CF640R coated 1 µm MagSensors were mixed with macrophages at a particle-to-cell ratio of 5:1 at 37°C for 5 min before image acquisition. The internalization of MagSensors were manipulated either with or without magnetic manipulation. Once the MagSensors were inside the cell, confocal fluorescence images were acquired using a Nikon Eclipse-Ti inverted microscope expanded with a rescan confocal microscopy (RCM) unit (Confocal.nl, Amsterdam, The Netherlands) (16) and Hamamatsu ORCA-Fusion C14440 digital CMOS camera. The 561 nm and 641 nm laser were employed to excite the plasma membrane-RFP and CF640R on MagSensors, respectively.

**6.12 *Toxoplasma* SAG-1 labeling.** The heated-killed *Toxoplasma* tachyzoites were sequentially labeled with 200 nm magnetic nanoparticles (CF488A labeled) and pHrodo Red STP following the methods described above. The prepared parasites were spun down on glass coverslip and maintained in PBS. *Toxoplasma* gondii major surface antigen (SAG1) was labeled with SAG1 Monoclonal Antibody (D61S) (final concentration: 2 µg/ml) for 15 min. Free primary antibody in solution was removed with 1xPBS washing for three times. Next, parasites were stained with Alexa647 conjugated chicken anti-mouse IgG secondary antibody (final concentration: 5 µg/ml) for 15 min. Subsequently, *Toxoplasma* tachyzoites were rinsed and imaged to record emission of CF488A (ex/em 488/515 nm, pHrodo Red (ex/em 561/586 nm), and Alexa647 (ex/em 640/680 nm).

**6.13 Stochastic optical reconstruction microscopy (STORM).** STORM imaging of CD45 molecules on macrophage plasma membranes were performed using a previously reported protocol (17). CD45 labeled macrophages were first fixed with 2% paraformaldehyde and then kept in freshly made imaging buffer containing oxygen scavengers and reducing agents (100 mM MEA and 1×GLOX solution in Buffer B containing 50 mM Tris-HCl (pH 8.0), 10 mM NaCl and 10% w/v glucose) and imaged at room temperature. STORM imaging was performed in TIRF mode using the Nikon Ti2 Eclipse microscope equipped with a TIRF × 100 oil-immersion objective (NA 1.49), an Andor iXon3 EMCCD camera with a pixel size of 105 nm and a perfect focus system (PFS). Image acquisition of AF647 labeled primary CD45 antibodies were acquired under the excitation of 637 nm laser at a power density of 10 kW/cm$^2$. Raw Images were obtained with an exposure time of 10 ms and 10000 consecutive frames. Fluorophores were stochastically activated using a 405-nm diode laser at the power density of 0.5 kW/cm$^2$.

**7. Scanning electron microscopy (SEM)**

Preparation of *Toxoplasma tachyzoites* SEM samples were carried out following a previously published protocol (18). In brief, heat-killed *Toxoplasma* tachyzoites with or without conjugation of 200 nm magnetic particles were deposited on glass coverslips at proper density. Parasites were washed three times with 0.1 M sodium phosphate buffer (0.07 M $Na_2HPO_4$ and 0.03 M $NaH_2PO_4$, pH 7.2) and subsequently fixed in Karnovsky's fixative (containing 2% (w/v) paraformaldehyde and 2.5% (v/v) glutaraldehyde in 0.1 M sodium phosphate buffer) at room temperature for 30 min. After being rinsed three times with sodium phosphate buffer (0.1 M, pH 7.2), the *Toxoplasma* tachyzoites were postfixed with 1% (v/v) osmium tetroxide (aqueous solution, Electron Microscopy Sciences) in sodium phosphate buffer (0.1 M, pH 7.2) at room temperature for 1 h. Next, samples were washed with Milli-Q water twice for 10 min each. Dehydration of tachyzoites were then performed using a series of ethanol solution with increasing volume fraction (35, 50, 76 and 95 volume %) for 10 min per wash and rinsed in pure ethanol twice for 10 min each. The hexamethyldisilazane

(HMDS; Electron Microscopy Sciences) was then used to immerse *Toxoplasma* tachyzoites twice for 10 min each. Finally, the HMDS was decanted, and samples were air-dried overnight in a desiccator. Next day, the dried samples were immobilized onto a SEM sample stub and sputter-coated with 4 nm Au/Pd alloy, and then imaged with FEI Quanta 600 SEM.

### 8. Images analysis

**8.1 Single-*Toxoplasma* tachyzoites localization and intensity determination.** The localization and pHrodo Red fluorescence intensity of heat-killed *Toxoplasma* tachyzoites were analyzed using a single-particle tracking MATLAB script as previously described (8).

**8.2 Analysis of the recruitment of Rab5 and LAMP1 within single phagosomes containing heat-killed *Toxoplasma tachyzoites* in live cell imaging.** The accumulation of Rab5 and LAMP1 on single phagosomes were analyzed using custom MTALAB algorithm that extract pixel intensities for Rab5 or LAMP1 fluorescence using *Toxoplasma* fluorescence-based masks. We use the analysis of Rab5-GFP over phagosomes as an example. Briefly, tiff file image stacks containing both *Toxoplasma* tachyzoites (red channel) and Rab5-GFP (green channel) fluorescence channels were compiled into MATLAB, and frames from the red channel were used to sequentially generate dynamic ROI masks for collecting pixels of interest within the green channel. To each red channel frame, a gaussian image filter was applied followed by background removal by subtraction of a user defined percentage of the mean unique pixel intensities. After setting negative pixel values to 0, rough *Toxoplasma* boundaries were constructed using the built-in MATLAB edge detection function with the Laplacian of Gaussian based zero-crossing algorithm. Morphological closing was then applied to binary images of constructed edges using the bwmorph MATLAB function to patch any noncontinuous edge sections. Edges corresponding to distinct *Toxoplasma* were then grouped using the bwboundaries MATLAB function, followed by user defined edge thickness padding to control final ROI mask sizes. Concurrently, peripheral background ROI masks are generated using the bwmorph "thicken" option. Finally, intensities of green channel pixels within the edge ROI coordinates were averaged and divide by the averaged green channel pixel intensities from the background ROI mask.

**8.3 Single-MagSensor Localization and Intensity Determination.** The localization and fluorescence intensity of MagSensors were analyzed using a single-particle tracking MATLAB script as previously described (8). Emission of pHrodo Red, CF640R, Rhodamine 110 were calculated as the background subtracted and integrated pixel intensities within in 2 μm from the localized centroid of particles. $FRET_0$, $AF568_{em}$ and $AF647_{em}$ in single-phagosome FRET-fusion experiment were determined with the same procedure.

**8.4 Analysis of the recruitment of actin filaments, Rab5, Rab7, and LAMP1 within single phagosomes containing MagSensors.** The accumulation of endocytic markers and actin filaments around single phagosomes were analyzed using MATLAB. We use the analysis of Rab5-GFP over phagosomes as an example. First, phagosomes containing MagSensors were individually outlined by circles (1 μm radius) using the single-particle tracking Matlab script mentioned above (8). Second, integrated fluorescence intensities of Rab5-GFP from all pixels inside each outline circle were obtained. Third, integrated intensities of all pixels along the perimeter of the circles with an outer radius of 1.5 μm and an inner radius of 2 μm were obtained. Finally, normalized Rab5-GFP intensities were calculate as the ratio of the integrated intensity from pixels inside the particle outline circles to that along the perimeter.

**8.5 Analysis of Fcγ receptor clustering and CD45 exclusion during MagSensor cell entry.** The fluorescence intensities of labeled Fcγ receptors and CD45 were analyzed in Image J. We use the analysis of Fcγ clustering during cell entry as an example. A line was drawn at the MagSensor-cell membrane interface and the fluorescence intensities of CF488A on MagSensors and CF640R on Fab fragments were plotted at each time frame. The intensity line plots at which frame that had the highest normalized intensity value for Fab and lowest normalized intensity value for CD45 at the particle—cell contact area were chosen as the intensity plots for the entire internalization event. This analysis method also applies to the measurement of CF45 exclusion during mag-*Toxoplasma* cell entry.

**8.6 STORM image analysis.** TIRF-STORM images were reconstructed using ImageJ plugin ThunderSTORM (19). The centroids in each frame were determined by fitting their point-spread functions (PSF) using the least-square Gaussian fitting method. Single molecules in high spatial densities were resolved by a multiple emitter fitting analysis (MFA) (20). Several filters were applied to remove noise in raw images.

i. Dim molecules were removed by applying a threshold of $N > \alpha b$, where $N$ represents the number of photons detected from a given molecule; b is the standard deviation of the signal in fitting region and the sensitivity factor $\alpha$ was set as 30 in this study (21).

ii. To eliminate repeated localization of the same single molecule in one frame, locations detected that were within proximity of 20-30 nm were grouped together and only molecule in each group with smallest localization inaccuracy was used as signal. Localization inaccuracy ($\delta_i$) was defined and calculated as

$$\delta_i = \sqrt{+\frac{\sigma_i^2 + a^2/12}{N} + \frac{8\pi\sigma_i^4 b^2}{a^2 N^2}}, \quad (4)$$

where $N$ represents the number of photons collected from a given molecule, $\sigma_i$is the fitted standard deviation of the PSFs of the detected molecules in either the x and y direction, $a$ is the pixel size of the CCD camera (105 nm) and $b$ is the standard deviation of background.

iii. To eliminate multiple blinking from a single molecule in sequential frames, locations appeared within the single pixel size of CCD camera in five consecutive frames were merged and fitted to one localization. The newly merged localizations were generated using averaged coordinates of pre-merged localizations. The photon numbers and the background signal level of the new localizations were determined as the sum values of those parameters from the pre-merged localizations. The localization inaccuracy of a merged molecule was calculated by plugging the newly acquired values into equation 4.

### 9. Simulation methods

We utilized a computational framework in which the pathogen surface was in close proximity to a portion of the macrophage surface. Surface proteins were represented as particles that could diffuse; receptors on the macrophage surface could reversibly bind immobile ligands on the pathogen surface. Dynamics of the particles were governed by a stochastic reaction-diffusion process. The shape of the membrane was characterized by a time-dependent Ginzburg-Landau (TDGL) model that accounted for the membrane bending energy and protein deformation energies when surface proteins were located in regions inconsistent with their natural length. The total magnitude of the force by which the pathogen surface was pushed against the macrophage surface was systematically varied. We utilized a hybrid method coupling the Gillespie algorithm for particle dynamics with a finite difference method to update the membrane shape in accordance with TDGL dynamics.

**9.1 Energetics of the membrane.** We considered a $2\ \mu\text{m} \times 2\ \mu\text{m}$ region of the macrophage surface that was parameterized by Cartesian coordinates $x$ and $y$. The position-dependent distance between the macrophage and pathogen surfaces was given by the function $z(x,y)$. The effective energy of the membrane and associated particles was given by (22):

$$E[z, C_{\text{CD45}}, C_{\text{bond}}] = \frac{1}{2}\iint dx\, dy\ \left[\kappa(\nabla^2 z_{\text{m}})^2 + k_{\text{CD45}}\, C_{\text{CD45}}\, (z - z_{\text{CD45}})^2\, H(z_{\text{CD45}} - z) + k_{\text{bond}}\, C_{\text{bond}}\, (z - z_{\text{bond}})^2 + k_{\text{mag}} C_{\text{mag}} (z - z_{\text{mag}})^2\right],$$

where $z_{\text{m}}(x,y)$ gives the shape of the macrophage surface, $C_{\text{CD45}}(x,y)$ is the concentration of CD45, $C_{\text{bond}}(x,y)$ is the concentration of receptor-ligand bonds, $\kappa$ is the bending rigidity of the membrane, $k_{\text{CD45}}$ and $k_{\text{bond}}$ are the spring stiffnesses associated with deforming a molecule relative to its natural length, and $H$ denotes the Heaviside step function. The first term in the integral accounts for energy contributions from local curvature of the membrane; $z_{\mathrm{m}}(x, y)$ was defined by subtracting the pathogen surface from $z(x, y)$. The Heaviside function imposed the condition that there was no energetic penalty when CD45 molecules were located at locations with a larger separation than the length of the molecule. The concentrations were defined in terms of the specific locations of particles, $C_i(\boldsymbol{r}) = \sum_{j=1}^{N_i} \delta(\boldsymbol{r} - \boldsymbol{r}_j^{(i)})$, where $\delta$ denotes the Dirac delta function and $\boldsymbol{r}_j^{(i)}$ is the position of particle $j$ of type $i$ (= CD45 or bond). The number of bonds dynamically changed due to receptor-ligand binding and dissociation. The final term was used to impose an external force akin to the magnetic-field imposed force in experiments. Here $C_{\mathrm{mag}}$ was uniform across the surface of the pathogen; the term can be regarded as applying a force distributed through space that pushes the two surfaces together. The value of $k_{\mathrm{mag}} C_{\mathrm{mag}}$ was systematically varied to change the magnitude of the force applied to the pathogen.

**9.2 Stochastic reaction-diffusion dynamics of proteins.** For numerical implementation, we discretized the membrane into a square lattice. The lattice spacing was $\Delta x = 13\ \mathrm{nm}$, and each lattice site could contain at most one particle from the same membrane. Particles diffused by means of a stochastic hopping process between neighboring lattice sites. The intrinsic hopping rate to a nearest neighbor lattice site was given in terms of the diffusion coefficient ($D$) by $\gamma = D/\Delta x^2$. To account for the fact that molecules are less likely to diffuse into energetically unfavorable regions, the rate was modified according to the method of Grima and Newman such that the rate of hopping from site $k$ to neighboring site $l$ was $\gamma_{k \to l} = \gamma \exp(-(E_l - E_k)/2k_{\mathrm{B}}T)$ (23).

Receptors on the macrophage surface could bind to ligands on the pathogen surface when sufficiently close. In the model, binding could occur when receptors and ligands occupied the same lattice site and the distance between surfaces was within $10\ \mathrm{nm}$ of the natural bond length, $z_{\mathrm{bond}}$. The binding rate was given by $k_{\mathrm{on}}(z) = k_{\mathrm{on}}^0 \exp(-(z - z_{\mathrm{bond}})^2/2\sigma^2)$, where $\sigma = 5\ \mathrm{nm}$ is a characteristic decay length. Bonds dissociated stochastically, with the off rate modified by the tension on the bond according to a slip-bond model, $k_{\mathrm{off}}(z) = k_{\mathrm{off}}^0 \exp(k_{\mathrm{bond}}\,(z - z_{\mathrm{bond}})/f_0)$, where $f_0 = 5.5\ \mathrm{pN}$ is a characteristic force.

Given the location of all particles and the distance profile $z(x, y)$, we calculated the rates associated with all hopping, binding, and unbinding reactions. A time step and specific event were chosen using the Gillespie algorithm (24). The state of the system was then updated.

**9.3 Dynamics of the membrane.** After each step of the Gillespie algorithm, the shape of the membrane was updated in accordance with the TDGL framework (25):

$$\frac{\partial z}{\partial t} = -M \frac{\delta E}{\delta z} + \zeta$$

$$= -M\left[\kappa\, \nabla^4 z + k_{\mathrm{CD45}}\, C_{\mathrm{CD45}}\,(z - z_{\mathrm{CD45}}) H(z_{\mathrm{CD45}} - z) + k_{\mathrm{bond}}\, C_{\mathrm{bond}}\,(z - z_{\mathrm{bond}}) + k_{\mathrm{mag}} C_{\mathrm{mag}}(z - z_{\mathrm{mag}})\right] + \zeta$$

Here, $M$ is a phenomenological constant associated with membrane relaxation and $\zeta$ is a random variable satisfying the fluctuation dissipation theorem (i.e., thermal noise). For each update in the Gillespie algorithm, we updated the state of the membrane by solving the governing differential equation using a forward finite difference scheme for the time derivative and a central finite difference scheme for the spatial derivatives. The time step varied according to the time interval determined by the Gillespie algorithm in the previous step. After updating the membrane shape, we returned to the previous Gillespie step and updated the rates to reflect the new state of the system. This iterative process was repeated until the desired time was reached.

Ten replicates were simulated for each set of conditions. The system was initialized by placing particles uniformly at random on the surface subject to single occupancy of lattice sites. All receptors started unbound. The surfaces were initialized at a distance of $20\ \mathrm{nm}$ at the center of the domain, which was the point at which they were closest. The pathogen surface was approximated as a rigid portion of a sphere with a radius of $3\ \mu\mathrm{m}$ curving away from the macrophage surface. Parameters used in the model (22) are included in Table S1.

**Table S1.** Parameters used in simulations.

| | |
|---|---|
| Concentration of receptors | 325 particles/mm$^2$ |
| Concentration of ligands | 800 particles/mm$^2$ |
| Concentration of CD45 | 54 particles/mm$^2$ |
| Diffusion coefficient of receptors | 0.01 mm$^2$/s |
| Diffusion coefficient of CD45 | 0.021 mm$^2$/s |
| Length of receptor-ligand bond ($z_{\mathrm{bond}}$) | 13 nm |
| Length of CD45 ($z_{\mathrm{CD45}}$) | 40 nm |
| Receptor-ligand bond stiffness ($k_{\mathrm{bond}}$) | 1.0 pN/nm |
| CD45 stiffness ($k_{\mathrm{CD45}}$) | 0.1 pN/nm |
| Bending rigidity of membrane ($\kappa$) | 40 $k_B T$ |
| Receptor-ligand binding rate ($k_{\mathrm{on}}$) | 57 nm$^2$/s |
| Receptor-ligand off rate ($k_{\mathrm{off}}$) | 0.2 s$^{-1}$ |

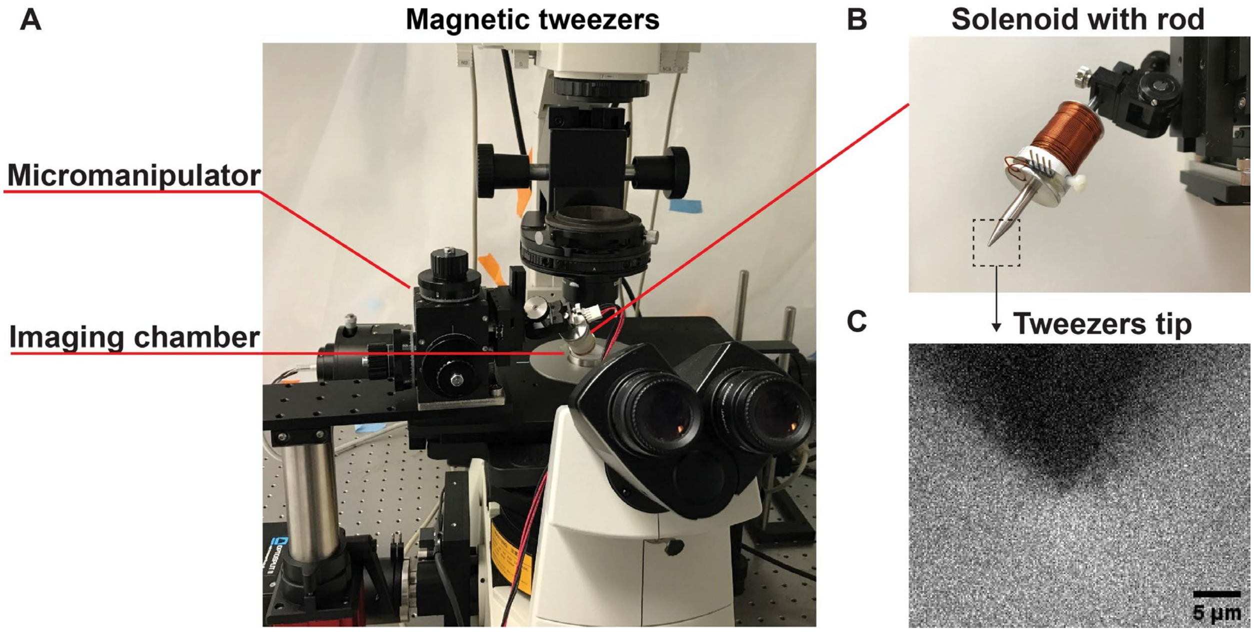


**Fig. S1.** Magnetic tweezers setup. (*A*) A picture showing the magnetic tweezers setup installed on a Nikon Eclipse-Ti inverted microscope. The displacement of magnetic tweezers tip was controlled by a micromanipulator. (*B*) A picture showing the solenoid with an inserted alloy rod. (*C*) Bright field image showing the magnetic tweezers tip.

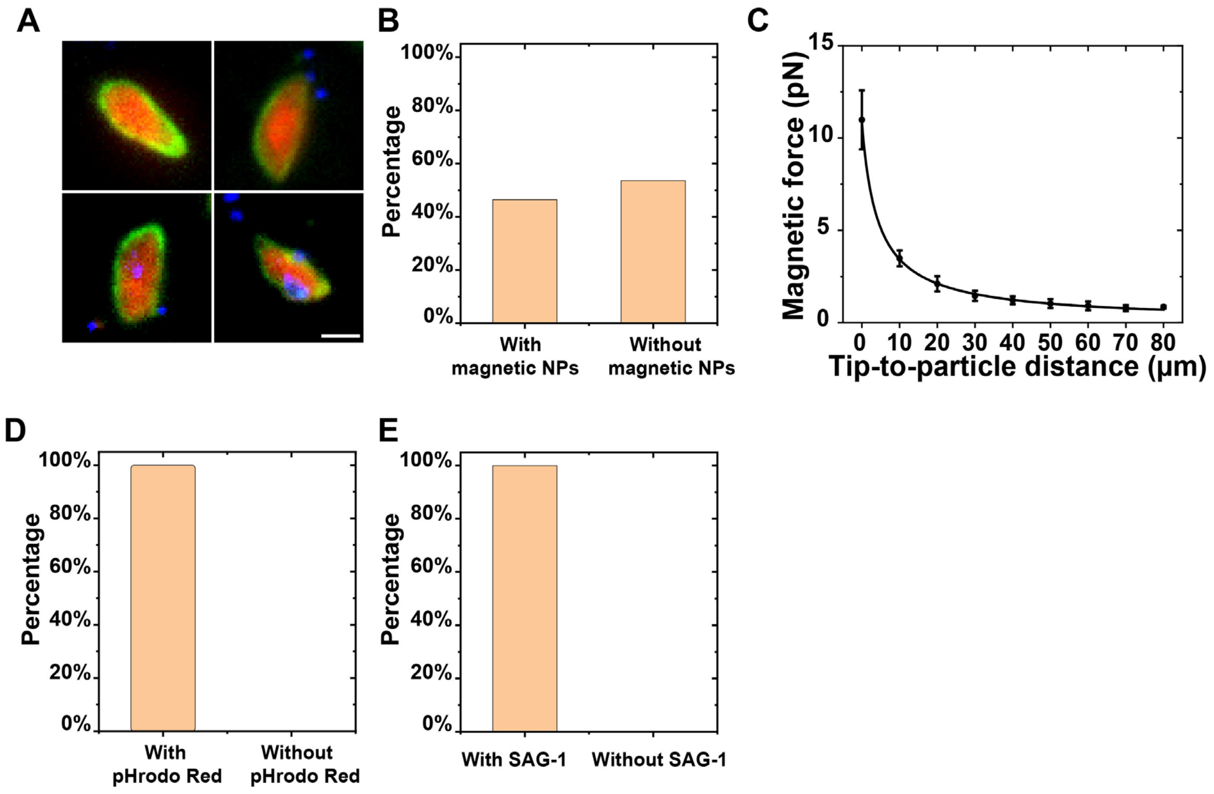


**Fig. S2.** Characterizations of mag-*Toxoplasma*. (*A*) Epi-fluorescence images show the immunostained SAG-1 on the surface of mag-*Toxoplasma*. Magnetic particles are shown in blue, pHrodo Red in red, and SAG-1 in green. Scale bar, 2 μm. (*B*) Bar graph showing the percentage of Toxoplasma tachyzoites labeled with magnetic nanoparticles. Statistically, 46.4% of the heat-killed *Toxoplasma* were labeled with at least one magnetic nanoparticle. (*C*) Calibration plots showing the magnetic force exerted on single 200 nm magnetic nanoparticles as a function of the distance from the bead to the tip of magnetic tweezers solenoid. Error bars are standard deviation from 6 samples. (*D*) Bar graph showing the percentage of *Toxoplasma* labeled with pH sensitive dye pHrodo Red. All tachyzoites were successfully labeled with pHrodo Red. (*E*) Bar graph showing the percentage of *Toxoplasma* labeled with anti-SAG 1 antibody. All tachyzoites were SAG-1 positive after magnetic particle conjugation and pHrodo Red labeling.

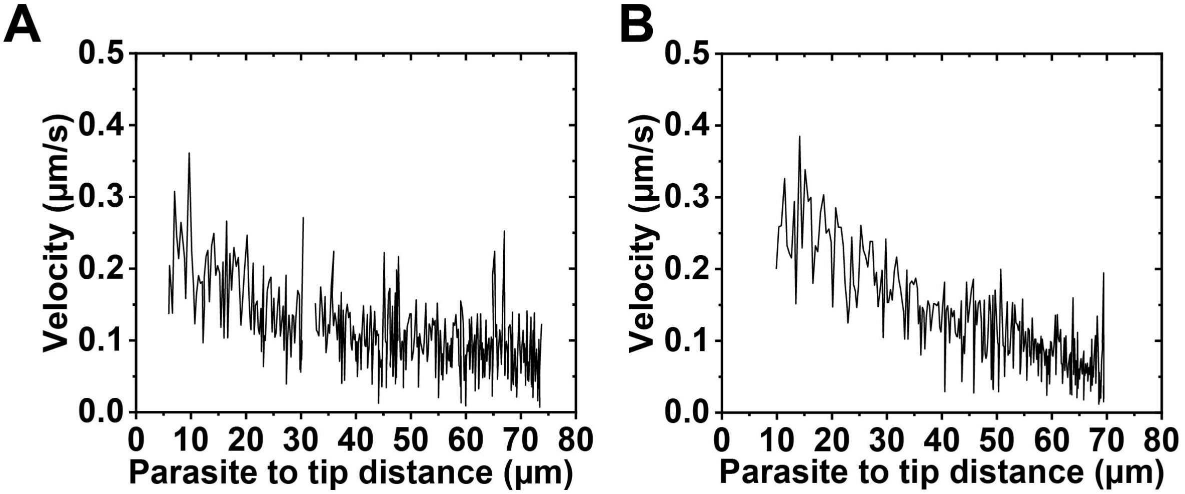


**Fig. S3.** Calibration plots showing the instantaneous velocity of two mag-*T*oxoplasma tachyzoites in PDMS base calibration medium under magnetic pulling, plotted as a function of the distance from the center of the Toxoplasma of interest to the tip of magnetic tweezers.

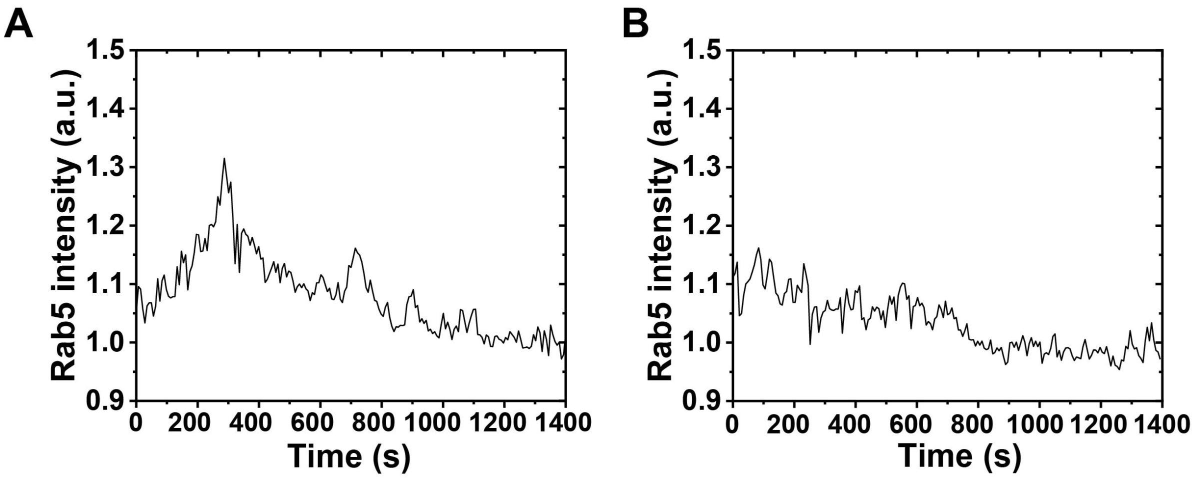


**Fig. S4.** Line plots showing Rab5 recruitment on two phagosomes containing mag-*Toxoplasma* without force (*A*) and with magnetic force applied (*B*).

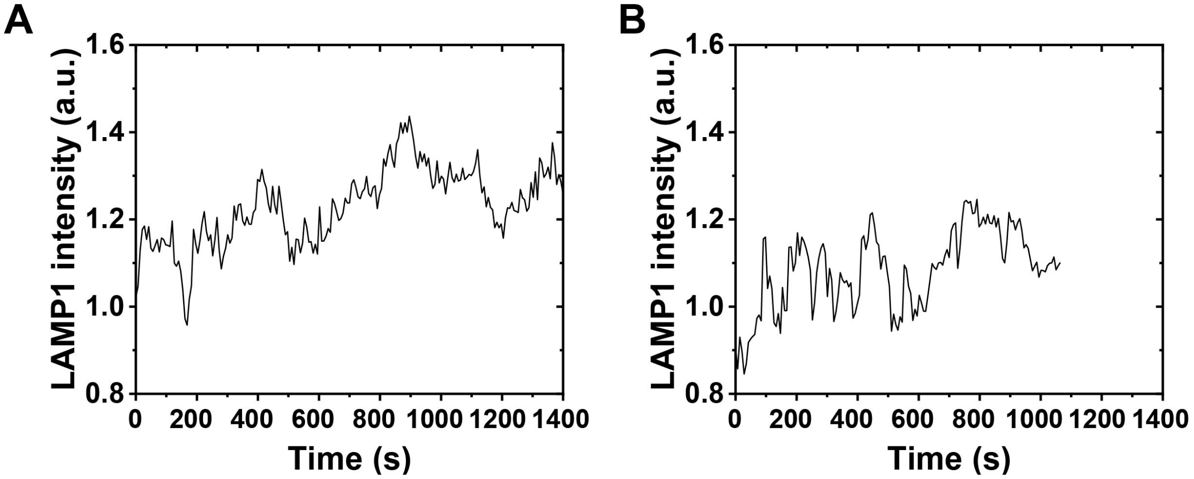


**Fig. S5.** Line plots showing LAMP1-GFP recruitment on phagosomes containing mag-*Toxoplasma* without magnetic force (*A*) and with magnetic force applied during parasite internalization (*B*).

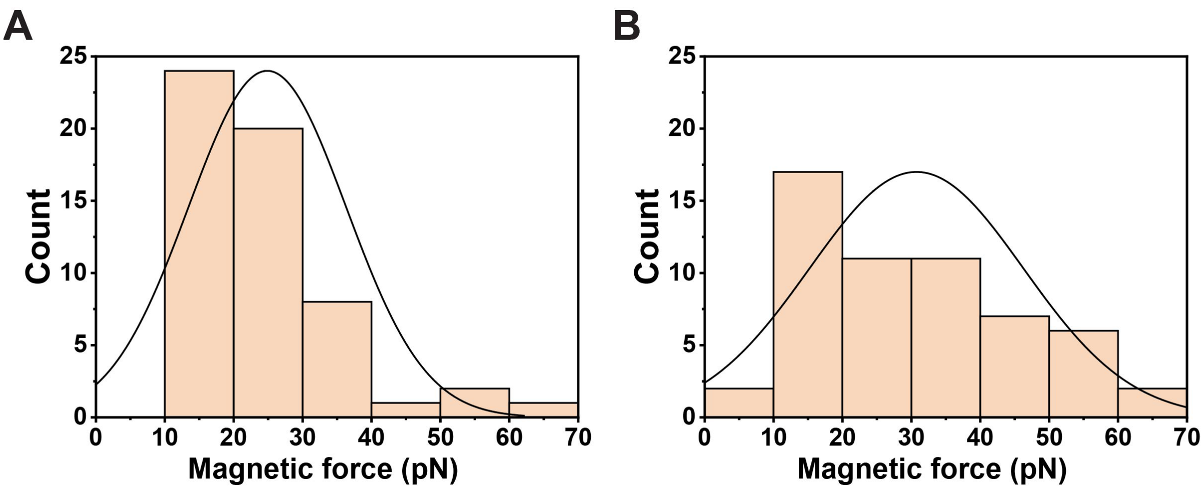


**Fig. S6.** (*A*) Distribution of the estimated magnetic pulling force applied on individual 1-μm magnetic beads at the begging of magnetic manipulation. (*B*) Distribution of the estimated magnetic pulling force exerted on individual 1-μm magnetic beads at the end of imaging. The average pulling force was 25 pN at the beginning of magnetic manipulation and 31 pN at the end of imaging as the bead moved closer to tip.

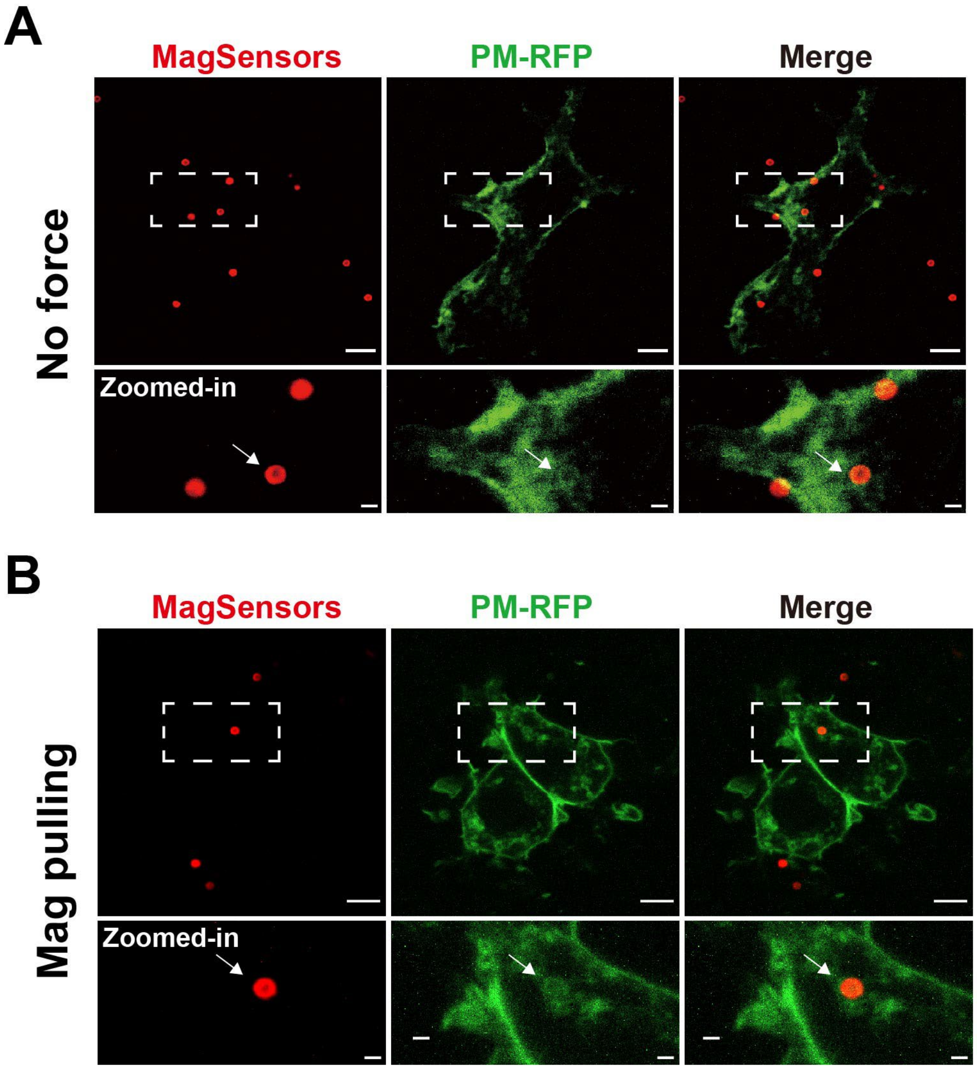


**Fig. S7.** Confirmation of the membrane encapsulation of MagSensors by plasma membrane without force (*A*) and with magnetic force applied. (*B*). The fluorescence of CF640R coated on MagSensors is shown in red and plasma membrane (PM)-RFP is shown green. Scale bars, 5 µm in zoomed-out images, 1 µm in zoomed-in images.

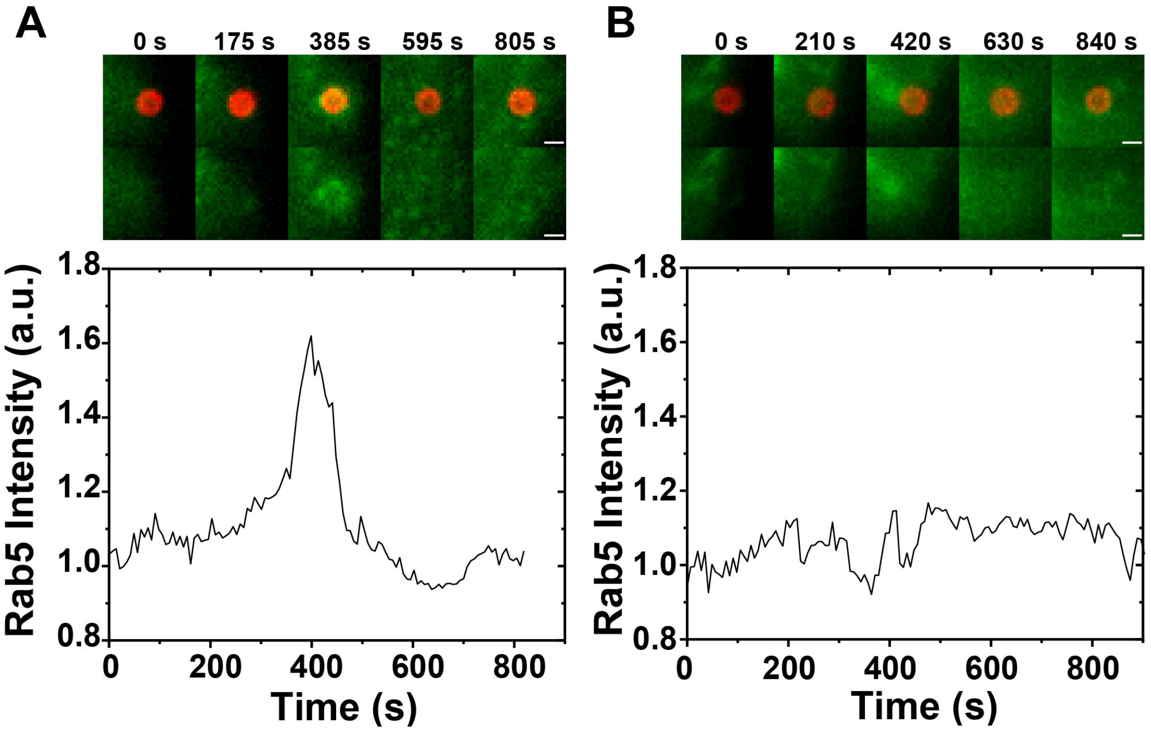


**Fig. S8.** Fluorescence images and line plots showing the recruitment of Rab5 on two phagosomes with no force applied (*A*) and with magnetic pulling (*B*) during their cell entry. In fluorescence images, Rab5-GFP is shown in green and pHrodo red on MagSensors shown in red. Scale bars, 1 μm.

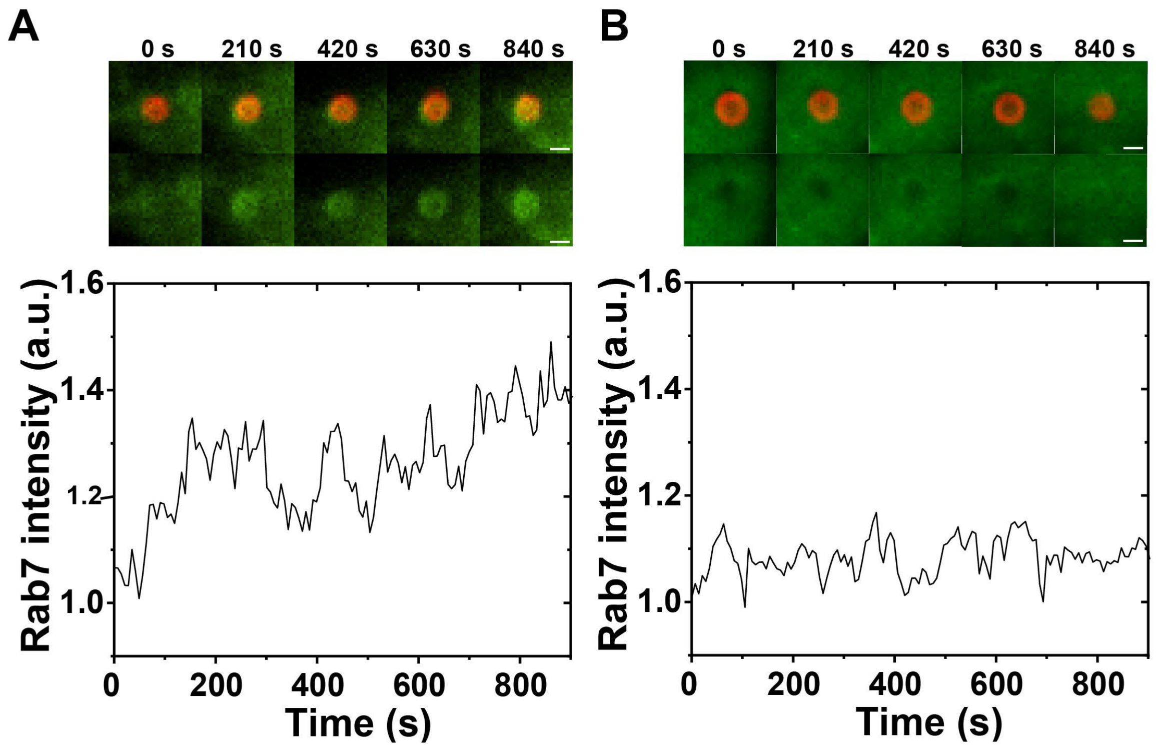


**Fig. S9.** Fluorescence images and line plots showing the recruitment of Rab7 on two phagosomes with no force applied (*A*) and with magnetic pulling (*B*) during their cell entry. In fluorescence images, Rab7-GFP is shown in green and pHrodo red on MagSensors shown in red. Scale bars, 1 μm.

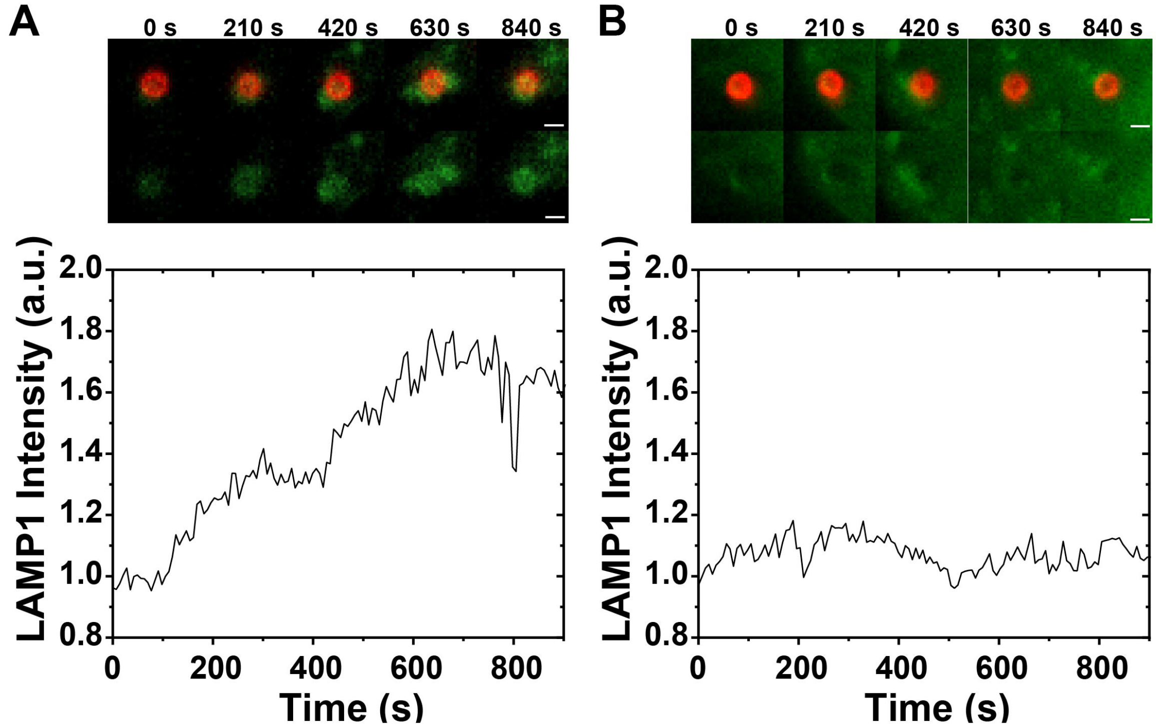


**Fig. S10.** Fluorescence images and line plots showing the recruitment of LAMP1 on two phagosomes with no force applied (*A*) and with magnetic pulling (*B*) during their cell entry. In fluorescence images, LAMP1-GFP is shown in green and pHrodo red on MagSensors shown in red. Scale bars, 1 μm.

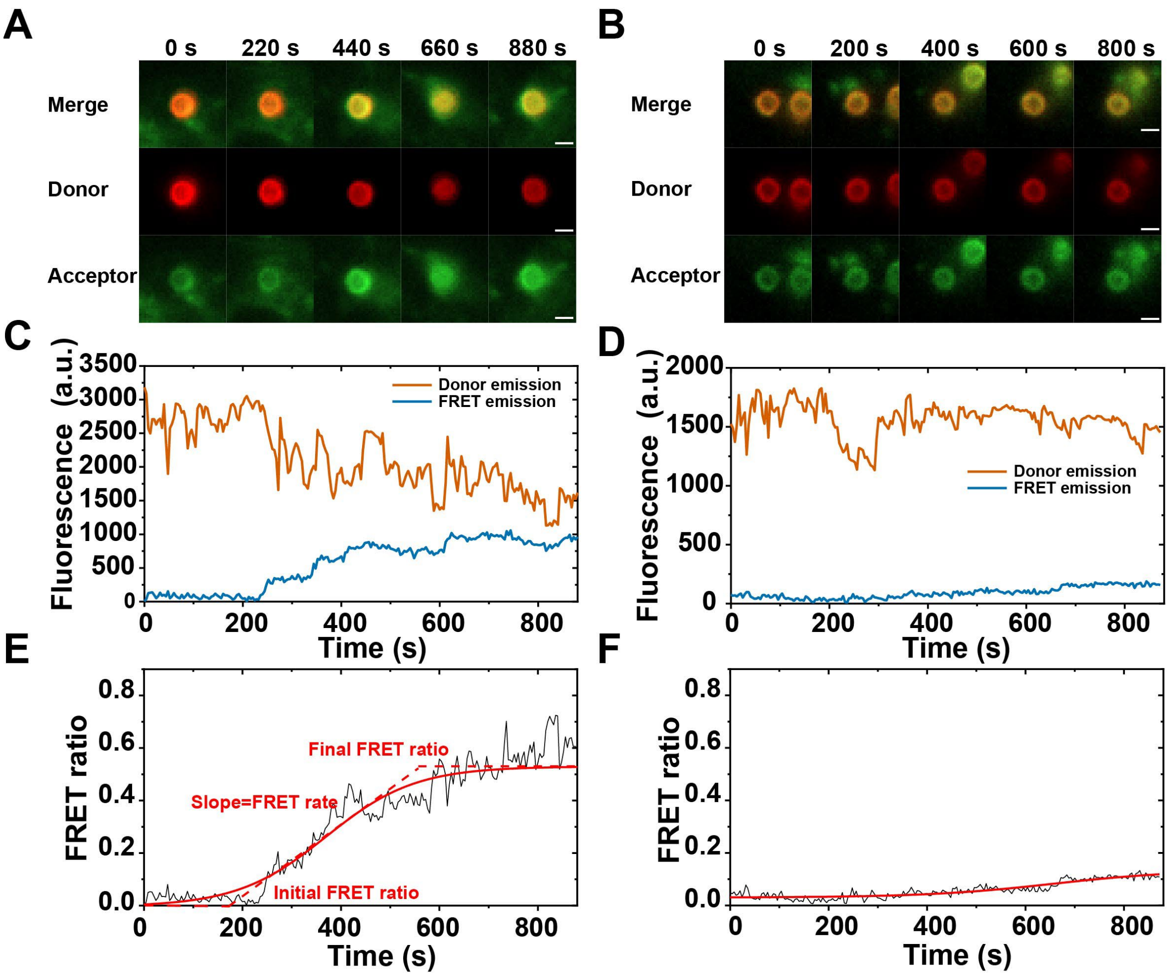


**Fig. S11.** Fluorescence images and line plots showing phagosome-lysosome fusion monitored by FRET microscopy without force (A, C, E) and with magnetic force applied (B, D, F). (*A* and *B*) Fluorescence images showing the change of donor emission (red) and FRET emission (green) from two MagSensor-containing phagosomes formed without (A) or with magnetic force applied (B). Scale bars, 1 µm. (*C* and *D*) The corresponding line plots showing donor and FRET fluorescence intensity change as a function of time of the two sample phagosomes in *A* and *B*. (*E* and *F*) The corresponding line plots showing FRET ratio change as a function of time of the two sample phagosomes in *A* and *B*. The FRET ratio curve is fitted with sigmoidal-Boltzmann function. FRET rate was determined as the slope at $FRET\ ratio = \frac{FRET\ ratio_{initial} + FRET\ ratio_{final}}{2}$.

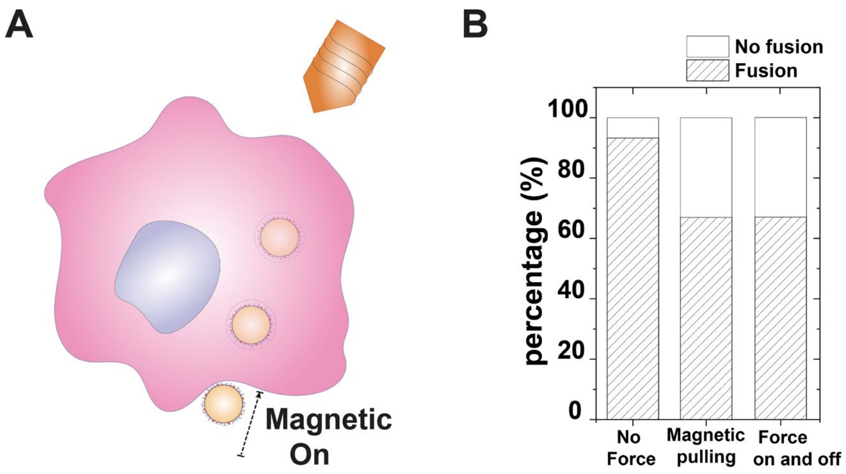


**Fig. S12.** Phagosome responses upon turning off magnetic manipulation after cell entry. (*A*) Schematic illustration of the experimental design in which the magnetic force was applied during cell entry but turned off after internalization. (*B*) Statistic result showing the percentage of phagosome-lysosome fusion under different modes of magnetic manipulation. From left to right: 14 out of 15, 10 out of 15, and 10 out of 15 phagosomes underwent fusion with lysosomes in the case of no magnetic force applied, pulling during cell entry and was kept on after, and pulling only during internalization, respectively.

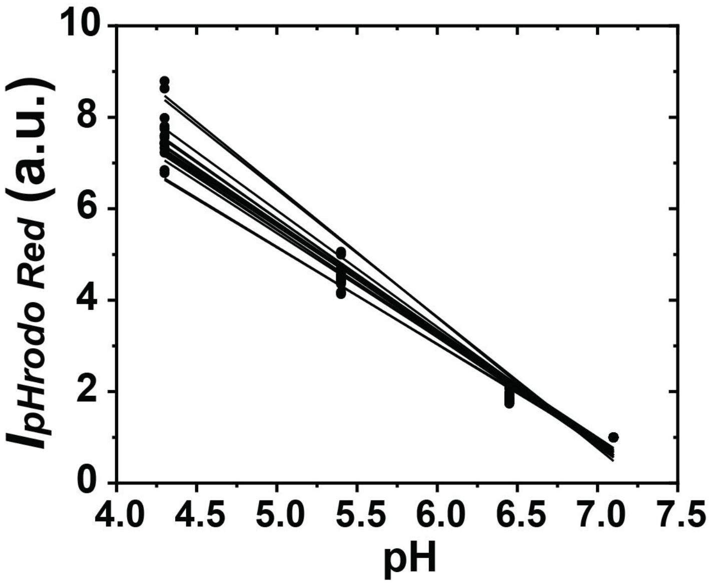


**Fig. S13.** Extracellular calibration curves for 15 mag-*Toxoplasma*. The pHrodo Red intensity was normalized to that at pH 7.1 at each sample. Black lines indicate linear fittings of the data. The average $R^2$ value of those fittings is 0.98.

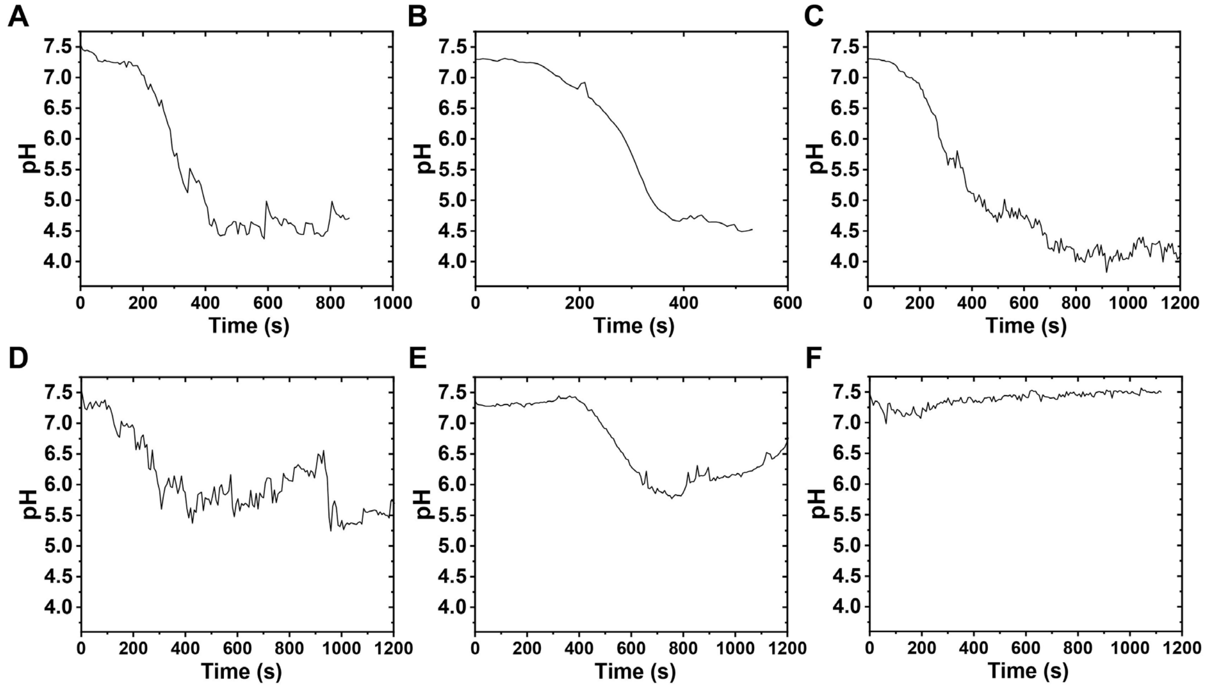


**Fig. S14.** Line plots showing the pH change of phagosomes containing mag-*Toxoplasma* as a function of time under conditions that no manipulation applied (*A*-*C*) and cell entry was magnetically manipulated (*D*-*F*).

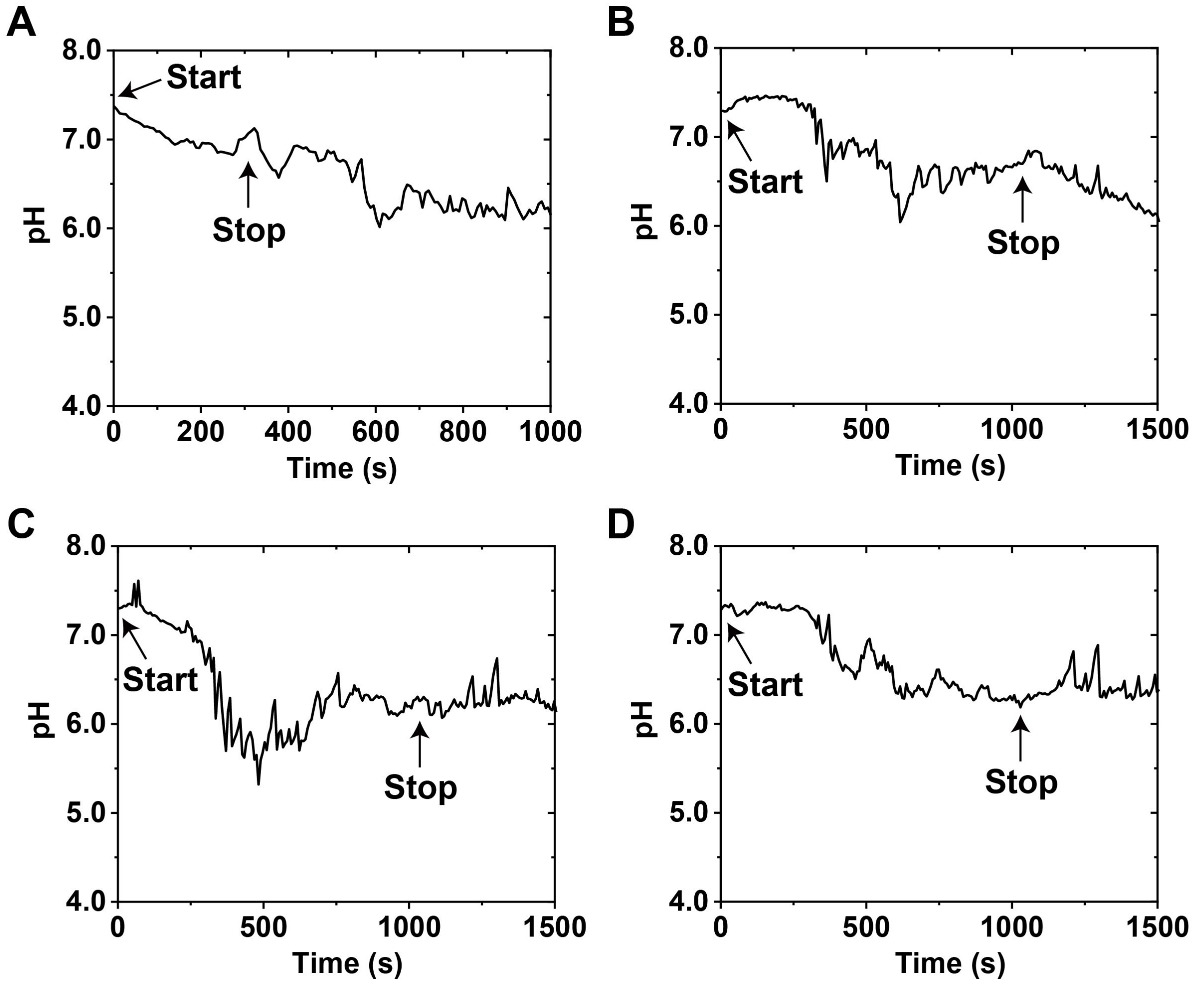


**Fig. S15.** Line plots showing the pH change of phagosomes containing mag-*Toxoplasma* as a function of time. Magnetic force was applied during the cell entry and was turned off after the formation of phagosome (on and off time points indicated in plots).

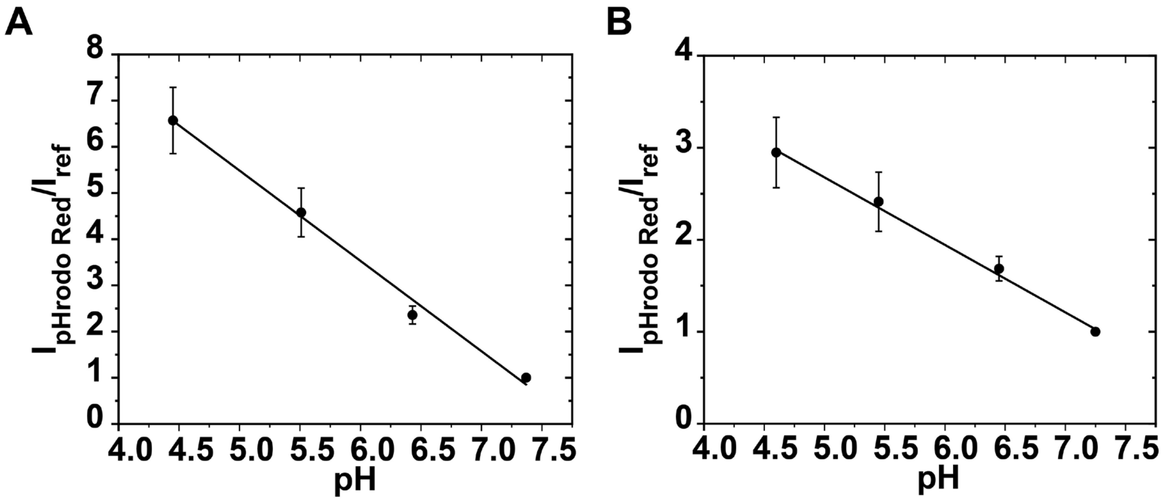


**Fig. S16.** Extracellular (*A*) and intracellular (*B*) pH calibration plots showing the fluorescence intensity change of the ratio of pHrodo Red and CF640R of individual MagSensors at different pH values. The plots are based on data from 18 and 9 particles in *A* and *B*, respectively. Error bars represents standard deviation.

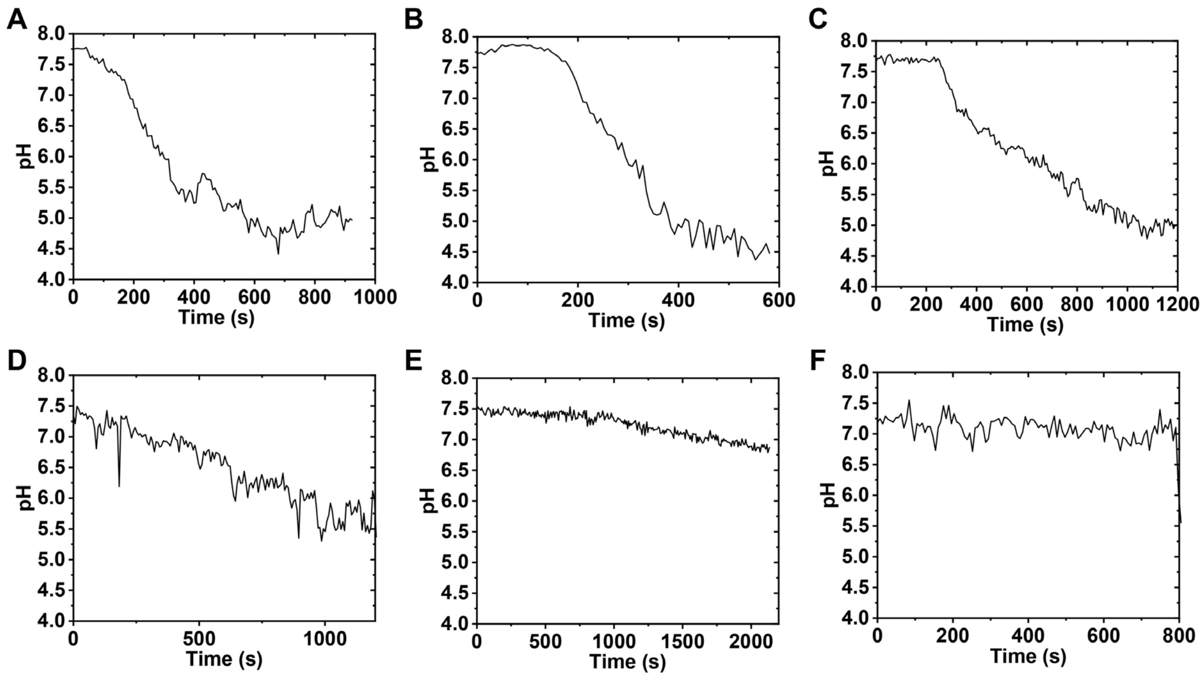


**Fig. S17.** Line plots showing the pH change of phagosomes containing pH-MagSensors as a function of time under conditions that no manipulation applied (*A*-*C*) and cell entry was magnetically manipulated (*D*-*F*).

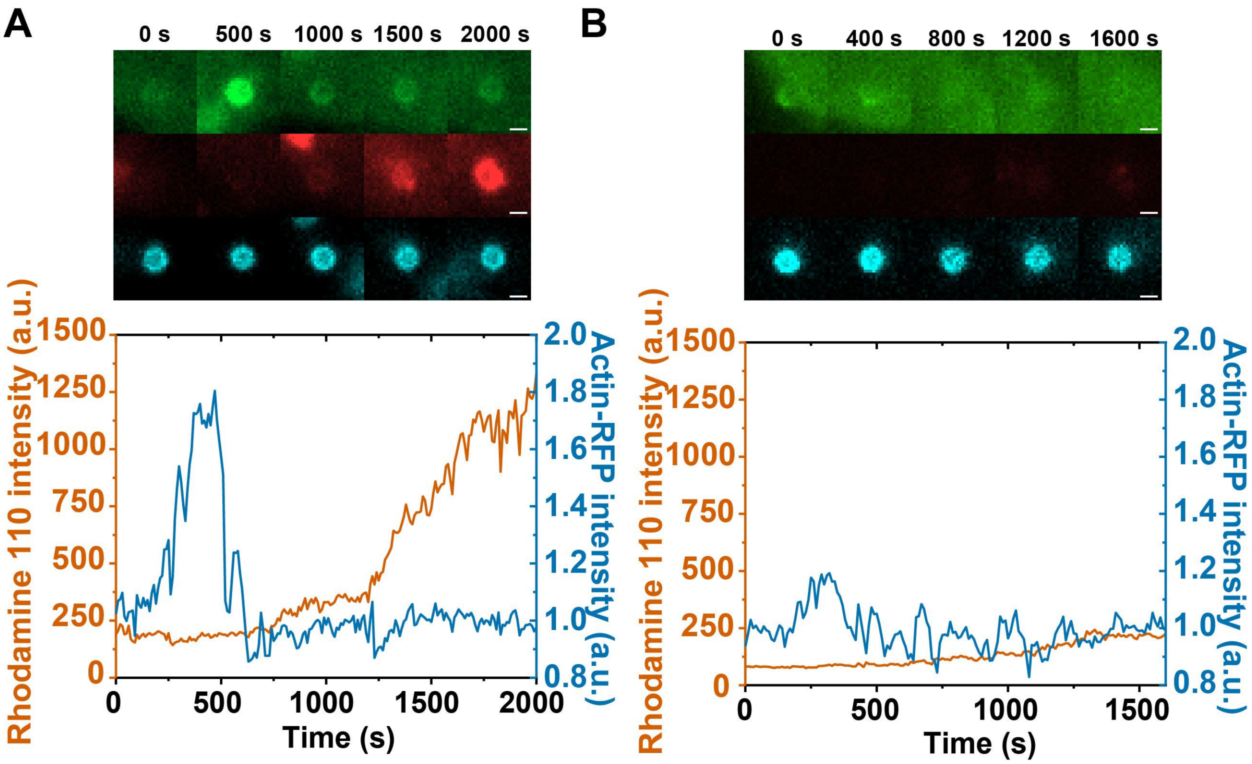


**Fig. S18.** Fluorescence images and line plots showing the measurement of proteolytic activity inside two maturing phagosomes with no force applied (*A*) and with magnetic pulling (*B*) during their cell entry. In fluorescence images, Actin-RFP was shown in green, ZFR-R110 was shown in red, and CF640R was shown in blue. Scale bars, 1 μm.

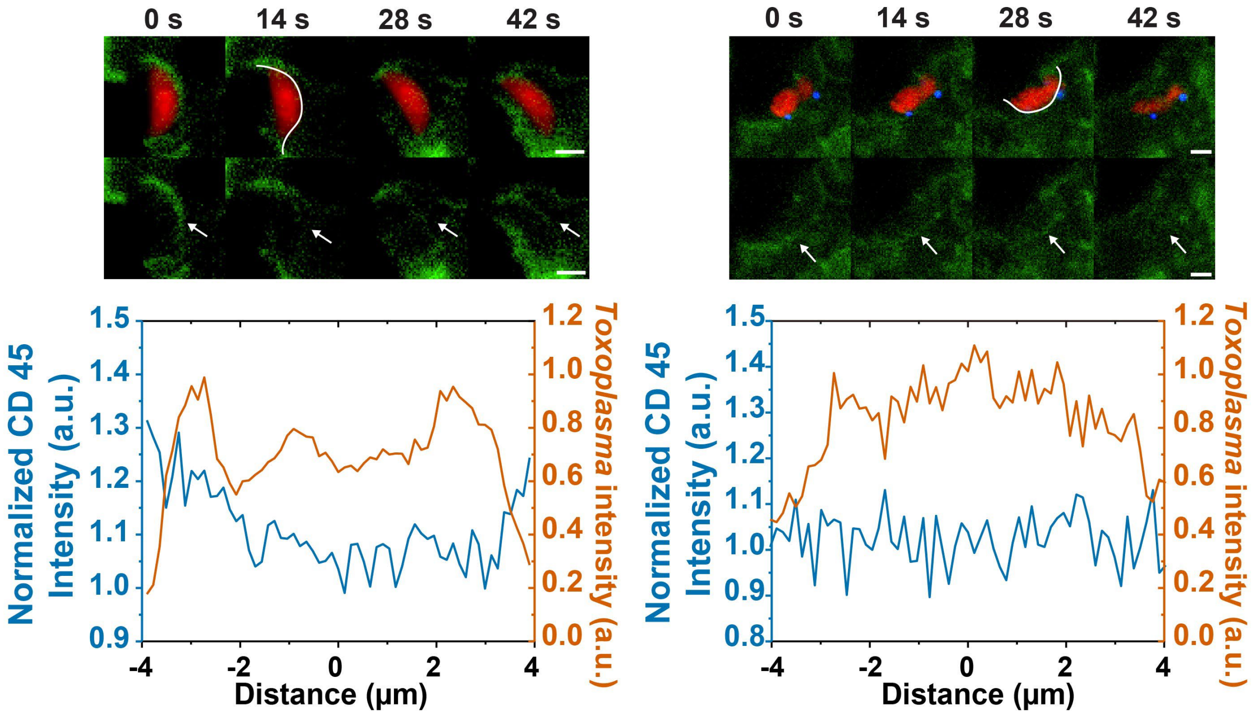


**Fig. S19.** Fluorescence images and line-scan intensity plots showing the CD45 (green) exclusion during the internalization of two sample heat-killed *Toxoplasma* (red) with no force applied (*A*) and with magnetic pulling (*B*), respectively. Magnetic nanoparticles are shown in blue. The arrows in the second row indicate the *Toxoplasma*-cell membrane binding area. Scale bars, 2 μm.

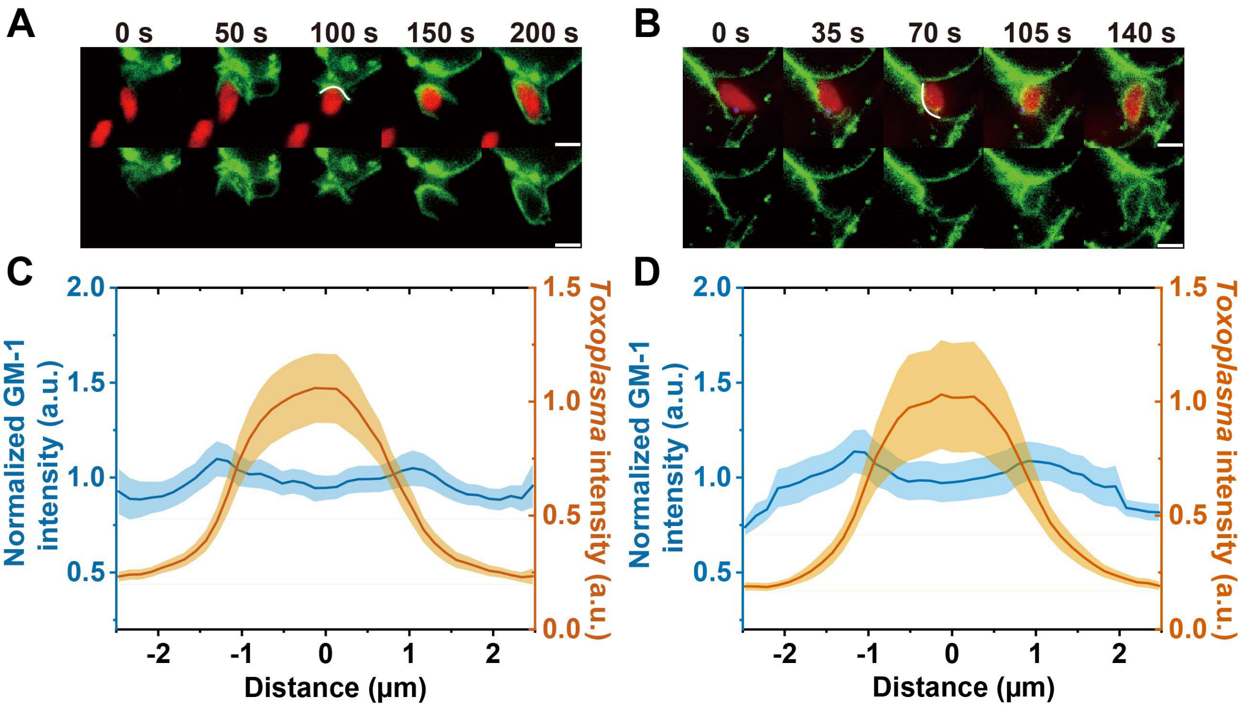


**Fig. S20.** (*A* and *B*) Fluorescence images showing the distribution of CTB labeled GM1 (green) in the phagocytic synapse formed between a heat-killed *Toxoplasma* (red) and a RAW264.7 cell, with no force applied (A, C) and with magnetic force (B, D). Scale bars, 2 µm. White lines indicate the contact surface between macrophages and parasite where line-scan fluorescence intensity was analyzed. (*C* and *D*) Line-scan intensity showing the fluorescence intensity of GM-1 and pHrodo Red labeled *Toxoplasma* at the parasite-macrophage contact site with no force applied (C)and with magnetic manipulation (D). Each line plot is an average from 10 individual cells. Shaded areas represent standard error of the mean.

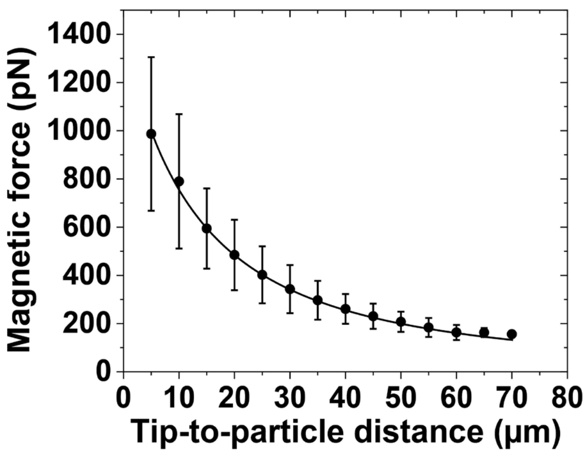

**Fig. S21.** Calibration plots showing the magnetic force exerted on single 2.8 μm magnetic beads plotted as a function of the distance from the bead to the tip of magnetic tweezers solenoid. Error bars are standard deviation from 12 beads.

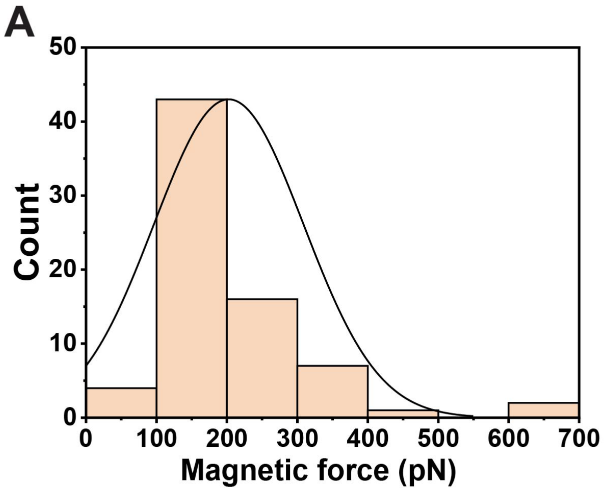


**Fig. S22.** (*A*) Distribution of the estimated magnetic force applied on individual 2.8 μm magnetic beads at the beginning of live cell imaging. (*B*) Distribution of the estimated magnetic pulling force exerted on individual 2.8 μm magnetic beads at the end of imaging. The average magnetic pulling force was 174 pN at the beginning of magnetic manipulation and 194 pN at the end of imaging as the bead moved closer to tip.

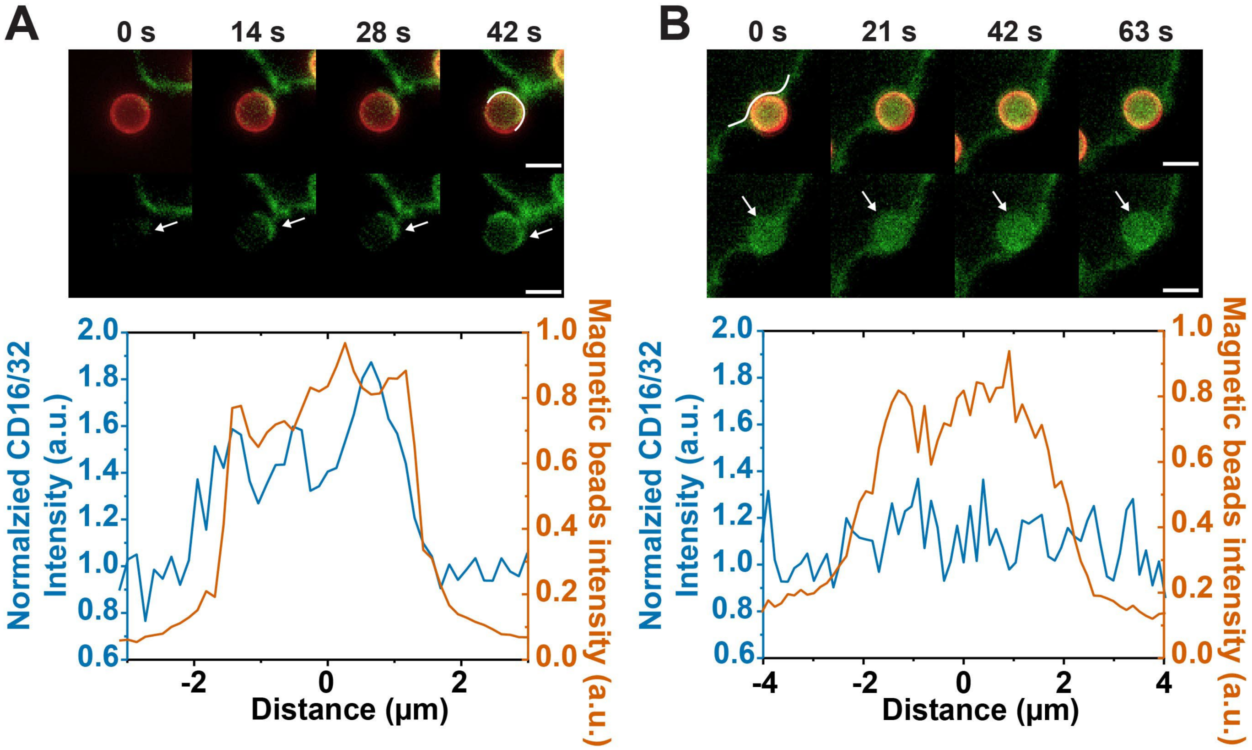


**Fig. S23.** Fluorescence images and line-scan intensity from single cells show clustering of FcγRs (CD16/32, shown in green) at the phagocytic synapse during the internalization of 2.8 μm magnetic beads (red), with no force (*A*) and with magnetic force (*B*), respectively. The line-scan plots of magnetic bead intensity are used to identify the beginning and end of the cell-bead contact area. White arrows in fluorescence images indicate the bead-cell membrane contact area. Scale bars, 3 μm.

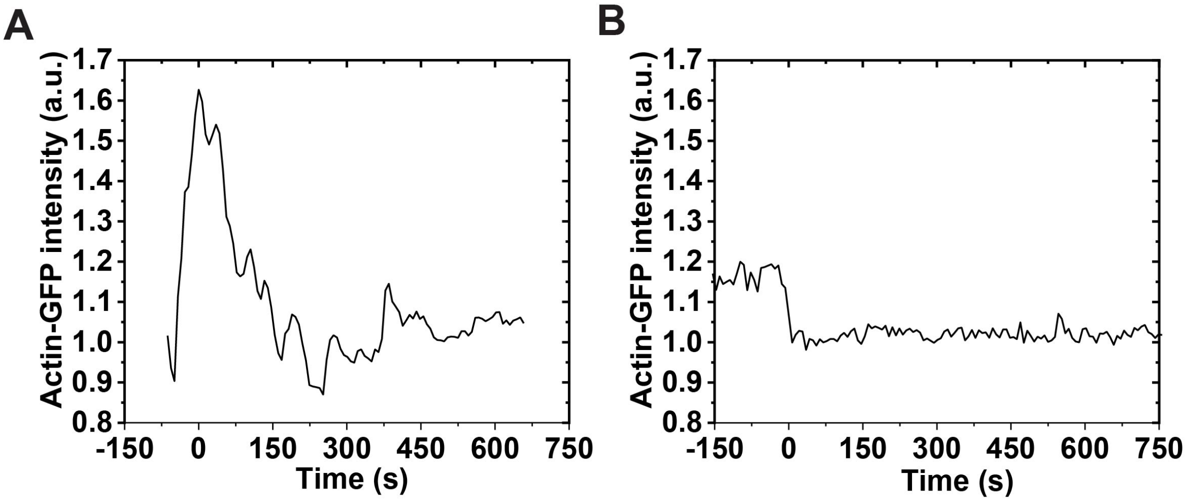


**Fig. S24.** Single cell data showing the actin-GFP intensity around nascent phagosome as a function of time with no magnetic force (*A*) and with magnetic pulling force applied (*B*). For consistent comparison between single cell data, time zero is defined as the point when actin intensity reaches the peak value.

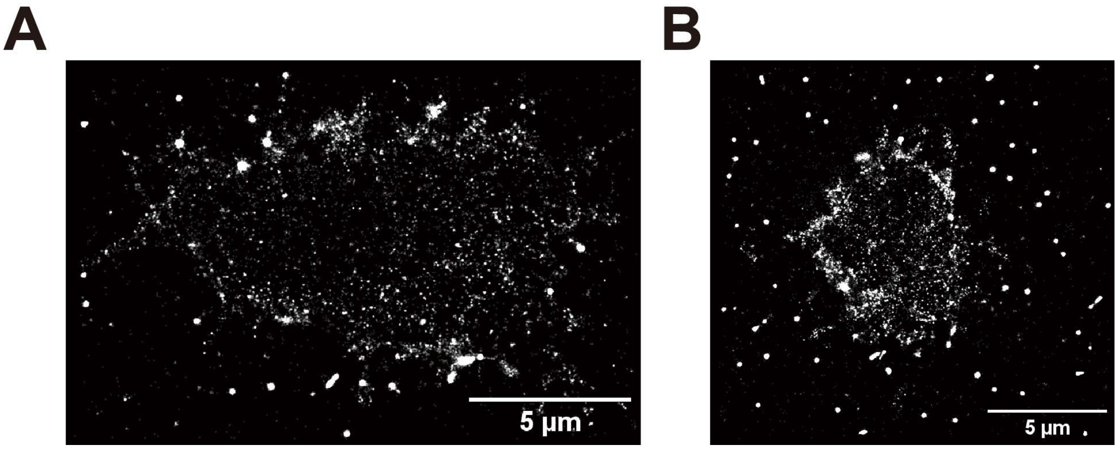


**Fig. S25.** TIRF-STORM images of immunostained CD45 in the plasma membrane of RAW264.7 cell.

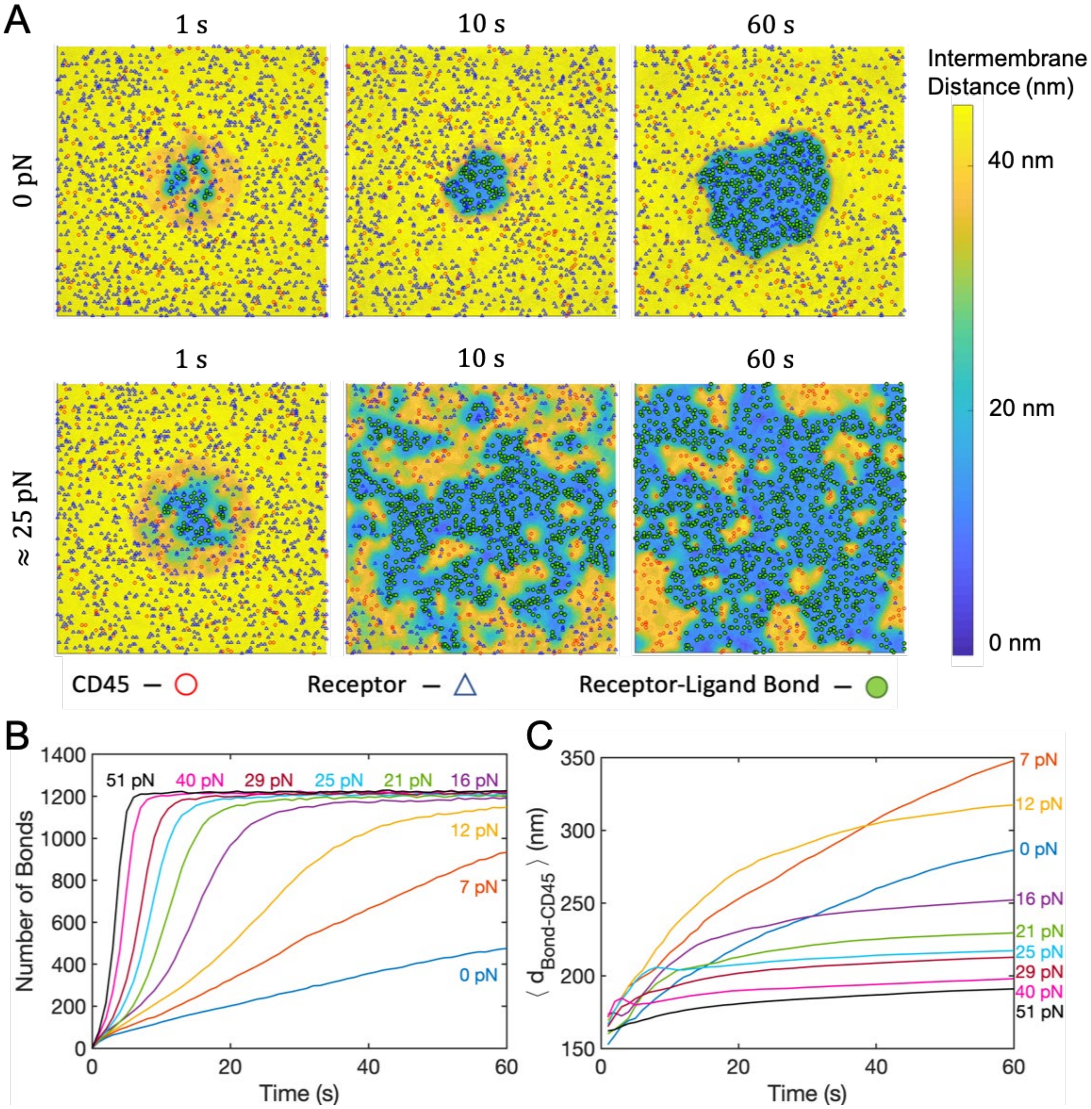


**Fig. S26.** Dynamics of early phagocytic synapse formation *in silico*. (A) Snapshots taken at three time points from simulation trajectories with 0 and ≈ 25 pN of applied force. Locations of free receptors (blue triangles), CD45 (red circles), and receptor-ligand bonds (filled green circles) are superimposed on a heat map representing the local distance between the two surfaces. Distances greater than 45 nm are shown in the same color (light yellow). Each frame represents a 2 $\mu$m × 2 $\mu$m patch of membrane. (B) Time dependence of receptor binding for different magnitudes of applied force, averaged over 10 independent trajectories. (C) The average distance from each receptor-ligand bond to the nearest CD45 molecule ($\langle d_{\mathrm{Bond-CD45}} \rangle$). The time-dependent average of 10 independent trajectories is shown for each applied force.

**Video S1.**

Representative simulation trajectories depicting the time evolution of the surface proteins and intermembrane distance. The applied force increases from left to right. Locations of free receptors (blue triangles), CD45 (red circles), and receptor-ligand bonds (filled green circles) are superimposed on a heat map representing the local distance between the two surfaces. Distances greater than 45 nm are shown in the same color (light yellow). Each frame represents a 2 $\mu m \times$ 2 $\mu m$ patch of membrane.

**Video S2.**

Representative simulation trajectories depicting the time evolution of the shape of the macrophage surface. The applied force increases from left to right. The surface is colored according to the local distance between the two surfaces. Distances greater than 45 nm are shown in the same color (light yellow). Each frame represents a 2 $\mu m \times$ 2 $\mu m$ patch of membrane.

**References for SI**


1. O. V. Vieira *et al.*, Modulation of Rab5 and Rab7 recruitment to phagosomes by phosphatidylinositol 3-kinase. *Mol Cell Biol* **23**, 2501-2514 (2003).
2. A. A. Minin *et al.*, Regulation of mitochondria distribution by RhoA and formins. *Journal of Cell Science* **119**, 659-670 (2006).
3. M. N. Teruel, T. A. Blanpied, K. Shen, G. J. Augustine, T. Meyer, A versatile microporation technique for the transfection of cultured CNS neurons. *J Neurosci Meth* **93**, 37-48 (1999).
4. W. L. Lee, D. Mason, A. D. Schreiber, S. Grinstein, Quantitative analysis of membrane remodeling at the phagocytic cup. *Mol Biol Cell* **18**, 2883-2892 (2007).
5. C. C. Scott *et al.*, Phosphatidylinositol-4,5-bisphosphate hydrolysis directs actin remodeling during phagocytosis. *J Cell Biol* **169**, 139-149 (2005).
6. D. S. Roos, R. G. K. Donald, N. S. Morrissette, A. L. C. Moulton, Molecular Tools for Genetic Dissection of the Protozoan Parasite Toxoplasma-Gondii. *Method Cell Biol* **45**, 27-63 (1994).
7. I. S. Behbahan, M. A. McBrian, S. K. Kurdistani, A protocol for measurement of intracellular pH. *Bio-protocol* **4**, e1027-e1027 (2014).
8. R. Parthasarathy, Rapid, accurate particle tracking by calculation of radial symmetry centers. *Nature Methods* **9**, 724-U291 (2012).
9. C. Zahn *et al.*, Measurement of the magnetic moment of single Magnetospirillum gryphiswaldense cells by magnetic tweezers. *Sci Rep* **7**, 3558 (2017).
10. F. Etoc *et al.*, Subcellular control of Rac-GTPase signalling by magnetogenetic manipulation inside living cells. *Nat Nanotechnol* **8**, 193-198 (2013).
11. A. M. Kaufmann, S. D. Goldman, J. P. Krise, A fluorescence resonance energy transfer-based approach for investigating late endosome-lysosome retrograde fusion events. *Anal Biochem* **386**, 91-97 (2009).
12. M. H. Bakalar *et al.*, Size-Dependent Segregation Controls Macrophage Phagocytosis of Antibody-Opsonized Targets. *Cell* **174**, 131-142 e113 (2018).
13. L. Sanchez, Y. Yi, Y. Yu, Effect of partial PEGylation on particle uptake by macrophages. *Nanoscale* **9**, 288-297 (2017).
14. M. Banerjee *et al.*, Cellubrevin/vesicle-associated membrane protein-3-mediated endocytosis and trafficking regulate platelet functions. *Blood* **130**, 2872-2883 (2017).
15. M. Howarth *et al.*, Monovalent, reduced-size quantum dots for imaging receptors on living cells. *Nat Methods* **5**, 397-399 (2008).
16. G. M. De Luca *et al.*, Re-scan confocal microscopy: scanning twice for better resolution. *Biomed Opt Express* **4**, 2644-2656 (2013).
17. M. Li *et al.*, Immobile ligands enhance FcgR-TLR2/1 crosstalk by promoting interface overlap of receptor clusters. *Biophysical Journal* **121**, 966-976 (2022).
18. M. D. Murtey, P. Ramasamy, Sample preparations for scanning electron microscopy–life sciences. *Modern electron microscopy in physical and life sciences*, 161-185 (2016).
19. M. Ovesny, P. Krizek, J. Borkovec, Z. K. Svindrych, G. M. Hagen, ThunderSTORM: a comprehensive ImageJ plug-in for PALM and STORM data analysis and super-resolution imaging. *Bioinformatics* **30**, 2389-2390 (2014).
20. F. Huang, S. L. Schwartz, J. M. Byars, K. A. Lidke, Simultaneous multiple-emitter fitting for single molecule super-resolution imaging. *Biomedical Optics Express* **2**, 1377-1393 (2011).
21. P. Krizek, I. Raska, G. M. Hagen, Minimizing detection errors in single molecule localization microscopy. *Optics Express* **19**, 3226-3235 (2011).
22. R. H. Pullen, S. M. Abel, Catch Bonds at T Cell Interfaces: Impact of Surface Reorganization and Membrane Fluctuations. *Biophys. J.* **113**, 120-131 (2017).
23. R. Grima, T. J. Newman, Accurate discretization of advection-diffusion equations. *Physical Review E* **70**, 036703 (2004).
24. D. T. Gillespie, Stochastic Simulation of Chemical Kinetics. *Annu. Rev. Phys. Chem.* **58**, 35-55 (2007).
25. S. Raychaudhuri, A. K. Chakraborty, M. Kardar, Effective Membrane Model of the Immunological Synapse. *Phys. Rev. Lett.* **91**, 208101 (2003).
